# Biomolecular design through latent reasoning with RFD4-Proteina

**DOI:** 10.64898/2026.11.12.766037

**Authors:** Nathaniel Corley, Kieran Didi, Maxwell W. Kazman, Rafael I. Brent, Magnus S. Bauer, Riti Biswas, Julia E. Bonzanini, Adam M. Chazin-Gray, Jie Chen, Andrew Favor, Maximilian Gantz, Henry A. Jones, Paul Kim, Margaret C. Lunn-Halbert, Nora K. McNamara-Bordewick, Saman Salike, DéJenaé L. See, Shajesh Sharma, Eric Sun, Preetham Venkatesh, Seth M. Woodbury, Kejia Wu, Yuyang Zhang, Joel D. Allen, Jasper Butcher, Lemuel Chang, Yehlin Cho, Alexis Courbet, Brian Coventry, Christian Dallago, Elliot S. Delfosse, Helen E. Eisenach, Mariana Garcia Sanchez, Tomas Geffner, Inna Goreshnik, Per Greisen, Andrew C. Hunt, David E. Kim, Donghyo Kim, Brianne R. King, Rohith Krishna, Yanjing Li, Zi Ming Li, Aiko Muraishi, Edin Muratspahíc, Oliver Nagl, Griffin H. Rangel, Antonia Ruder, Hojeong Shin, Zefei Xu, Yuwei Yang, Max Crispin, Neil P. King, Frank DiMaio, Gaurav Bhardwaj, Michael Bronstein, Karsten Kreis, Donald Hilvert, Anthony P. Green, Arash Vahdat, David Baker

**Affiliations:** Institute for Protein Design, University of Washington, Seattle, WA, USA; Department of Biochemistry, University of Washington, Seattle, WA, USA; Department of Bioengineering, University of Washington, Seattle, WA, USA; NVIDIA Corporation, Santa Clara, CA, USA; Department of Computer Science, University of Oxford, Oxford, UK; Graduate Program in Biological Physics, Structure and Design, University of Washington, Seattle, WA, USA; Graduate Program in Molecular Engineering, University of Washington, Seattle, WA, USA; MOE Key Laboratory of Bioinformatics, School of Life Sciences, Tsinghua University, Beijing, China; Beijing Frontier Research Center for Biological Structure, Tsinghua University, Beijing, China; Paul G. Allen School of Computer Science and Engineering, University of Washington, Seattle, WA, USA; Manchester Institute of Biotechnology, The University of Manchester, Manchester, UK; Department of Chemistry, The University of Manchester, Manchester, UK; School of Biological Sciences, University of Southampton, Southampton, UK; Department of Medicinal Chemistry, University of Washington, Seattle, WA, USA; Howard Hughes Medical Institute, University of Washington, Seattle, WA, USA; Department of Biosystems Science and Engineering, ETH Zurich, Basel, Switzerland; Department of Biostatistics and Bioinformatics, Duke University, Durham, NC, USA; BioMap Research, Palo Alto, CA, USA; Department of Chemical Engineering, University of Washington, Seattle, WA, USA; AITHYRA, Research Institute for Biomedical Artificial Intelligence, Austrian Academy of Sciences, Vienna, Austria; Laboratory of Organic Chemistry, ETH Zurich, Zurich, Switzerland; Institute for Innovations in Drug Disposition and Delivery, University of Washington, Seattle, WA, USA; Max Planck Institute for Terrestrial Microbiology, Marburg, Germany

## Abstract

Biomolecular design requires finding chemically and geometrically compatible arrangements of interacting atoms and molecules. The complexity of problems which can be tackled by deep learning design models is limited by both the molecular landscape they encompass and their ability to explore that landscape effectively. Here we introduce RFD4-Proteina, which addresses these limitations through intrinsic and extrinsic reasoning within a shared chemical and conformational latent space. The latent space encompasses standard and modified amino acids, sugars, DNA, and RNA, with force-field supervision favoring physically plausible atomic reconstructions. Within this space, intrinsic reasoning uses recurrent refinement to adaptively allocate computation, while extrinsic reasoning guides search with arbitrary constraints such as energetic or geometric criteria. Across *in silico* protein design benchmarks, RFD4-Proteina achieves higher design success rates at greater generation throughput than all previous methods evaluated, with further gains from conformational flexibility, recurrent refinement, and search. The same model performs equivalently to or better than state-of-the-art specialist methods for side-chain packing, sequence design, conformational sampling, and design refinement. We demonstrate the power and versatility of RFD4-Proteina experimentally by designing specific and cross-reactive binders to therapeutically relevant targets, recognizing phosphorylations, controlling glycosylation, and creating enzymes that catalyze both bond-breaking and bond-forming reactions, including a first proof-of-concept demonstration of a *de novo* designed enzyme for active pharmaceutical ingredient synthesis. By unifying diverse chemistry, conformational flexibility, and scalable reasoning in a single physically grounded latent space, RFD4-Proteina expands the frontier of biomolecular design.

## 1 Introduction

Biomolecular function arises from precise interactions between diverse chemical species. Recent advances in computational protein design have enabled the engineering of proteins with specified binding and catalytic activities [1–5]. Robust design of increasingly complex functions, however, requires accounting for the coupling between molecular identity, structural conformation, and interaction geometry. Current approaches address distinct aspects of this problem. RFdiffusion3 (RFD3) supports all-atom generation across diverse molecular contexts but does not design high-quality sequences for its generated structures, limiting simultaneous optimization of molecular composition and conformation [6]. Proteina-Complexa co-generates protein sequence and structure while broadening exploration through reward-guided search, but relies on a protein-specific representation with fixed-conformation interacting partners [7, 8]. Outstanding challenges include representing molecular composition and conformation across arbitrary biomolecular systems and efficiently navigating the resulting space to satisfy multiple interdependent constraints.

We set out to encode diverse chemical components, including standard and modified amino acids, sugars, DNA, and RNA, in a shared latent space and train a diffusion model to generate chemical identity and conformation jointly. Physics-based methods complement this generative model in three ways: they supervise training toward energetically favorable conformations, supply synthetic data to extend coverage of under-represented chemistries, and provide energy evaluations to steer generation. Because designs must satisfy complex chemical and geometric constraints, we add reasoning at two levels: intrinsic reasoning through recurrent refinement within the network, and extrinsic reasoning through guided search over generated candidates.

## 2 Biomolecular reasoning in a shared chemical and conformational space

We developed RFD4-Proteina to reason over molecular identities and conformations within a shared latent space (Fig. 1).

To construct a unified representation, we trained an autoencoder to encode each standard or modified amino acid, sugar, or nucleotide as an anchor-atom coordinate (e.g., alpha carbon for amino acids) and a 32-dimensional latent vector (Fig. 1a). Other molecular components retain explicit atom identities and connectivity. Distances in the learned latent space reflect chemical relationships: encodings of modified amino acids are closest to their unmodified parents in 44% of cases, while RNA nucleotides are closest to their corresponding DNA counterparts in 99% of cases (Fig. 1a, Supplementary Fig. 7). To favor physically plausible reconstructions, we incorporated a Rosetta-energy loss, differentiated through decoded coordinates using TMol, during fine-tuning [9, 10]. Energy-supervised fine-tuning lowered mean Rosetta energy by 1.07 kcal/mol per residue and reduced severe bond-angle deviations by 62% relative to reconstruction-only fine-tuning (Supplementary Fig. 8; Supplementary Section S6.1).

We next trained a flow-matching model to jointly generate molecular coordinates and latent vectors [11], extending the partially latent formulation introduced in La-Proteina and adopted by Proteina-Complexa [7, 12]. Compared to previous sequence-structure codesign methods, we enhanced the model along four key axes. First, we expanded the training data with physically simulated synthetic examples and structure-prediction distillation data across a range of modalities including small molecules, nucleic acids, and protein interfaces (Supplementary Section S1). When sampling conditioning information during training, we tuned features to match realworld design problems and promote the model’s understanding of practically useful constraints. For example, we sampled tip-atom positional constraints in pockets resembling enzyme active sites, leveraging abundant protein-only data to yield realistic atomic motif scaffolding tasks (Supplementary Table S4). Second, we strengthened the pair representation through triangle multiplicative updates every other transformer block, more expressive communication from single to pair features, and a wider hidden dimension (Fig. 1b). Third, we enabled intrinsic reasoning within trajectories by performing recycling at each diffusion time step rather than as an upstream feature-processing stage [6, 13–15]. The number of recurrent passes can vary across timesteps, allowing greater refinement at critical stages of the trajectory (Supplementary Section S3). The use of recycling at each timestep parallels concurrent developments in flow-based reasoning models [16–18]. Finally, we enabled extrinsic reasoning through search across trajectories, shifting the sampling distribution towards designs that satisfy arbitrary user-provided metrics. Extrinsic reasoning can be performed in a feature- and timestep-dependent manner, optimizing for each metric precisely when the corresponding molecular features become resolved.

**Fig. 1.**
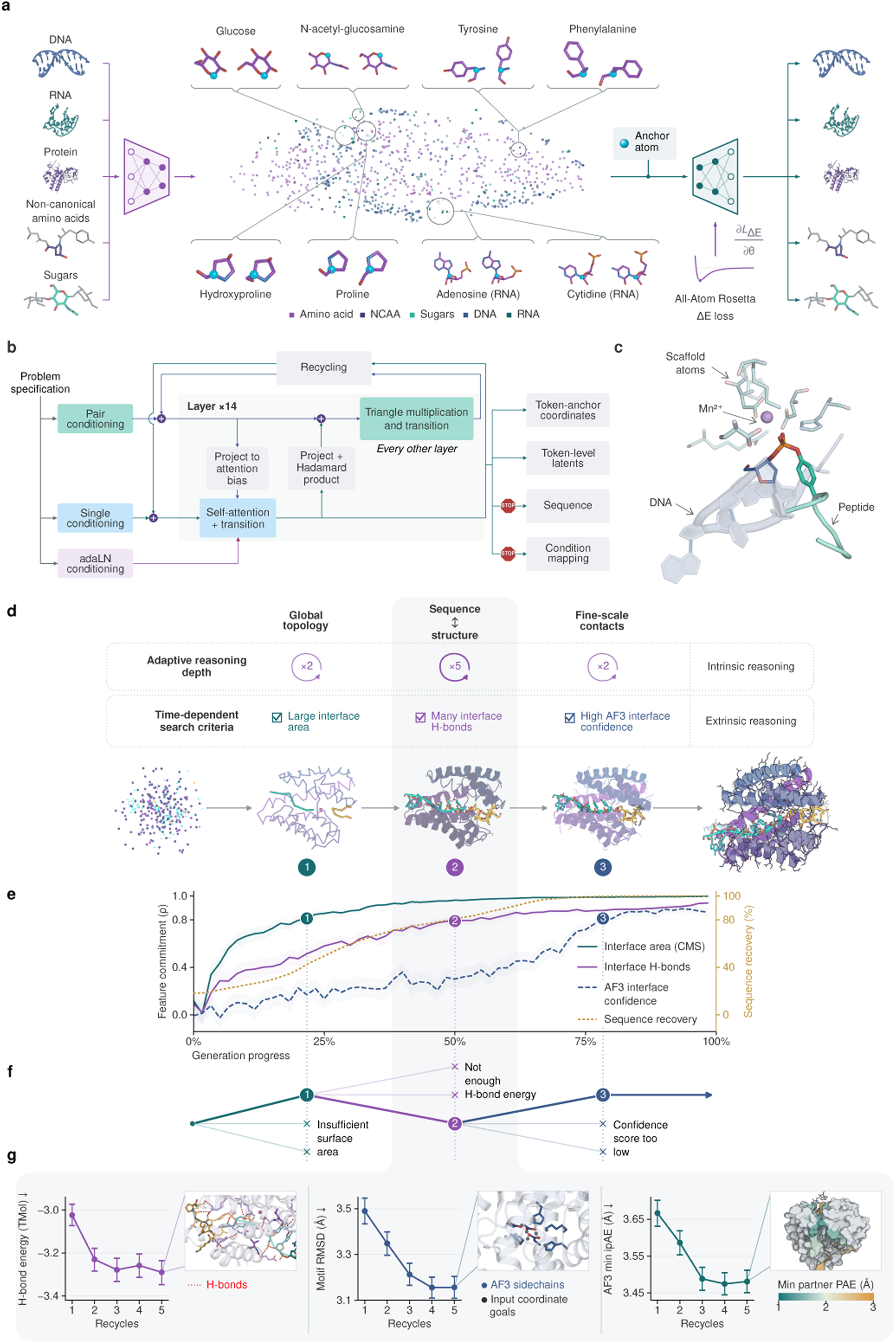
Biomolecular reasoning in a shared chemical and conformational space. **a,** An autoencoder represents standard and modified amino acids, sugars, DNA, and RNA using anchor-atom coordinates and 32-dimensional latent vectors. For the t-SNE projection, residues from PDB validation structures spanning these molecular classes were encoded. Posterior means were projected into two dimensions and colored by token type. A differentiable Rosetta-energy loss supervises atomic reconstruction. **b,** The problem specification is featurized into single, pair, and adaptive layer-normalization (adaLN) conditioning for a denoising network with recurrent updates. The network predicts velocities for latent vectors and anchor-atom coordinates; auxiliary heads predict insertion positions of unindexed residues and the resolved sequence, which conditions the decoder. STOP symbols denote detached gradients (Supplementary Section S3). **c–g,** One illustrative enzyme-design task demonstrates intrinsic reasoning and temporal search. **c,** Input constraints comprise scaffold atoms, a manganese ion, and a flexible topoisomerase-derived peptide linked to DNA through a 3’-phosphotyrosyl bond. **d,** Generation trajectory showing adaptive recurrent refinement and stage-specific search criteria. Numbers inside the circular arrows indicate the number of recurrent passes at each step under the adaptive recycling schedule. **e,** Feature commitment across 500 trajectories: Spearman correlations (*ρ*) compare interface area (contact molecular surface), hydrogen bonding, and AlphaFold 3 interface confidences between intermediate rollouts and the final structure. Numbered search checkpoints mark the first crossings of *ρ* = 0.8. Sequence recovery measures agreement with the final sequence. **f,** Temporal search branches trajectories and terminates those that fail stage-specific continuation criteria (Supplementary Section S7.1.1). **g,** Effect of one to five recurrent passes per timestep at steps 20–30 of a 60-step trajectory, evaluated on AF3 refolds of the final sequences. Plots show TMol-calculated [10] interface hydrogen-bond energy, active-site motif RMSD, and minimum interface predicted aligned error (min-iPAE).

It is instructive to examine the model’s generation process on a complex test case: designing an enzyme to cleave the covalent bond between a peptide and DNA in a flexible substrate (Fig. 1c). This task combines prescribed tip-atom coordinates, a modified active-site amino acid, a manganese ion, and the nonstandard protein-DNA linkage. We evaluated designs for agreement with atomic constraints, interface complementarity, hydrogen bonding, and structure-prediction confidence (Supplementary Table S21). We observed that design objectives become resolved at distinct time points during generation (Fig. 1d,e). Across 500 trajectories, we launched independent rollouts and compared their properties with final outcomes. Interface area became predictable early, followed by hydrogen bonding, and finally AlphaFold 3 (AF3) interface confidence [19]; correlations with final outcomes reached *ρ* 0.8 at approximately 20%, 50%, and 80% of the way through generation, respectively. Meanwhile, sequence recovery rose from 50% to 97% as generation progressed from one-quarter to two-thirds completion. The model therefore continued selecting residue identities while refining local interface interactions, consistent with sequence–structure reasoning during the middle portion of generation.

Our extrinsic reasoning approach makes use of these differing convergence rates across distinct molecular properties. In this example, we select trajectories during inference-time search based successively on interface area, hydrogen bonding, and AF3 interface confidence (Fig. 1f). At each checkpoint, we discontinue trajectories that depart from the current criterion as they are unlikely to be rescued by subsequent steps. Such a temporal search efficiently navigates the solution space to find outputs that fulfill all criteria. Compared to näıve sampling, temporal search for our example bond-cleaving enzyme yielded three times as many unique passing designs per GPU-hour (Supplementary Table S18).

RFD4-Proteina performs intrinsic reasoning within each denoising step through repeated internal updates that refine its prediction. Increasing refinement from one to five internal updates at intermediate denoising steps produced designs whose AF3-predicted structures had more favorable interface hydrogen-bond energy (as calculated online by TMol [10]), more closely matched the prescribed active-site geometry, and exhibited higher interface confidence (Fig. 1g). These improvements in AF3-refolded structures suggest that recurrent refinement within this intermediate step window increases compatibility between the designed sequence and intended geometry. The effects depended on when refinement was applied: early refinement improved activesite geometry, whereas intermediate refinement also improved interface hydrogen-bond energy and AF3 interface confidence (Supplementary Fig. 10). We observed that the optimal reasoning schedule varies between modalities and tasks (Supplementary Fig. 11). In practice, we find that small-scale pilots are helpful in determining the best reasoning configurations for a given design challenge.

## 3 State-of-the-art computational design across molecular contexts

We compared RFD4-Proteina *in silico* with generative, hallucination, and refinement methods across protein-binding, small-molecule-binding, enzyme-scaffolding, and DNA-binding benchmarks [20]. To evaluate designs, we used AF3 to independently predict structures from the designed sequences, then assessed their agreement with the intended structures and their adherence to task-specific structural and confidence criteria. We measured both generation throughput and clustered pass rate, defined as distinct passing structural clusters per original attempt (Supplementary Section S8).

Across all four benchmarks, RFD4-Proteina solved more design challenges than any individual baseline while achieving higher clustered pass rates at greater generation throughput, without structure-predictor feedback during generation (Fig. 2). For protein binding, high-thinking RFD4-Proteina achieves 10.6% clustered pass rate versus 5.9% for the next-best model, BindCraft, while attaining 42-fold higher throughput [20]. On an updated Atomic Motif Enzyme (AME) benchmark, RFD4-Proteina achieves 12.1% clustered pass rate versus 5.7% for AlphaProtein-novo codesign, with 5.9-fold higher throughput. For small-molecule binding, RFD4-Proteina achieves a 6.1-fold higher clustered pass rate than the next-best model, RFD3 (14.3% versus 2.4%), with 3.3-fold higher generation throughput and passing designs for all 77 targets versus 64 [6]. On the three-target DNA benchmark from the RFdiffusion3 preprint [21], RFD4-Proteina achieves 3.5% clustered pass rate versus 0.8% for the next-best baseline, BoltzGen, and 0.5% for fixed-DNA RFD3. Including AF3 evaluation time, high-thinking RFD4-Proteina yielded 5.0-, 10.4-, 6.0-, and 5.5-fold more passing clusters per GPU-hour than the next-best baseline for protein binding, small-molecule binding, AME, and DNA binding, respectively (Fig. 2; Supplementary Section S8).

**Fig. 2.**
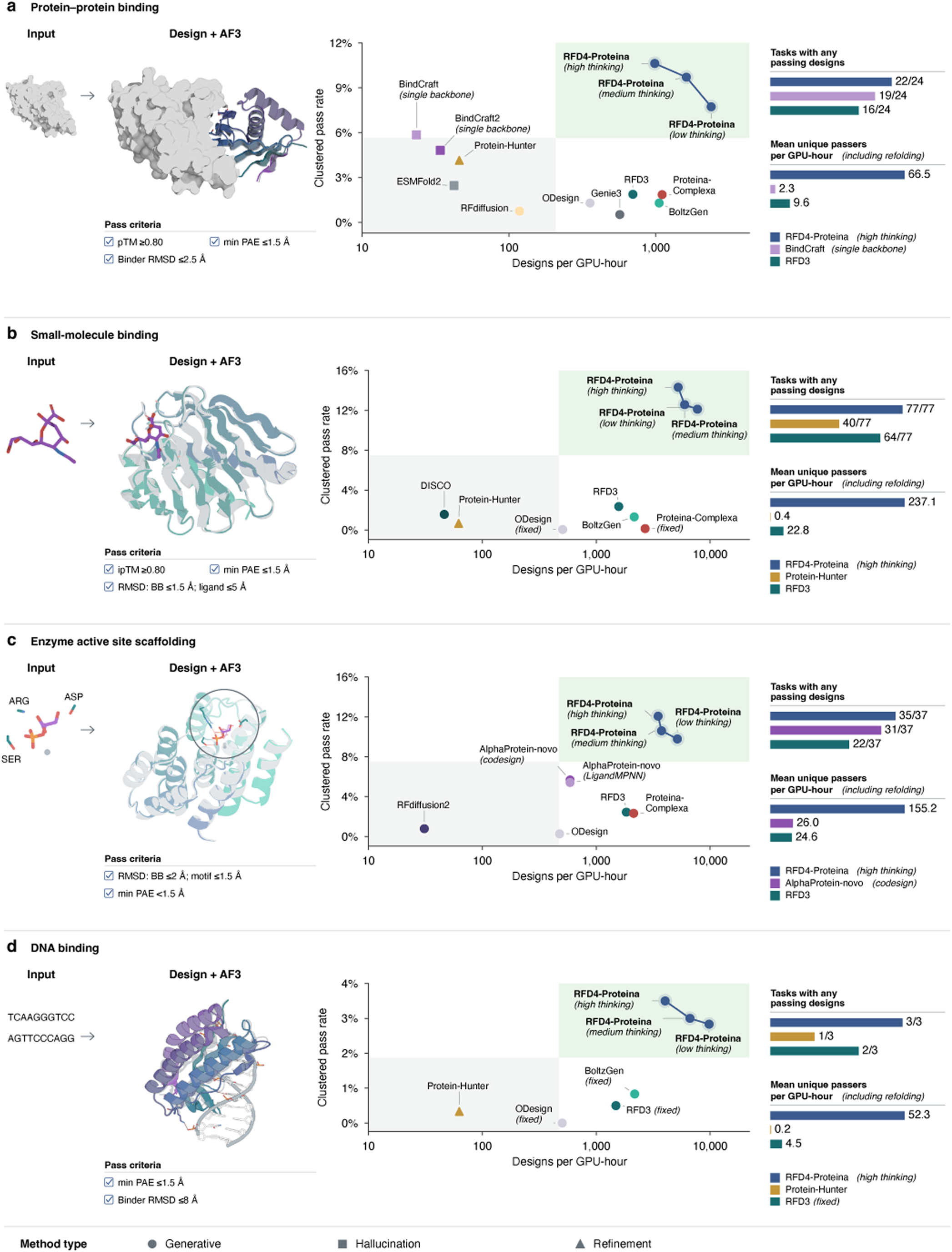
State-of-the-art computational design performance and efficiency across molecular contexts. **a–d,** Protein-binder design (union of Proteina-Complexa and RFdiffusion3 benchmarks, 24 targets) [6, 7], small-molecule binder design (STUDIO-170 subset from DISCO, 77 targets) [22], Atomic Motif Enzyme (AME) scaffolding (37 targets) [2], and DNA-binder design (three targets from the RFdiffusion3 preprint) [21]. Molecular views show inputs and design/AlphaFold3 (AF3) overlays. Scatter plots compare clustered pass rate with generation throughput (designs per GPU-hour). Clustered pass rate is distinct passing structural clusters divided by 200 original attempts per target, including failures, averaged across targets; passing structures are clustered using Foldseek (TM-score threshold 0.5). Right-hand plots show target coverage (at least one passing design) and mean target-level passing clusters per GPU-hour, including downstream AF3 evaluation. Each original attempt contributes at most one backbone–sequence pair to final AF3 evaluation. For methods that generate and filter multiple candidates internally, only the pipeline-selected pair is evaluated. Full-pipeline protein-binding comparisons are provided in Supplementary Fig. 16. Small-molecule and DNA conformations are generated unless labeled “fixed”; AME ligand coordinates are fully fixed. Protein-binding protocols condition on target backbone coordinates alone or include target side-chain coordinates. For methods supporting both modes, we show the mode with the higher benchmark-wide clustered pass rate. Low, medium, and high thinking denote increasing, task-specific recurrent-refinement budgets without structure-predictor feedback. Generation throughput includes intrinsic sequence design and model-specific internal prediction but excludes downstream AF3 evaluation. Task-specific success criteria, conditioning configurations, and timing procedures are detailed in Supplementary Section S8.

We next tested whether increasing intrinsic reasoning improved design success rates. Low-, medium-, and high-thinking settings allocate increasing computational budgets to recurrent refinement during generation (Fig. 1c–e), using task-specific schedules (Supplementary Section S8). Across these settings, clustered pass rates rose from 7.7% to 9.7% to 10.6% for protein binding, from 12.1% to 12.6% to 14.3% for small-molecule binding, from 9.8% to 10.6% to 12.1% for AME, and from 2.8% to 3.0% to 3.5% for DNA binding (Fig. 2). Generation throughput decreased with increasing computation: for protein binders, for example, designs per GPU-hour decreased from 2,388 with low-thinking to 986 with high-thinking. Additional recurrent refinement therefore increased the yield of distinct passing designs per attempt at the cost of generation throughput.

We also examined whether allowing the molecular context to change conformation during generation improved design success. With high-thinking RFD4-Proteina, cogenerating the ligand conformation with the designed protein rather than conditioning on a fixed conformer increased the clustered pass rate from 8.1% to 14.3% (Supplementary Fig. 12). Allowing target side chains and DNA conformations to change was likewise associated with higher clustered pass rates for protein binding (9.6% to 10.6%) and DNA binding (2.0% to 3.5%), respectively. Conditioning only on the target’s internal coordinates, without specifying its global frame, further increased the protein-binding clustered pass rate from 10.6% to 11.8%; we report this distinct task separately from Figure 2 (Supplementary Fig. 13; Supplementary Section S8.4.1).

We next asked whether extrinsic reasoning could improve design success by incorporating readouts from additional tools. Search can improve design efficiency by using external tools to avoid spending compute on unpromising trajectories (Fig. 1f). We see that on harder tasks, where there is the most to gain from saving compute, search can significantly increase sampling efficiency. To make the benchmark tasks more representative of typical design campaigns, we also evaluated search on six enzyme-scaffolding targets with an introduced ligand-burial requirement and on small-molecule design tasks with an additional molecular-interaction requirement. On these harder tasks, search allows the model to cancel unpromising trajectories as soon as these features are resolved, improving the efficiency of generating unique passing designs by 22.6% and 24.6% for the enzyme and small molecule tasks, respectively (Supplementary Sections S7.1.2 and S7.1.3). On easier tasks or tasks where all objectives are resolved late in the trajectory, we found that the efficiency gains from search do not outweigh the computational overhead (Supplementary Section S8.5).

To better understand these gains in performance, we investigated which aspects of the model’s architecture and training contribute to design success. We trained ablation models at reduced scale and compared them at matched training steps (Supplementary Section S3.1.6). First, we ablated triangle multiplication and found that these operations primarily inform global structural organization, reducing the token anchor placement loss by 9.7% while leaving the latent loss nearly unchanged (0.36% reduction). This improvement in topology bolsters agreement with AF3 refolding, including the backbone RMSD, minimum ipAE, and clustered pass rates across all benchmarks in Fig. 2. We then compared the Hadamard product operation used for the single-to-pair update against the elementwise sum used in prior work [12, 23], finding improvements on nearly all of the core refolding metrics and pass rates. The Hadamard product yielded a similar training loss at convergence but a lower loss early in training, suggesting that its advantage may stem from improved training dynamics. Next, we compared the intrinsic reasoning used in RFD4-Proteina against the self-conditioning used in prior work [1, 7, 12, 23]. We found that intrinsic reasoning provided a clear advantage, improving pass rates and minimum ipAE on all benchmarks while improving backbone RMSD on all except DNA binding. Finally, we investigated the effect of explorative modeling [24], a method in which each training example is run *K* times with independent noise samples at the same sampled time, and only the best output is used to backpropagate gradients. In RFD4-Proteina, we set *K* = 4 and backpropagate through the candidate with the lowest combined coordinate and latent flow-matching loss, observing improvements across all tasks except the AME benchmark. Beyond these architecture and training protocols, featurization changes also significantly improved geometry and stereochemical adherence, as detailed in Supplementary Sections S2.1.3 and S2.3.3.

## 4 Extending the generative framework to specialized molecular workflows

The preceding benchmarks test the generation of new designs. We next asked whether the same model could support additional molecular workflows used to build, evaluate, and refine designs, many of which are currently addressed by specialized models. We examined six use-cases spanning sequence assignment, conformational sampling, and refinement of existing structures (Fig. 3).

**Fig. 3.**
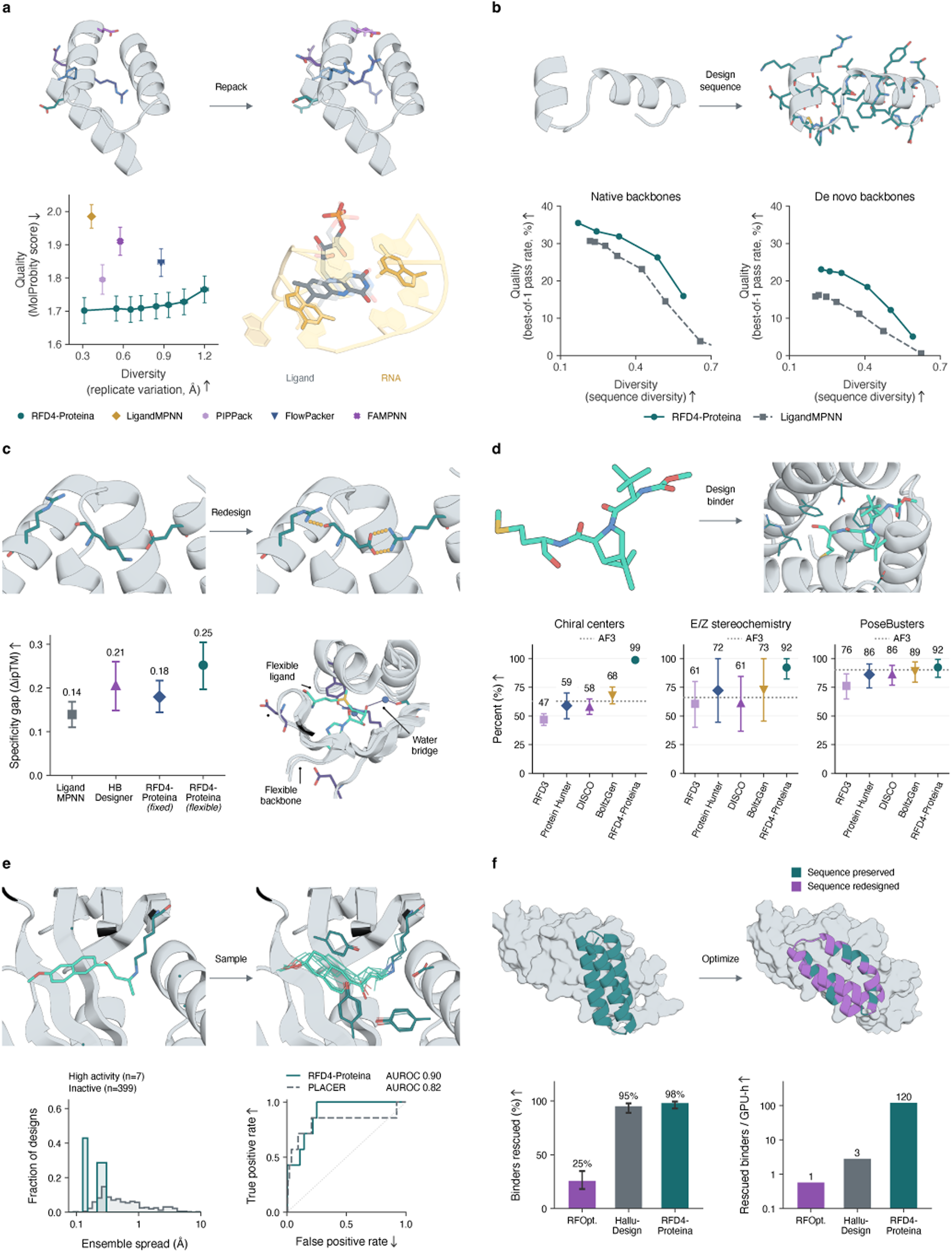
RFD4-Proteina outperforms specialized models across molecular tasks. **a,** Side-chain packing quality (MolProbity score; lower is better) versus diversity (pairwise RMSD) on 114 PackBench targets [25]. RFD4-Proteina is shown at eight sampling-noise settings across the quality-diversity trade-off, establishing a new Pareto frontier. The inset shows joint packing of RNA bases and a flexible ligand in an aptamer complex. **b,** Single-sequence AF3 refolding pass rate versus sequence diversity (one minus mean pairwise sequence identity). Left, 156 native backbones shared by both methods, with passing defined as backbone RMSD *≤* 1.5 Å. Right, 340 RFdiffusion3 enzyme back-bones involving fixed-sequence motifs, with passing defined as backbone RMSD *≤* 2.0 Å and motif all-atom RMSD *≤* 1.5 Å. At matched diversity RFD4-Proteina achieves a higher pass rate than LigandMPNN on both sets. **c,** Specificity gap within HBDesigner families, measured as difference in AF3 ipTM, upon flexible-backbone sequence redesign. The lower-right illustration shows a sampled SAM pocket with a ligand, backbone movement, conditioned motif sites and predicted water bridges. **d,** Adherence to chirality, E/Z double-bond stereochemistry, and PoseBusters pass rates across 43 ligands that do not appear in the training data for any of the evaluated models. Dotted lines show AF3 results when refolding RFD4-Proteina designs. RFD4-Proteina has higher stereochemical adherence and PoseBusters pass rates than the other methods shown. **e,** Retroaldolase ensembles and retrospective activity ranking [26]: seven highly active and 399 inactive designs, with 16 conformers per design. Smaller RFD4-Proteina ensemble spread distinguished highly active from inactive designs, with a higher observed AUROC than PLACER for the intermediate shown. **f,** Computational rescue of 102 binders from the RFD1 paper that fail *in silico* filters (six attempts each). Rescue is plotted as a percentage of the 102 original binders; throughput is rescued binders per GPU-hour. Partial diffusion with RFD4-Proteina rescues more designs than other methods in a fraction of the time.

### Side-chain packing

We first evaluated side chain packing when given the protein sequence and backbone coordinates. We followed the PackBench benchmark [25] and used the native backbones of 114 CASP14 and CASP15 targets with valid outputs from every model. RFD4-Proteina achieved a lower mean MolProbity score than LigandMPNN at similar conformational diversity (1.70 versus 1.99; lower is better; Fig. 3a) [27]. Greater sampling noise broadened mean pairwise side-chain variation from 0.31 to 1.20 Å, while the mean MolProbity score rose only from 1.70 to 1.77, indicating that RFD4-Proteina generates diverse conformations with-out compromising structural quality. Unlike other models, RFD4-Proteina side chain packing extends beyond proteins. We demonstrate this capability by jointly sampling RNA base rotamers and small-molecule conformations within an RNA aptamer–flavin mononucleotide (FMN) complex (Fig. 3a).

### Fixed-backbone sequence design

We next evaluated RFD4-Proteina as an inverse folding tool by resampling side-chain identities while keeping the backbone fixed. For each backbone, we generated eight sequences each with RFD4-Proteina and LigandMPNN and evaluated them by MSA-free AF3 refolding, adjusting the sampling parameters of each model to modulate the quality–diversity trade-off. We evaluated the models on a set of native backbones held out from training and a set of *de novo* backbones generated with an orthogonal method (RFD3). In both cases, RFD4-Proteina consistently achieved higher pass rates than LigandMPNN at matched diversity (Fig. 3b), establishing a new Pareto frontier for inverse folding.

### Flexible-backbone sequence design

We then relaxed the requirement of keeping the backbone coordinates fixed, allowing the model to refine local geometry during generation. In addition, we leveraged search with TMol [10] to optimize hydrogen bond energies at the interfaces of protein dimers. Making additional hydrogen bonds can improve target specificity, approximated *in silico* using AF3 ΔipTM between on- and off-targets, by introducing more precise contacts at interfaces (for example, see [28]). Across 13 HBDesigner families of on/off-targets, flexible-backbone RFD4-Proteina achieved a higher mean predicted specificity gap than the fixed-backbone protocol (0.25 versus 0.18 ΔipTM) and the highest mean among all methods evaluated (Fig. 3c). RFD4-Proteina is also more versatile than previous methods, as it can incorporate non-protein molecular context, support arbitrary combinations of fixed and flexible coordinates, and model physical properties like putative water bridges using TMol.

### Ligand stereochemistry

We assessed the ability of RFD4-Proteina to respect input ligand chemistry by designing binders against 43 CCD ligands absent from every evaluated model’s training data (Fig. 3d). We then used AF3 to refold these output complexes from RFD4-Proteina. We found that RFD4-Proteina outperforms both AF3 and the next-best generative method on adherence to chirality (98.8%, 62.9%, 67.9%), double-bond stereochemistry (92.2%, 66.1%, 72.8%), and physical plausibility measured by PoseBusters (92.3%, 90.2%, 89.1%) [29]. RFD4-Proteina maintained high stereochemical adherence on ligands absent from its training data (Supplementary Table S3).

### Ensemble-based filtering

In addition to conformational reasoning during *de novo* design, RFD4-Proteina also exhibits strong signal when used for retrospective campaign analysis. Following PLACER, we sample local conformational ensembles for designed enzyme active sites, using the consistency of independent outputs as a proxy for catalytic motif preorganization (Fig. 3e) [26]. We applied this procedure to the retrospective retroaldolase analysis used in PLACER, the current state-of-the-art. Although we did not optimize the model for this task, we found that the conformational spread of RFD4-Proteina outputs distinguished active designs as effectively as PLACER (Fig. 3e, Supplementary Fig. 22).

### Design refinement

Last, we asked whether existing designs could provide useful starting points even if they had failed experimental testing. We partially diffused published RFdiffusion binder designs that did not pass more modern AF3 structural filters, and assessed the ability of various models to rescue these designs *in silico*. With six attempts per design, RFD4-Proteina rescued 100 of 102 designs, compared to 97 with HalluDesign and 26 with RFOptimization, while doing so in a fraction of the time (Fig. 3f) [30, 31].

## 5 Experimental validation

For protein production and characterization at the Institute for Protein Design, we used established workflows [32].

### 5.1 Designing specific protein binders

There have been significant advances in protein binder design in recent years [1, 7, 13, 20, 33–39]. However, controlling specificity remains a challenge, whether the goal is cross-reactivity between species or selectivity among closely related off-targets. We therefore evaluated the ability of RFD4-Proteina to design for a variety of tasks in which specificity was a key constraint (Fig. 4).

**Fig. 4.**
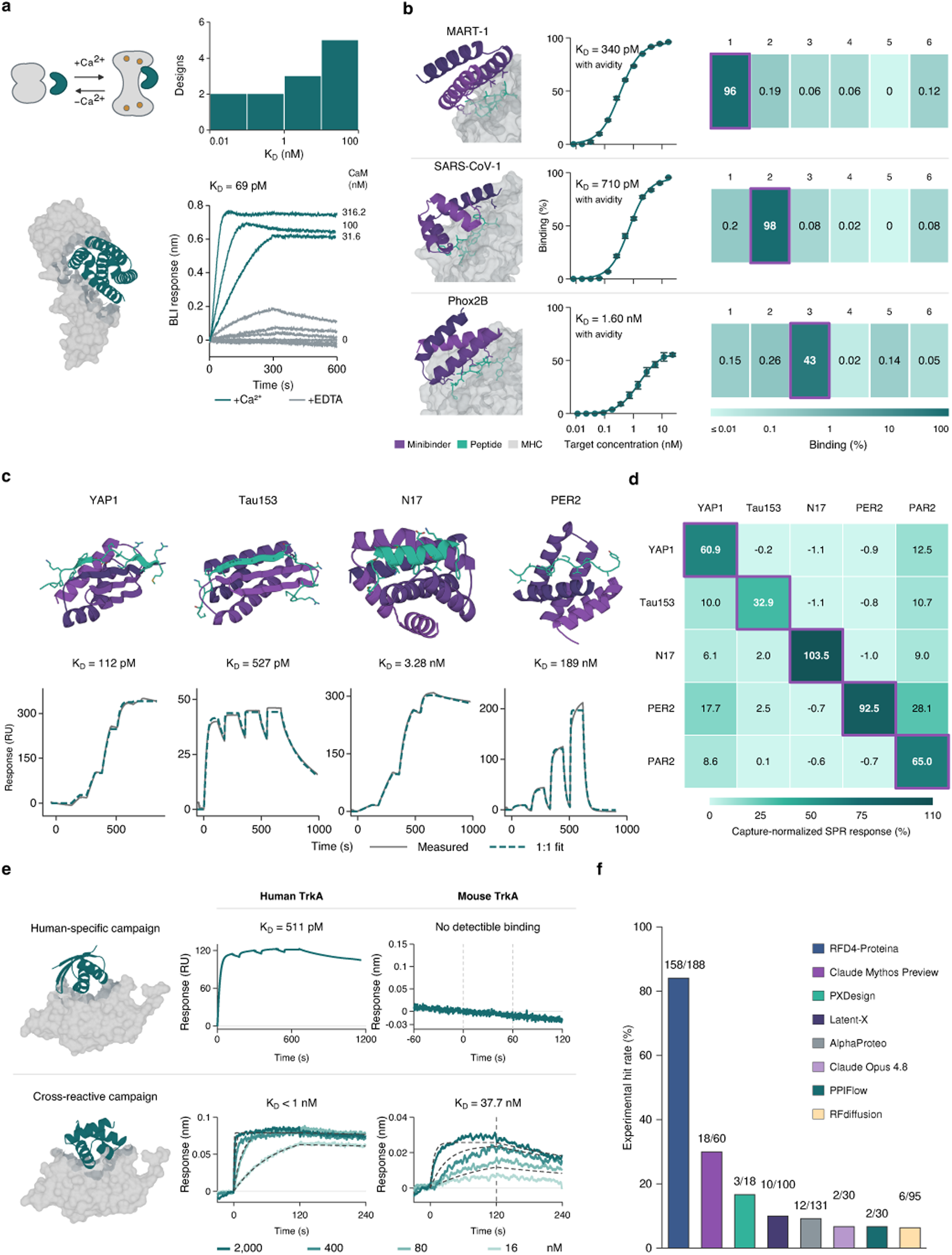
Binder specificity across protein targets and molecular states. **a,** Calcium-dependent calmodulin binding: schematic contrasting apo and calcium-bound calmodulin, affinity distribution of successes, and an example binder with biolayer interferometry (BLI) curves (apparent *K_D_* = 69 pM; 95% confidence interval, 3.4–92 pM). Curves show one replicate, with responses at 0, 31.6, 100, and 316.2 nM with calcium (teal; zero-concentration control in gray) or at 0, 31.6, 100, 316.2, and 1,000 nM with EDTA (gray). The affinity estimate combines two independent replicates and is conditional on the assay’s lower off-rate limit (Supplementary Section S13.1). **b,** Generated interfaces, measured yeast-display titrations and specificity measurements for MART-1, SARS-CoV-1 and Phox2B peptide–MHC binders. The displayed apparent *K_D_* values of 340 pM, 710 pM and 1.60 nM, respectively, are empirical half-maximal concentrations with avidity, measured by tetramer staining. Specificity across six pMHCs was measured in a separate single-concentration pMHC-tetramer screen. Peptide sequences and HLA alleles are shown in Supplementary Fig. 24. **c,** Refolded structures and SPR traces for YAP1, Tau153, N17 and PER2. Gray lines show measured responses and dashed teal lines show 1:1 kinetic fits (*K_D_* = 112 pM, 527 pM, 3.28 nM and 189 nM, respectively). **d,** Measured IDR specificity from a separate all-by-all SPR run. Rows show YAP1 B9, Tau153 G12, N17 B12, PER2 D8 and PAR2 E7 against their five peptide targets, in the same column order. Values show the SPR response at 1,000 nM binder as a percentage of the captured-peptide response (one measurement per pair). **e,** Human-TrkA-specific (upper row) and human/mouse-cross-reactive (lower row) designs, with columns showing the design objective, generated complex, and human- and mouse-TrkA binding measurements. For the human-specific design, the human-TrkA SPR single-cycle sensorgram has a fitted *K_D_* = 511 pM. The mouse-TrkA BLI trace was measured at one binder concentration (1.61 `µ`M); teal shows the loaded-tip response minus the matched bare Protein A response (ΔmTrkA). For the cross-species design, human and mouse TrkA BLI titrations give *K_D_ <* 1 nM and *K_D_* = 37.7 nM, respectively. The vertical dashed line at 120 s in the lower mouse-TrkA plot marks the start of dissociation. **f,** Experimental hit rates against human TrkA (158/188 RFD4-Proteina designs with *K_D_ <* 1 `µ`M) from various *de novo* design campaigns.

Calmodulin is a highly conserved protein that switches conformation in the presence of Ca^2+^, thereby participating in essential signaling cascades across all eukaryotes. These pathways are often mediated by conformation-dependent binding of other proteins to calmodulin. We used RFD4-Proteina to explicitly design for this behavior, aiming to bind only the Ca^2+^-bound form of calmodulin (Fig. 4a). Of 18 designs tested, 12 showed calcium-dependent binding with a quantified apparent *K_D_ <* 1 µM and no detectable binding in the absence of calcium. Apparent affinities in the presence of calcium ranged from 69 pM to 37 nM (Supplementary Table S31).

Peptide–MHC (pMHC) binding presents a distinct biochemical specificity challenge. Binders must distinguish the target peptide from other peptides presented by the same MHC molecule. Recognizing pMHCs is key to identifying cancerous or infected cells, but necessitates minimal binding to sequence-similar peptides from healthy cells. Previous work has shown that *de novo* designed minibinders can specifically bind pMHC complexes [40–42]. To evaluate whether RFD4-Proteina could engineer such precision, we designed binders against three on-target pMHC complexes (Fig. 4b). Yeast-display titrations with pMHC tetramers yielded apparent half-maximal binding concentrations of 340 pM, 710 pM, and 1.60 nM for binders targeting MART-1, SARS-CoV-1 membrane protein, and Phox2B peptides, respectively. We then screened for specificity against the other target peptides and three additional off-targets, obtaining over 100-fold specificity to the on-target in all cases. The off-targets for the Phox2B and SARS-CoV-1 peptides were especially challenging, differing from the on-targets at only 1 and 2 residues, respectively.

Intrinsically disordered regions (IDRs) are implicated in a wide range of disorders, spanning neurodegenerative disease, chronic pain, and cancer. Many therapeutic applications rely on binding to such regions, but they present a significant challenge to generative models, which typically reason only over structured proteins. Existing approaches rely on either predetermined template libraries based on fully extended conformations, or on the assumption that the target can form a stable secondary structural element when bound [43, 44]. We tested whether RFD4-Proteina’s conformational reasoning could enable binder design against eight IDR peptides: five known to adopt secondary structure when bound and three without such structural propensity (Fig. 4c; Supplementary Fig. 25). We obtained tight binders against all but one in each category. These include to our knowledge the first *de novo* designed binders against the tested epitopes of YAP1 and Tau, with the strongest hits having an affinity of 112 pM and 527 pM, respectively. Beyond binding, it is essential for potential IDR peptide therapeutics to form precise sidechain-level interactions, since there is no consistent structural signal to confer specificity through simpler mechanisms such as shape complementarity. To assess target specificity, we screened the best binders against five IDR peptides in an all-by-all comparison (Fig. 4d). Each binder showed a higher response to its intended target than to the off-target peptides, with the degree of discrimination varying among binders.

Tropomyosin receptor kinases (TrkA, TrkB, and TrkC) are promising targets in neuropathic pain and cancers including neuroblastoma [45]. Although human TrkA (hTrkA) is an established benchmark within *de novo* binder design [1, 34–36, 39, 46], controlling recognition across species and closely related paralogues presents a more demanding test. We therefore designed hTrkA binders to avoid mouse TrkA (mTrkA), which shares 86% sequence identity across the modeled target region, and the human paralogues TrkB and TrkC (Fig. 4e). We obtained genes encoding 94 designs and assayed the four strongest hTrkA binders against these off-targets, observing no binding in all cases. We then asked whether RFD4-Proteina could achieve both cross-reactivity against hTrkA and mTrkA as well as specificity against hTrkB and hTrkC. Of 94 designs, 18 exhibited the desired cross-reactivity and specificity patterns, with the strongest showing sub-nanomolar affinity to hTrkA and 37.7 nM against mTrkA. When considering the canonical benchmark task of unconstrained binding to hTrkA, RFD4-Proteina demonstrated a step change in experimental success, yielding 158 submicromolar hits out of 188 designs collectively (84%) compared to 18/60 (30%) hits for the next most successful campaign, Claude Mythos Preview with access to open-source protein design tools (Fig. 4f). We also designed 94 binders each to hTrkB and hTrkC, obtaining one sub-micromolar hit to TrkB (30.2 nM) and 20 screening hits for TrkC (precise affinities not yet determined; Supplementary Table S34). To our knowledge, these constitute the first computationally designed binders against either target.

### 5.2 Reasoning over non-protein modalities

In N-linked glycosylation, a common form of post-translational modification, glycans are attached to asparagine residues co- and post-translationally, and subsequently trimmed by enzymes in the ER and Golgi. Preventing this downstream processing triggers an immune response to the underprocessed glycan, enabling its use as a vaccine adjuvant [47]. Engineering this behavior requires a high level of molecular precision: *de novo* designs must confine steric shielding to the core GlcNAc to block cleavage while retaining terminal mannose accessibility for recognition (Fig. 5a). Prior work has established the ability to retain oligomannose-type glycans spanning Man5–Man8, but no designs completely blocked processing to Man9 or Glc1Man9 [48]. We asked whether RFD4-Proteina could extend this control to smaller proteins that retain predominantly Man9 and Glc1Man9 while preserving terminal mannose accessibility. RFD4-Proteina substantially increased *in silico* design success rates over the previous RFDAA-based method, and an initial plate-scale screen yielded glycoproteins that retained predominantly Man9 and Glc1Man9, as designed (Fig. 5b). Compared to previous campaigns, we found a higher success rate of underprocessing across all LC– MS/MS-detected structures, with 8/9 designs having *>*90% oligomannose on occupied sites (Fig. 5c) [48]. RFD4-Proteina achieved this control with substantially smaller proteins than was previously necessary, thereby providing additional flexibility for downstream applications.

**Fig. 5.**
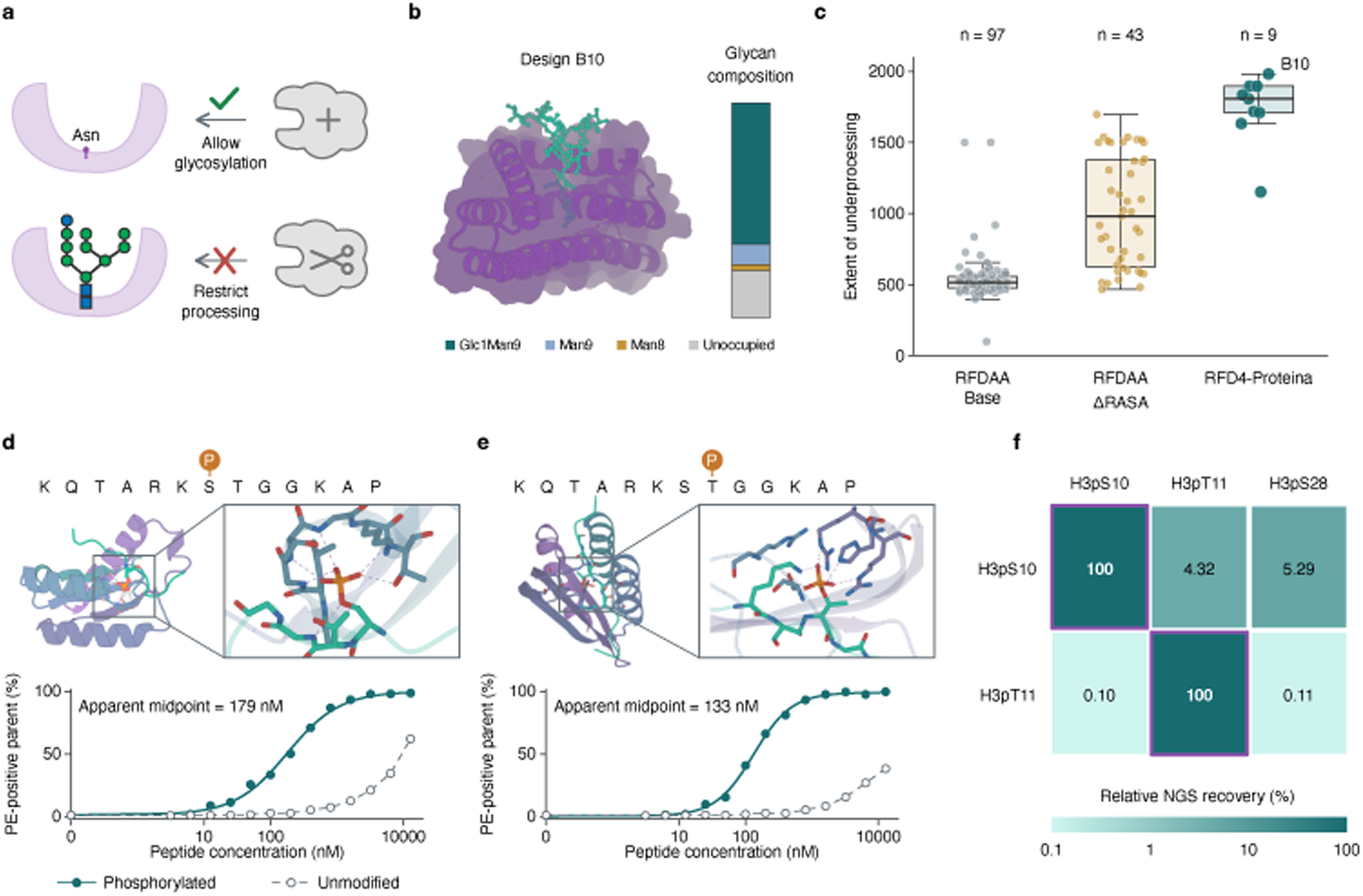
Design of glycosylation and phosphopeptide binding. **a,** Design objective: allow glycan attachment to asparagine (Asn) on the designed protein (purple; upper schematic), then shield the glycan core to restrict processing while leaving terminal mannose residues exposed (lower schematic). Gray shapes depict glycosylation and processing machinery. **b,** Design B10 with a modeled Man_9_GlcNAc_2_ glycan and glycan composition measured by LC–MS/MS. The stacked bar includes Glc_1_Man_9_ (teal), Man_9_ (blue), Man_8_ (orange), and unoccupied sites (gray); no other glycoforms were detected for this design. **c,** LC–MS/MS underprocessing-score distributions for 97 RFDAA, 43 ΔRASA-filtered RFDAA [48], and 9 RFD4-Proteina profiles. The score weights occupied glycoform abundances more highly for less-processed species (100–2000 scale). Profiles below 5% occupancy are excluded. **d,e,** Predicted structures and paired yeast-display titrations for example designs against H3pS10 and H3pT11, respectively. Dashed lines connect unmodified-peptide measurements, which did not plateau over the tested range. **f,** Sequencing-depth-normalized NGS recovery for the same H3pS10- and H3pT11-directed designs (rows) against H3pS10, H3pT11, and H3pS28 peptides (columns), from one selected population per condition. Values are normalized to recovery against the intended target (purple outline, 100%).

Phosphorylation, another post-translational modification, regulates molecular interactions across all domains of life. Recent work has demonstrated the design of proteins that exhibit phosphorylation-dependent binding to phosphotyrosine-containing peptides [49]. We used RFD4-Proteina to target two phosphorylated variants of the histone H3 tail, pS10 and pT11, obtaining phosphorylation-dependent binders against both sites (Fig. 5d–f). We further characterized selected pS10 and pT11 hits to test discrimination between adjacent phosphorylation sites. These hits preferentially bound their phosphorylated targets and each maintained selectivity against phosphorylation of the adjacent off-target site, highlighting the precise context-dependent binding engineered by RFD4-Proteina (Fig. 5f).

### 5.3 Designing bond-breaking and bond-forming enzymes

*De novo* enzyme design promises efficient, sustainable, and selective catalysts for countless chemical transformations. Recent work has shown that such design is possible through precise scaffolding of catalytic active sites [2, 50–58]. However, designing enzymes around more complex active sites and transition states remains a challenge. We tested whether RFD4-Proteina’s step-function *in silico* improvement at enzyme active-site scaffolding translates to two such challenges: hydrolytic bond cleavage by a bimetallic enzyme and stereoselective bond formation by a cysteine-based ligase (Fig. 6).

**Fig. 6.**
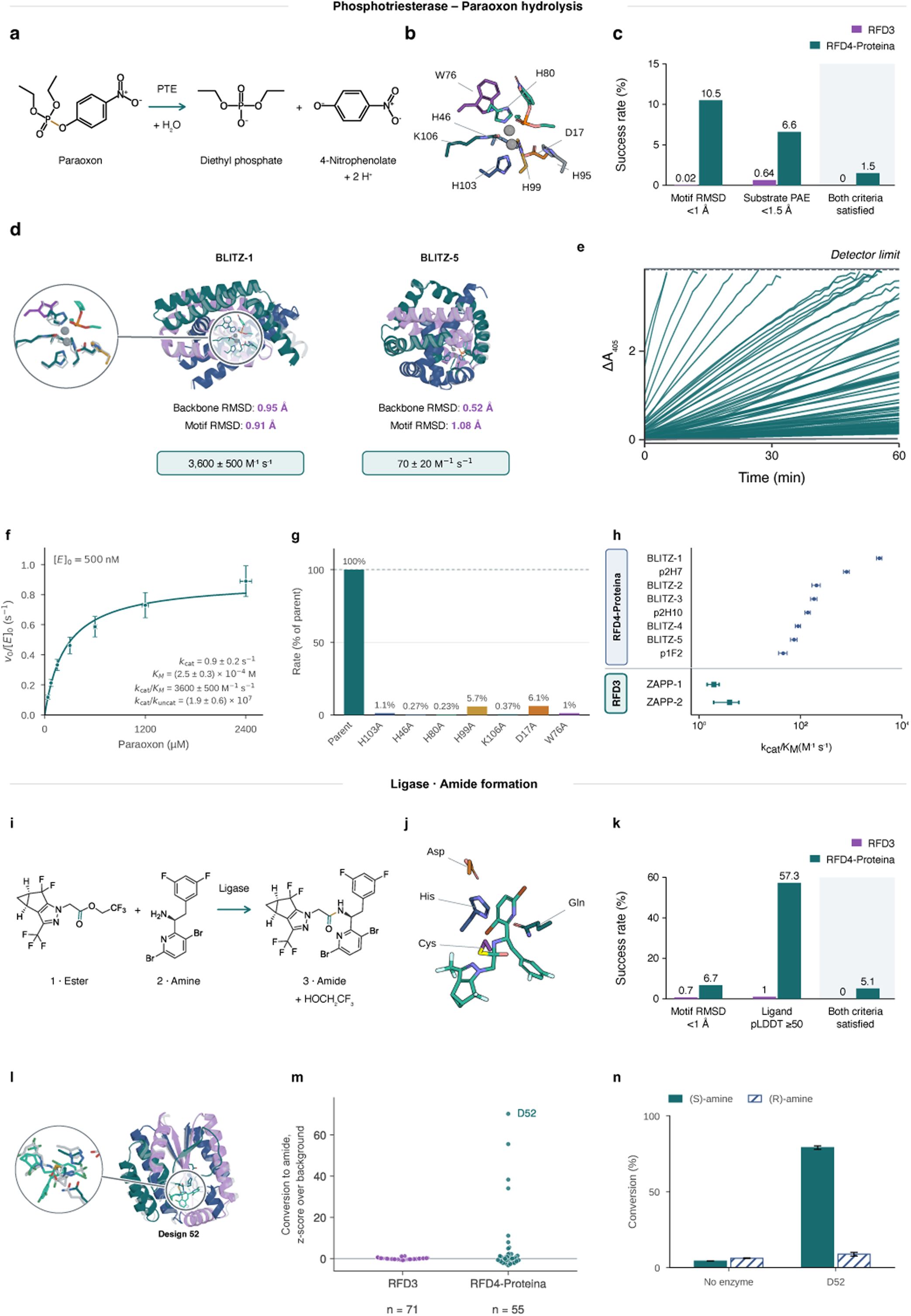
Enzyme design for bond-breaking and bond-forming chemistry. **a,** Paraoxon hydrolysis by a phosphotriesterase (PTE). **b,** Eight-residue BLITZ-1 active site with carbamylated lysine (KCX) and two zinc ions (gray spheres). **c,** Computational success after AF3 re-prediction, using the same 47 motif atoms and 18 substrate atoms for both models: motif RMSD *<* 1 Å and minimum protein–substrate PAE *<* 1.5 Å. Both criteria were satisfied by 1.5% of RFD4-Proteina designs and 0% of RFD3 designs in this benchmark. **d,** Active PTE designs (in color) and AF3 re-predictions (gray), with the BLITZ-1 active site magnified; boxes give *k*_cat_*/K_M_* (*±* SD) at 500 nM enzyme. **e,** Activity screen of 288 RFD4-Proteina designs (89 active, spanning 25 of 47 scaffold families); RFD3 yielded no active designs among 192 tested for this theozyme. **f,** BLITZ-1 kinetics at 500 nM enzyme. Points show mean turnover rates from three technical replicates and the curve shows the nominal Michaelis–Menten fit. Parameter annotations give Monte Carlo means and propagated SD. **g,** Paraoxon-hydrolysis rates for BLITZ-1 and alanine variants at seven first-shell catalytic residues. Rate is calculated as the slope of *A*_405_ over the first 60 s following the initial read, normalized to the parent slope and expressed as a percentage. See Supplementary Fig. 32 for full traces. **h,** *k*_cat_*/K_M_* of eight RFD4-Proteina designs and the published RFD3 designs ZAPP-1 and ZAPP-2. **i,** Amide formation from an activated ester and the (*S*)-amine, a step in lenacapavir synthesis. **j,** Designed ligase active site, with catalytic and oxyanion-stabilizing residues labeled. **k,** Computational bench-mark success on the ligase task: motif RMSD *<* 1 Å and ligand pLDDT *≥* 50. **l,** Design 52 aligned to its AF3 re-prediction (gray), with the active site magnified. **m,** Ligase campaign conversion relative to background, *z* = (*x̅*_design_ *− x̅*_background_)*/s*_background_. Each point is one design: 71 RFD3 designs screened in lysate and 55 RFD4-Proteina designs screened after purification. **n,** Amide conversion with enantiopure (*S*)-or (*R*)-amine for D52 and no-enzyme controls. Bars show means *±* population SD of three measurements.

We first tested whether RFD4-Proteina could design phosphotriesterases that hydrolyze paraoxon, an organophosphate pesticide and potent neurotoxin (Fig. 6a) [59]. Our previous RFD3 designs scaffolded six catalytic residues around two metal cofactors and the paraoxon substrate, yielding active enzymes with comparatively low catalytic efficiencies (up to 4 ± 3 M*^−^*^1^ s*^−^*^1^) [6]. We hypothesized that adding two more catalytic residues could improve catalysis, but would require more precise active site scaffolding (Fig. 6b). On this eight-residue theozyme, 1.5% of RFD4-Proteina designs passed common motif-RMSD and substrate-confidence criteria, compared with 0% for RFD3 (of a 10,000 design sample, Fig. 6c). Experimental screening yielded 89 active RFD4-Proteina designs among 288 tested, spanning 25 of 47 scaffold families, whereas none of 192 RFD3 designs showed activity (Fig. 6d,e). We characterized eight designs from five scaffold families, with the best design, BLITZ-1, achieving *k*_cat_*/K_M_* = 3,600±500 M*^−^*^1^ s*^−^*^1^ and *k*_cat_ = 0.9±0.2 s*^−^*^1^, a (1.9±0.6) × 10^7^- fold rate enhancement over uncatalyzed hydrolysis (Fig. 6f; Supplementary Fig. 31). The characterized hits exceeded the catalytic efficiencies of our earlier six-residue-theozyme phosphotriesterases by 10- to 1,000-fold (Fig. 6h). Alanine substitutions of the seven first-shell catalytic residues markedly reduced or abolished observed paraoxon-hydrolysis rates in the eluate screen (Fig. 6g; Supplementary Fig. 32), providing strong evidence that the catalysis occurred in the designed active site.

We next asked whether RFD4-Proteina could design bond-forming enzymes, an increasingly important tool for low-cost and sustainable synthesis of small-molecule therapeutics [60]. As a test case, we targeted a step in the synthesis of lenacapavir, a highly effective HIV-prevention drug whose manufacturing cost remains a barrier to global access [61]. The step involves amide bond formation between an activated ester and the (*S*)-enantiomer of a sterically demanding chiral amine (Fig. 6i). Such a transformation is challenging because the catalyst must efficiently favor the desired aminolysis pathway over competing hydrolysis of the activated ester.

We applied the strategy described in [62, 63], scaffolding a cysteine protease catalytic triad [57] with RFD4-Proteina (Fig. 6j) and using multi-state AF3 filtering to bias designs toward aminolysis. Once again, we found that RFD4-Proteina yielded orders of magnitude higher *in silico* success rates than RFD3 (Fig. 6k), and therefore obtained genes encoding 67 designs for experimental testing. Four of these showed clear amide bond formation with the desired (*S*)-amine above background (Supplementary Fig. 33); our previous campaign with RFD3 yielded no hits from 72 designs (Fig. 6m). The most effective ligase converted 79% of the (*S*)-amine to the desired amide and relies on the catalytic cysteine for activity, with a 12-fold reduction in conversion upon mutation to alanine [62]. This design is also highly enantioselective, showing a 28-fold preference for the desired (*S*)-amine over the undesired (*R*)-amine (Fig. 6n) [62]. The companion study by Salike et al. describes the ligase design methodology and structural determinants of activity in detail [62]. To our knowledge, these results provide the first proof-of-concept demonstration of a *de novo* designed enzyme catalyzing a step in active pharmaceutical ingredient (API) synthesis.

## 6 Discussion

We set out to enable the design of proteins that satisfy multiple interdependent chemical and geometric constraints. RFD4-Proteina generates molecular identity and conformation more efficiently and across a wider range of challenges than previous methods. Across four *in silico* protein design benchmarks, high-thinking RFD4-Proteina achieves higher computational success rates and produces 5.0–10.4-fold more distinct passing structural clusters per GPU-hour than the next-best methods, even when including the time to refold designs with a structure prediction oracle. RFD4- Proteina also generated passing designs for targets missed by every evaluated baseline: three small-molecule targets and three enzyme-scaffolding tasks (Fig. 2). RFD4-Proteina’s versatile conditioning framework enables its use for a variety of tasks beyond standard protein design, performing equivalently to or better than specialized tools without additional training (Fig. 3).

These *in silico* capabilities have translated into experimentally validated function across a range of design problems. We used RFD4-Proteina to design proteins specific to conformational states of calmodulin, peptide-MHC complexes, therapeutically relevant unstructured peptides, and homologues of the Trk protein family (Fig. 4). We also used RFD4-Proteina to precisely control glycosylation and to design binders against post-translational modifications (Fig. 5). Last, we used RFD4-Proteina to design enzymes that catalyze both bond-breaking and bond-forming reactions, yielding orders of magnitude improvement over prior methods (Fig. 6).

RFD4-Proteina integrates physics-based modeling with machine learning at multiple levels, using TMol’s differentiable, GPU-accelerated implementation of Rosetta for computational efficiency [9, 10]. First, fine-tuning the decoder with energy supervision yields lower-energy protein designs with fewer bond-angle deviations (Supplementary Fig. 8). Second, backpropagation of energy gradients enables direct steering of denoising trajectories using energy-based criteria. Third, energy-based criteria can guide extrinsic search (Fig. 3c; Supplementary Section S7.1.4). Collectively, this close interplay between generative machine learning and physics-based modeling is a step towards integrating these two long-standing approaches.

Methodological advances that contribute to RFD4-Proteina’s performance span data processing and preparation, model architecture, inference-time reasoning, and model acceleration. We built on AtomWorks, an open source tool for biomolecular data processing [64], to support new synthetic training sets and provide high-quality features and conditioning to downstream models. With the increase in quality and throughput of physical molecular simulations [65–68], we anticipate that performant data processing will be required to keep pace with the expansion of synthetic datasets. Further, the sophisticated sampling of conditions implemented within AtomWorks should lower the barrier for the community to fine-tune RFD4-Proteina on new datasets and design problems.

Architecturally, we strengthened the model by adding triangle multiplicative operations directly to the diffusion transformer and departing from the up-front processing that is prevalent in other biomolecular deep learning models [6, 13, 14, 19, 69, 70]. This approach grants the generative process more expressivity to reason over the interdependent features of a biomolecular complex. Furthermore, adding recycling loops and removing self conditioning improves computational design success, paralleling concurrent findings that recurrent depth is another axis for scaling flow and diffusion models [16, 71, 72]. By investigating how different parts of the diffusion process reason over different features [73–76], we built an inference-time reasoning protocol to adaptively allocate more exploration with search or more depth with recycles at critical points in the trajectory. We anticipate these reasoning techniques will be most useful for highly complex protein design campaigns.

We believe that RFD4-Proteina marks a step-change improvement in computational design capabilities and efficiency. For applications where *in silico* filters yield better wet-lab results, we anticipate that RFD4-Proteina will improve and accelerate experimental outcomes. In cases where existing computational metrics have not yet translated to experimental success, the ability to generate diverse candidates meeting arbitrary constraints provides a basis for systematic testing of novel filtering hypotheses. We hope that RFD4-Proteina will be a useful tool in both expanding the frontier of protein design and improving our understanding of biomolecules through design.

By releasing under a permissive license all code for data processing, model architecture, training, and inference-time reasoning, along with the RFD4-Proteina model weights, we hope to provide the community with an improved foundation for future biomolecular model development and chemical hypothesis testing.

## Code availability

Source code is available at https://github.com/RosettaCommons/RFD4-Proteina.

## Acknowledgments

We gratefully acknowledge the Jen-Hsun and Lori Huang Foundation for providing a computational grant that was transformative for this study. This work was funded by the Gates Foundation (INV-010680 to R.B., N.P.K. and D.B.; INV-043758 to R.I.B., M.C.L.H., N.K.M.B., Sh.S., P.V., J.B., A.C., D.K., R.K., Y.L., A.M., N.P.K. and D.B.; INV-092109 to D.B.; INV-092950 to A.P.G. and D.B.). This work was supported by a gift from Schmidt Sciences for Enzyme Design via Deep Learning awarded to David Baker. A.P.G. also acknowledges support from an EPSRC Centre-to-Centre Partnership with the Institute for Protein Design (EP/Z531157/1).

Additional support came from a gift from Microsoft (to R.B., D.L.S., Sh.S., E.S., B.R.K., R.K. and D.B.); the Open Philanthropy Project Improving Protein Design Fund (to A.M.C.G., B.C. and D.B.); the Grantham Foundation for enzyme design (to N.C., M.G., Sa.S., J.B., E.S.D. and B.R.K.); and the IPD Enzyme Design Fund (to J.C., Sa.S., S.M.W., K.W. and E.S.D.). This work was also supported by the National Science Foundation (CHE-2226466 to F.D. and D.B.); the National Institute of Allergy and Infectious Diseases of the National Institutes of Health (U19AI181881 to N.C., M.W.K., R.B., M.C.L.H., Sh.S., Y.L., N.P.K., F.D. and D.B.); and the Advanced Research Projects Agency for Health APECx Program (1AY1AX000036 to M.S.B., M.C.L.H., H.E.E., R.K., N.P.K., F.D. and D.B.). J.E.B., H.S. and D.B. were supported as part of the Matchmakers team through the Cancer Grand Challenges partnership, funded by Cancer Research UK (CGCATF-2023/100008), the National Cancer Institute (OT2CA297288-01) and the Mark Foundation for Cancer Research. D.B. was supported by the Howard Hughes Medical Institute. Z.X. was supported by a scholarship from the Tianjin Institute of Industrial Biotechnology, Chinese Academy of Sciences (TIB-CAS). N.C. was also supported by the Open Molecular Software Foundation. S.M.W. was supported by the National Science Foundation Graduate Research Fellowship Program (DGE-2140004). S.M.W., D.K. and D.B. were supported by the Defense Advanced Research Projects Agency (HR00112520029; CFDA No. 12.910).

We are grateful to Dr. John Walker, Dr. John Dillon, Dr. Trevor Laird and Dr. Mark Krook for helpful discussions on enzymatic synthesis of pharmaceutical intermediates. We thank Nicole Roullier for assistance with experimental screening. We thank David Binder for assistance with the antibody–antigen distillation dataset. We thank Andrew Kubaney, Raktim Mitra, Enisha Sehgal and Shivansh Goyal for useful conversations regarding DNA-binding protein design and benchmarking. We also thank Kyle Gion, Anthony Costa, and Ben Fauber for support with software acceleration and release. We thank Jenny Cronin, Kandise VanWormer and Luki Goldschmidt for supporting the running of the Institute for Protein Design.

## Notes

https://rfd4-proteina.org/

https://github.com/RosettaCommons/RFD4-Proteina

### Competing Interest Statement

The authors have declared no competing interest.

