## Supplemental Materials for "Biomolecular design through latent reasoning with RFD4-Proteina"

Supplementary Information

### Supplementary Information

### Part I: Model Description

#### S1 Training datasets and preprocessing

##### S1.1 Dataset mixtures

We trained the autoencoder and flow model on mixtures of experimental structures, predicted structures and synthetic molecular data. Dataset mixtures and sampling probabilities for each training stage are listed in Table S1; the training stages are described in Section S4.2.

**Table S1**: Training datasets and sampling probabilities.

| Dataset | Autoencoder (%) | Flow pretraining (%) | Flow fine-tuning (%) |
|---|---|---|---|
| AFDB monomers [37, 77] | 50 | 71.5 | — |
| PDB interfaces | 9 | 14.5 | 85 |
| Teddymer dimers [7] | 5 | 5 | — |
| PDB noncanonical/modified amino-acid environments | 4 | — | — |
| PDB glycans | 4 | — | — |
| Reglycosylated structures (Re-Glyco) [78] | 6 | — | — |
| Synthetic noncanonical/modified peptides | 4 | — | — |
| Noncanonical amino-acid scaffold distillation | 2 | — | — |
| Protein–DNA distillation | 6 | 0.3 | — |
| RNA monomer distillation | 6 | 1 | — |
| CP-SEA, AFDB-derived [79] | 1 | 0.5 | — |
| CP-SEA, PDB-derived [79] | 1 | 0.5 | — |
| PLINDER [80] | — | 3.5 | 10 |
| OMol-25 [65] | — | 1 | — |
| Antibody–antigen distillation | 0.5 | 0.5 | — |
| PDB antibody interfaces | 0.5 | 0.5 | 5 |
| TCR distillation | 0.5 | 0.5 | — |
| PDB TCR interfaces | 0.5 | 0.5 | — |
| PDB protein–DNA interfaces | — | 0.2 | — |
| Total | 100 | 100 | 100 |

Distilled datasets include ESMFold2 predictions of modified peptides, protein–DNA complexes and antibody–antigen complexes [15]; RNA monomer predictions, as described in the RF3/AtomWorks supplementary methods [64]; and T-cell receptor (TCR)–peptide–MHC predictions, as described elsewhere [33]. We used AtomWorks [64] for structure parsing and bond assignment. Within PDB interface streams, sampling weights account for interface clusters, molecular composition and alternate conformers. Of note, we use AtomWorks to ensure we sample alternate conformations when provided in the deposited crystal structure, rather than subsetting to the highest occupancy like prior methods [14, 19]. Dataset-specific filters, raw structures, sampling weights, parsing options, and training transforms are specified in the released configurations and code.

##### S1.2 Data selection and provenance

Experimental structures used for training were restricted to those deposited before 2024-01-01. Source locations and eligibility filters are recorded in the dataset configurations. We refer readers to `configs/releases/`, `configs/datasets/`, and the data-loading pipelines in the released code for additional implementation details.

#### S1.3 Data annotations

For efficiency, we pre-annotated datasets with CATH and interchain interaction labels. These interaction and fold class annotations were subsampled online during training to use as conditioning information (details in Section S2.1). Table S2 summarizes their availability across the training sources.

For CATH annotation, we searched each protein chain independently against the CATH v4.4 S95 domain representatives (95% sequence identity) [81] using Foldseek [82]. We adapted the public domain-annotation pipeline associated with TED [83, 84]. The primary differences are that we assign the selected reference's CATH code to the entire query chain without doing domain segmentation, and we assign CATH codes using weaker Foldseek hits identified as structural neighbors (labeled **N** classification in the public pipeline).

We also used AtomWorks to annotate multi-chain datasets with six geometric criteria adapted from PLIP [85]: hydrogen bonding, pi-stacking, pi-cation, salt bridges, hydrophobic interactions, and metal coordination.

**Table S2** Offline annotations available in the training sources.

| Dataset | CATH | Interchain interactions |
|---|---|---|
| PDB interfaces (general, antibody, TCR and protein–DNA) | ✓ | ✓ |
| AFDB protein monomers | ✓ | × |
| Teddymer dimers | ✓ | ✓ |
| PDB PTM crops | × | × |
| PDB glycans | ✓ | ✓ |
| Re-Glyco glycan distillation | ✓ | ✓ |
| NCAA/PTM peptide distillation | × | × |
| NCAA scaffold distillation | × | × |
| Protein–DNA distillation | ✓ | ✓ |
| RNA monomer distillation | × | × |
| CP-SEA (AFDB- and PDB-derived) | × | ✓ |
| PLINDER | ✓ | ✓ |
| OMol-25 | × | ✓ |
| Antibody–antigen and TCR distillation | × | ✓ |

### S2 Molecular representation and conditioning

This section describes how molecular structures and design constraints are represented and provided to the model.

#### S2.1 Conditioning

##### S2.1.1 Overview of supported conditioning

Conditioning on desired qualities steers generation toward designs with those properties. Supplementary Fig. 1 summarizes the supported conditions, and whether they apply at the atom, residue, or chain level. Their input representations are listed in Table S7; we provide a high-level description of each condition below.

- **Absolute coordinates** specify the positions of selected atoms in the shared global coordinate frame.
- **Internal coordinates** specify motif geometry while leaving each coordinate group's placement free (Section S2.1.2).
- **Indexing and unindexing** specify or leave free the sequence positions of motif residues; positions within an unindexed group can retain specified sequence gaps.
- **Sequence probabilities** specify residue identities, preferences or exclusions at selected positions (Section S2.1.4).

- **Hotspots** are target atoms within 4.5 Å of at least one binder atom. A subset of these contacts is provided during training.
- **Fold topology** specifies CATH class (secondary-structure composition), architecture (spatial arrangement of secondary-structure elements) or topology (their connectivity) [81]. Training-label assignment is described in Section S1.3.
- **Protein secondary structure** specifies helix, strand or loop assignments computed from the C$\alpha$ trace using P-SEA [86].
- **Nucleic-acid base-pair topology** specifies which nucleotides pair, including crossing pairs in pseudoknots, without prescribing their three-dimensional arrangement. Partners can lie within or between chains; explicitly unpaired positions are distinguished from unspecified positions. Training labels combine hydrogen-bond counts from AtomWorks' PLIP-style detector [64, 85] with geometric filters on base-plane separation, buckle and propeller angles.
- **Solvent accessibility** is the solvent-accessible surface area of an atom in the complex divided by that of the isolated atom, using a 1.4 Å probe. Relative areas below 0.02, from 0.02 to 0.1 inclusive, and above 0.1 define buried, intermediate and exposed states, respectively.
- **Per-chain centroids** specify approximate chain positions using the unweighted mean of their resolved atom coordinates after cropping. During training, each centroid receives independent isotropic Gaussian noise with a standard deviation of 5 Å per coordinate axis to allow variation in placement.
- **Hydrogen bonding** labels donor and acceptor atoms with a donor–acceptor distance of 0.5–4.1 Å and a donor–hydrogen–acceptor angle of at least 100°. Atoms can carry both roles.
- **$\pi$-stacking** labels atoms in aromatic rings whose centers are within 5.5 Å, whose planes are within 30° of parallel or perpendicular, and whose projected center offset is at most 2 Å in at least one ring plane.
- **Salt bridges** label oppositely charged atoms within 5.5 Å of one another.
- **Cation–$\pi$ interactions** label a cation and aromatic-ring atoms when the cation is within 6 Å of the ring center and its projection onto the ring plane is within 2 Å of that center. Cationic and aromatic roles are encoded separately, with a combined state for atoms participating in both roles.
- **Metal coordination** labels metal atoms and coordinating nitrogen, oxygen or sulfur atoms within 3 Å.
- **Hydrophobic contacts** label carbon atoms within 4 Å of one another, where each carbon is bonded only to carbon or hydrogen atoms.

The six interaction conditions above describe interchain contacts, annotated using AtomWorks implementations of PLIP-style criteria [64, 85] (Section S1.3). They identify participating atoms without specifying explicit partner pairs; hydrogen-bond and cation–$\pi$ inputs also distinguish interaction roles. Stereochemical constraints are encoded separately using idealized local coordinates (Section S2.1.3).

##### S2.1.2 Internal coordinate conditioning

A common requirement in protein design is the ability to provide the internal structure of multiple input motifs without specifying their relative positions in three-dimensional space. For example, in antibody design, one typically wants to provide the structures of the target and the framework while allowing the model to choose the correct dock. Prior methods typically accomplish this using all-by-all token-level distance conditioning within motif groups and not between them [1, 13, 87], while RF-antibody also includes dihedral information to provide residue orientations [33].

In this work, we instead introduce internal coordinate conditioning as a new method for accomplishing the same underlying task. Rather than converting the internal structures of the motifs to distances, we simply center and randomly rotate them before providing the resulting coordinate values to the model. This information

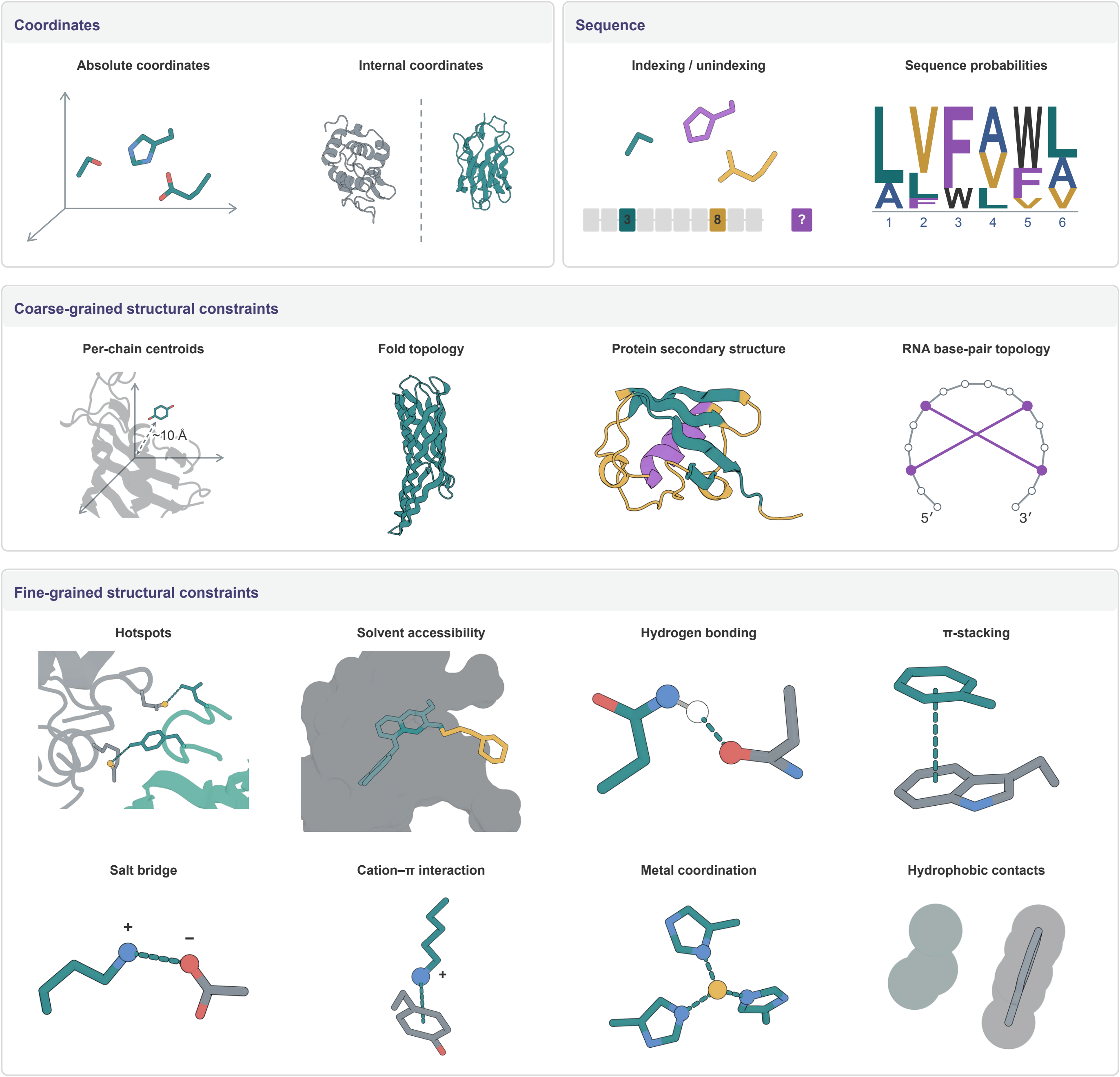

**Supplementary Fig. 1 Supported conditioning in RFD4-Proteina.** Illustrative inputs specify coordinates, sequence preferences, structural features and atom-level interaction roles. Internal coordinates allow independent motif placement; RNA base-pair topology specifies pairing partners, including crossing pairs in pseudoknots.

can be provided with atom-level precision and can distinguish between reflections. Supplementary Fig. 2a illustrates the independent centering and rotation of input motifs.

##### S2.1.3 Stereochemistry featurization

Various methods of featurizing chirality have been employed in the biomolecular machine learning literature, with mixed success. Rather than gradient-style features [69], full reference conformers [19], or direct chiral guidance vectors at inference time [70], with RFD4-Proteina we introduce a new form of stereochemical conditioning building upon the internal coordinate conditioning described in Section S2.1.2.

First, we compute an idealized conformation for the given stereochemistry type, such as tetrahedral chirality or E/Z stereochemistry. For example, the provided conformation would be an exact tetrahedron for common tetrahedral chirality. We perform this regularization so as not to leak any additional details to the model beyond the chirality. We also center the idealized conformation and rotationally align it to the current noisy coordinates. The resulting coordinate vectors constitute the model's stereochemistry signal (Supplementary Fig. 2b). We provide six such chirality tracks because one atom may participate in the idealized conformations of several different stereocenters.

To amplify the stereochemical signal, we supervise tetrahedral chirality with a signed-volume loss. Let $\hat{v}$ and $v_{\text{ref}}$ be the predicted and reference scalar triple products of the three edges from a given vertex, normalized by the product of the reference edge lengths. For each center, the penalty is

$$\ell_\chi = \max(0, m - s\hat{v})^2, \qquad m = \min(0.2, |v_{\text{ref}}|/2), \qquad s = \text{sign}(v_{\text{ref}}),$$

where $s$ specifies the reference handedness and $m$ is a positive margin. The penalty is zero when the predicted tetrahedron has the correct handedness and $s\hat{v} \geq m$, so that it penalizes nearly planar or mirrored tetrahedra without requiring their exact reference volume. Notably, this term does not include an explicit E/Z loss.

Using this formulation, RFD4-Proteina respects the input stereochemistry much better than previous design or structure prediction models (Fig. 3d). Importantly, adding a chiral loss alone does not lead to meaningful improvement: when retaining the loss term but removing the rotational alignment of the chiral features, tetrahedral adherence drops from 96.34% to 60.63% and E/Z adherence from 89.28% to 78.87%.

We assessed generalization to novel small molecules by comparing results on the deleaked validation set used in Fig. 3d to those on a subset of the small-molecule benchmark that was present in all models' training data (Supplementary Table S3). RFD4-Proteina had slightly higher stereochemical adherence on the unseen ligand set, whereas the other evaluated models had lower adherence. RFD4-Proteina does have a reduced PoseBusters pass rate on the held-out set, but this difference is the least among the models tested. It is worth noting that the ligands in the held-out set may be inherently more challenging, so that drops in stereochemical and geometric pass rates cannot be entirely attributed to overfitting.

**Table S3** Performance change between leaked and unseen ligands.

| Model | Tetrahedral chirality | E/Z adherence | PoseBusters |
|---|---|---|---|
| RFD3 + LigandMPNN | -4.2 | -9.6 | -14.3 |
| ProteinHunter | -18.2 | -13.7 | -11.5 |
| DISCO | -9.3 | -6.7 | -8.5 |
| BoltzGen | -16.2 | -14.2 | -8.2 |
| AF3 | -9.8 | -20.5 | -6.7 |
| RFD4-Proteina | **+1.0** | **+1.8** | **-5.3** |

Values are percentage-point differences (unseen minus training-overlap ligands); higher values indicate better generalization. AF3 denotes refolding of RFD4-Proteina outputs.

###### S2.1.4 Sequence distribution conditioning

RFD4-Proteina's sequence-structure codesign outperforms built-for-purpose methods such as ProteinMPNN and LigandMPNN across a range of scenarios [27, 88] (Fig. 3). Nevertheless, the logit-based sequence regression used in ProteinMPNN and related models offers the advantage that the position-wise residue type distributions can be easily biased, allowing users to exclude certain residue types and/or non-deterministically favor others. For example, this capability has proven valuable for

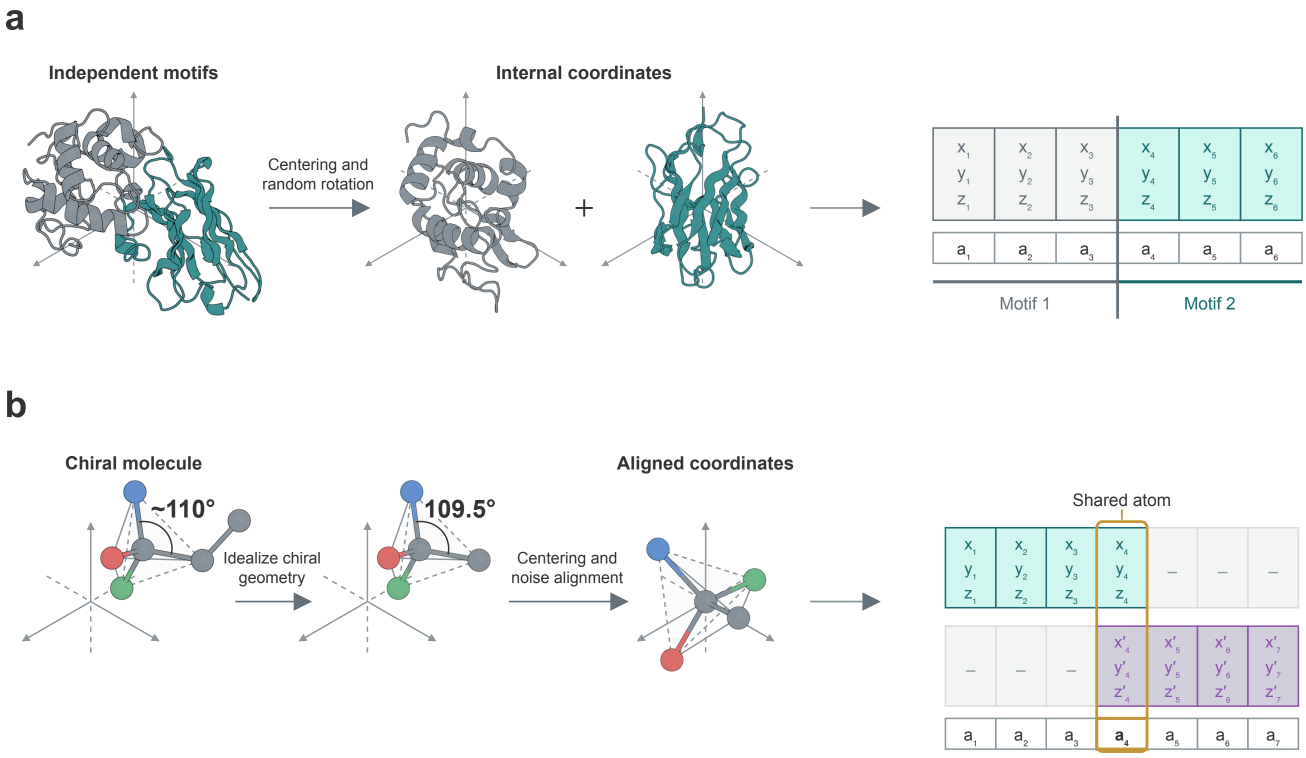

**Supplementary Fig. 2 Internal coordinate conditioning and chiral features.** **a**, Independently centered and rotated motif coordinates form atom-level features. **b**, Idealized chiral geometries are aligned to noisy coordinates. Separate tracks retain contributions from overlapping centers at shared atoms; two of six tracks are shown.

multistate design within the TiedMPNN methodology [89] and to enforce practical constraints such as avoiding cysteine residues that may impact expression in *E. coli*.

For this reason, RFD4-Proteina includes a conditioning type that allows users to specify arbitrary position-wise residue-type probabilities. This scheme is highly flexible, supporting scenarios from chain-wide conditioning such as excluding cysteines to position-specific constraints such as requiring a negatively charged amino acid in an active site. We also do not require the specified probability mass to sum to 1, enabling users to bias toward certain residue distributions without hard-enforcing them. By incorporating these constraints online as the model reasons over sequence and structure, the model can shape the backbone such that the given sequence constraints make chemical sense. Such a level of control is impossible with post-hoc methods like ProteinMPNN that cannot remodel the backbone.

For the model to learn this effectively, the sampled probabilities must be accurate when aggregated across training examples. Let $Y$ denote the ground-truth residue identity, $\pi_a = P(Y = a)$ its frequency in the training data, and $\mathbf{q}$ a complete probability vector. For any density $p_0(\mathbf{q})$ on normalized probability vectors with $\mathbb{E}_{p_0}[q_a] = \pi_a$, we can construct calibrated training pairs by sampling

$$p(\mathbf{q} \mid Y = a) = \frac{q_a}{\pi_a}\, p_0(\mathbf{q}), \tag{S1}$$

as Bayes' rule then gives

$$P(Y = a \mid \mathbf{q}) = \frac{\pi_a\, p(\mathbf{q} \mid Y = a)}{\sum_b \pi_b\, p(\mathbf{q} \mid Y = b)} = q_a \tag{S2}$$

as desired.

We therefore have significant flexibility in selecting $p_0(\mathbf{q})$, but for convenience of sampling, we use $p_0(\mathbf{q}) = \text{Dirichlet}(\alpha\boldsymbol{\pi})$ such that

$$\mathbf{q} \mid Y = a \sim \text{Dirichlet}(\alpha\boldsymbol{\pi} + \mathbf{e}_a), \tag{S3}$$

where $\mathbf{e}_a$ is the one-hot vector for the ground-truth residue. The total concentration $\alpha$ remains a free parameter, set to $\alpha = 3$ in RFD4-Proteina. We use the UniProt amino-acid frequencies to estimate $\pi$ and uniform frequencies for RNA and DNA. Sometimes a subset of residues is revealed to have probability 0, in which case the background frequencies for the remaining residues are renormalized to ensure that calibration remains correct. To enable partial specification of the conditioning vector $\mathbf{q}$, we also sample a mask that governs which probabilities are shown to the model.

#### S2.2 Design tasks

Training tasks are constructed by selecting which parts of an example to reveal to the model as conditioning and which to ask the network to generate. For example, we may reveal the backbone coordinates and sequence identity of a target chain, and ask the model to generate the sequence and structure of the masked binder.

Table S4 summarizes the supported tasks. Tasks are sampled only when the example contains the required molecular context. Design tasks control conditioning types that broadly define a problem: sequence conditioning, absolute coordinate conditioning, internal coordinate conditioning, and unindexing. Additional constraints such as binder design hotspots and RNA base-pairing are scoped to the associated design tasks. All other conditioning types are instead formulated as broader "design task modifiers" and sampled where applicable.

**Table S4** Design tasks used to train RFD4-Proteina.

| Task | Description |
|---|---|
| Unconditional generation | No sequence or structural conditioning. |
| Structure prediction | Generate structure given sequence. |
| Inverse folding | Generate sequence and rotamers given backbone coordinates. |
| Motif scaffolding | Scaffold multi-residue structural fragments. |
| Tip-atom scaffolding | Scaffold small groups of protein atomic coordinates. |
| Partial-ligand tip-atom scaffolding | Scaffold partial ligand coordinates and nearby protein tip-atoms. |
| Pocket tip-atom scaffolding | Scaffold protein tip atoms within pocket-like regions. |
| Partial-ligand conditioning | Generate the structure given sampled ligand fragments. |
| Binder design with target structure | Generate a binder given target coordinates and sampled interface hotspots. |
| Protein design with DNA structure | Generate a protein binder to a given DNA sequence and coordinates. |
| Protein design with DNA sequence | Generate a protein–DNA complex involving a given DNA sequence. |
| Nucleic-acid secondary-structure conditioning | Generate RNA given full or partial base-pairing information, optionally with a structural motif. |
| Antibody/TCR framework conditioning | Generate variable regions given framework geometry and, where present, a target. |

#### S2.3 Data representation

##### S2.3.1 Centering and coordinate frames

One question that all protein design models must contend with is how to center the input structures to obtain a regularized training signal. If examples are always centered on the origin, then whenever absolute coordinate conditioning is provided at inference time, this information will also necessarily specify a position relative to the designed protein's centroid. The natural solution is to center training examples on any absolute coordinate conditioning, thereby leaking nothing about their position within the overall ground-truth structure. However, as noted in [2], using the standard linear flow matching interpolant and origin-centered noise leaks the ground-truth centroid $\bar{x}_1 = \bar{x}_t/t$ to the model in training, leading to undesirable behavior at inference. RFdiffusion2 and RFdiffusion3 therefore fall back to global centering in training and introduce a mandatory "ORI token" for specifying motif placement relative to the global centroid [2, 6]. In [7] the centroid-leakage issue is instead addressed by adding a stochastic global offset to the input noise sample in training, allowing the ground-truth to be centered on any absolute coordinate conditioning.

In this work, we center on provided absolute coordinate conditioning in training, thus removing the requirement for an ORI token present in earlier RFdiffusion models. We also adapt the global offset from [7] to be computed independently per molecule and to apply only to regions without absolute coordinate conditioning (Supplementary Fig. 3). In this way, we prevent the model from back-calculating the offset vector based on how it affects regions with known ground-truth coordinates. To support users for whom the ORI token was a useful source of design control, we instead introduce *optional* per-chain centroid conditioning (Section S2.1), providing even greater flexibility than the ORI token.

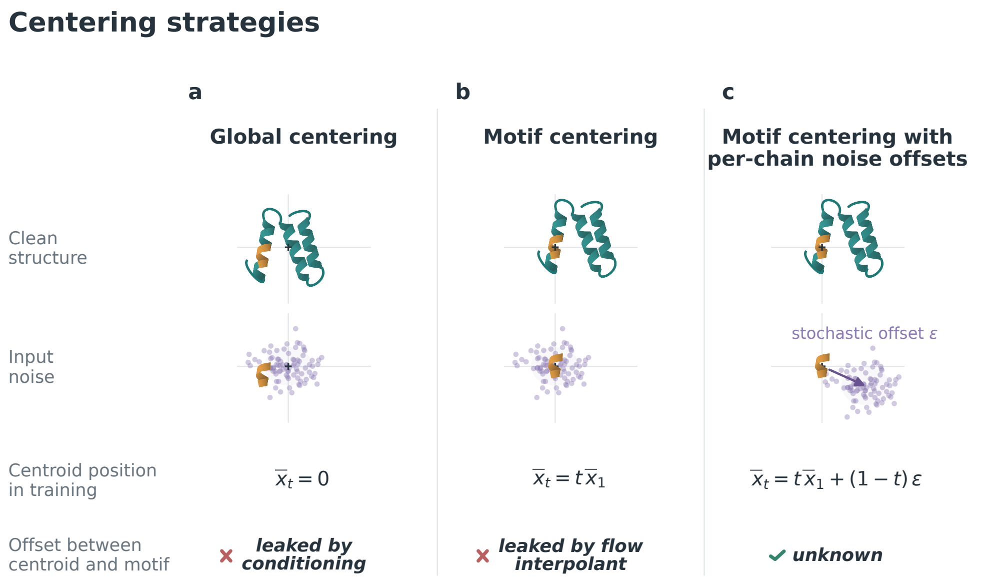

**Supplementary Fig. 3 Centering strategies.** **a**, With global centering, the coordinates of the motif specify its offset from the global centroid. **b**, With motif centering and origin-centered noise, the flow interpolant reveals the ground-truth centroid during training. **c**, Stochastic noise offsets enable motif centering without leaking the relative positions of the motif and the global centroid. These offsets apply only to regions without absolute coordinate conditioning to avoid revealing the offset $\epsilon$ to the model.

##### S2.3.2 Cropping

Our standard crops follow contiguous and spatial strategies similar to those of AlphaFold 3 [19]. Contiguous cropping randomly allocates the token budget across chains and selects a consecutive token segment from each, while spatial cropping retains the nearest tokens around a sampled query atom, which must be near an interface for multi-chain examples.

For protein and nucleic-acid binder tasks, a dedicated crop reserves space for the complete binding protein when it fits within a specified fraction of the token budget, then fills the remaining budget with nearby target context. An additional protein–DNA crop, introduced in RFdiffusion3 [6], samples a protein–DNA contact and retains contiguous protein and DNA segments around it, including base-paired partner nucleotides.

We use dynamic cropping to vary the token budget across training examples. For standard contiguous and spatial crops, with probability 0.8 we sample an integer crop budget uniformly from $\lceil 0.9N \rceil$ to $N$, where $N$ is the example's token count before cropping, then clip to the maximum crop size. Otherwise, we retain the available tokens up to the maximum budget. During pretraining, the flow model is trained with a crop size of 256, while in fine-tuning this is increased to 384. The autoencoder uses fixed 256-token crops.

##### S2.3.3 Bond graph positional encoding

The vast majority of biomolecular machine learning models encode within-polymer residue positions with clipped, signed index gaps, following the precedent set by AlphaFold2. Models that support non-polymer components typically introduce a separate feature that provides either pairwise bond existence and bond order, shortest-path distances within the bond graph, or a graph-based positional encoding [7, 19, 69]. When non-polymers are bonded to polymers, for example as a covalent modification of a protein chain, the bonded polymer residue typically participates in both the polymer and the non-polymer features. While these methods do in principle reveal the full bond graph, reconstructing it requires the model to navigate asymmetric treatments of polymers and non-polymers while aggregating connectivity information across many token pairs and/or input features.

In this work, we encode molecular connectivity with two pair features (Supplementary Fig. 4). The first is the unsigned distance between tokens in the bond graph, defined across the full molecule for both polymers and non-polymers. The second specifies the direction of canonical polymer bonds, taking values of $\pm 1$ between directly bonded protein or nucleic acid tokens. For peptide bonds, this distinguishes the residue contributing the carbonyl carbon from the one contributing the amide nitrogen. Separating connectivity from polymer direction also accommodates disulfide and isopeptide bonds between chains, for which bond-graph distances are meaningful but signed residue-index offsets are not.

In training, we compute bond-graph distance before cropping to preserve distances across intra-molecular crop boundaries. At inference, users thus have the flexibility to provide such gaps, enabling use-cases such as designing binders against cropped targets or providing an unindexed motif with a fixed-length internal sequence gap (e.g. N-X-T, where X is arbitrary).

We compared this novel connectivity featurization to AF3-style connectivity features, using the same ablation setup as in Section S3.1.6. We assembled a benchmark of 90 common post-translational modifications, and without providing any coordinate information, we unconditionally designed 8 monomers incorporating each of them. We found that adopting the RFD4-Proteina bond features sharply reduced the frequency of unphysically long bonds and severe clashes (Tables S5 and S6). In particular, the mean number of unphysically long polymer–nonpolymer junction bonds per design fell drastically from 0.68 to 0.01, aligning with the intuition that these junctions are precisely where per-chain connectivity features are most convoluted. Of note, this accuracy was obtained from featurization improvements alone, rather than introducing a bond loss as done in other models such as AlphaFold3 (where they introduce a protein-ligand bond loss in fine-tuning to address exactly this issue).

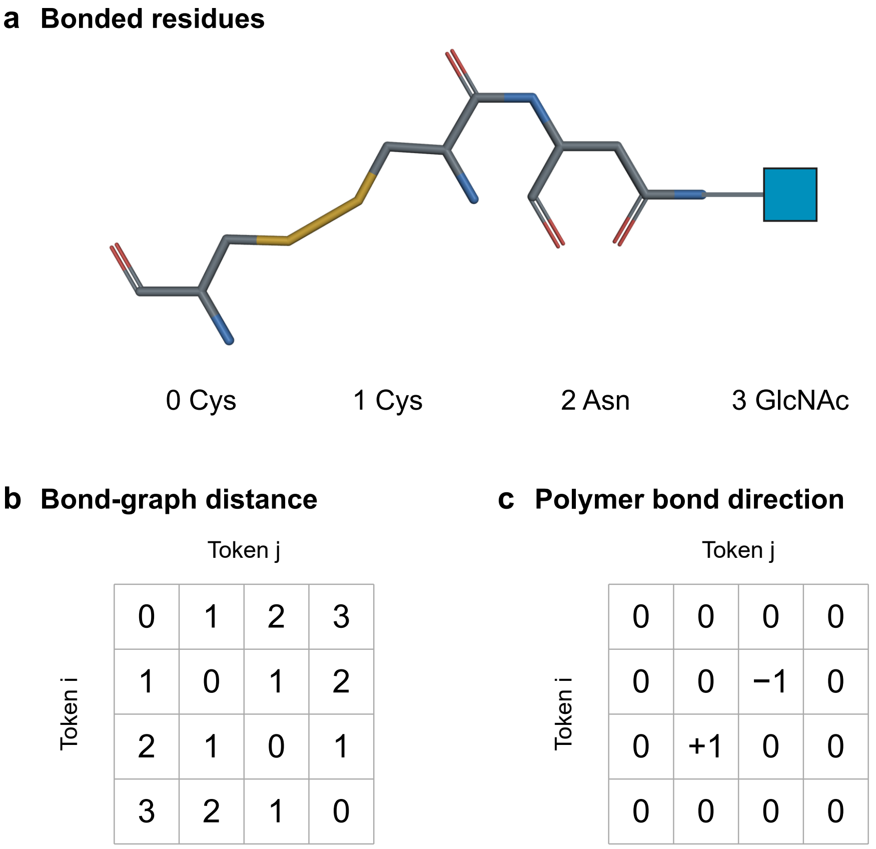

**Supplementary Fig. 4 Bond-graph distance and polymer bond direction. a**, Schematic of a complicated connectivity pattern: two cysteines connected by a disulfide bond, one of which is peptide-bonded to a glycosylated asparagine. **b**, Shortest-path bond graph distances between tokens. **c**, Polymer bond direction is nonzero only for the canonical peptide linkage between tokens 1 and 2.

| | Mean pairs per design | |
|---|---|---|
| Long-bond category | AF3-style | Ours |
| All bonds | 1.21 | $2.22 \times 10^{-2}$ |
| Within a residue | $1.32 \times 10^{-2}$ | $1.04 \times 10^{-2}$ |
| Between residues, same chain | 0.51 | $7 \times 10^{-4}$ |
| Polymer–nonpolymer junction | 0.68 | $1.11 \times 10^{-2}$ |

**Table S5 Unphysically long bonds in the PTM benchmark.** Each evaluation used eight designs per PTM across all 90 PTMs. Bonds are flagged above 1.8 Å for C/H/N/O-only pairs, 2.4 Å for other pairs drawn only from C/H/N/O/P/S, and 3.6 Å otherwise.

##### S2.3.4 Small-molecule featurization

We represent small molecules using element type, formal charge, bond connectivity and order, aromaticity, and local stereochemistry. Unlike AF3-like models [19], we do not embed atom-name strings or supply RDKit-generated reference conformers. Atom names depend on naming conventions rather than molecular chemistry, while reference conformers supply a precomputed geometric prior. Combined with synthetic molecular training data from OMol-25 [65] and consistent structure parsing and bond assignment using AtomWorks [64], these chemical inputs support higher PoseBusters pass rates than all evaluated baselines (Fig. 3d).

| Severe-clash category | Mean pairs per design | |
|---|---|---|
| | AF3-style | Ours |
| All pairs | 0.29 | $2.1 \times 10^{-3}$ |
| Within a polymer residue | $1.4 \times 10^{-3}$ | **0.00** |
| Within a nonpolymer residue | 0.28 | **0.00** |
| Polymer–polymer, different residues | $8.3 \times 10^{-3}$ | $1.4 \times 10^{-3}$ |
| Polymer–nonpolymer | $1.4 \times 10^{-3}$ | **0.00** |
| Nonpolymer–nonpolymer, different residues | $2.1 \times 10^{-3}$ | $7 \times 10^{-4}$ |

**Table S6 Severe clashes by partner type in the PTM benchmark.** Each evaluation used eight designs per PTM across all 90 PTMs. A severe clash is a nonbonded atom pair separated by less than 1.0 Å.

#### S2.4 Model input features

The model processes molecular tokens alongside additional in-context tokens that specify unindexed motifs. These conditioning tokens enable motif scaffolding without assigning the motif predefined sequence positions [2].

Input features are concatenated and linearly projected to initialize the single and pair representations. Selected conditioning features instead enter through adaptive layer normalization (Supplementary Fig. 5). Table S7 lists the complete feature inventory; Section S2.1 describes the supported conditions.

**Table S7**: Flow-model input features. Shapes omit the batch dimension and refer to encoded features before the shared input projections, or to stored recycled representations. $N$ is the padded token count, including in-context tokens; atom-level conditions use 37 slots per token. Type identifies the affected representation or adaptive layer normalization (adaLN). Recycled representations are projected and added separately; the padding mask controls attention rather than contributing input channels. Time embeddings condition both the single track and initial pair representation; CATH labels and the recycle indicator condition only the single track.

| **Feature & Shape** | **Type** | **Description** |
|---|---|---|
| Noisy token coordinates $[N, 3]$ | single | Current geometry |
| Noisy local latents $[N, 32]$ | single | Current sequence and conformation |
| Previous single representation $[N, 768]$ | single | Reuse token-level reasoning |
| Token type $[N, 32]$ | single | Molecular class |
| Token padding mask $[N]$ | single, pair | Exclude padding |
| Formal charge $[N, 16]$ | single | Charge of atomized tokens |
| Element $[N, 16]$ | single | Element of atomized tokens |
| Stereochemistry $[N, 18]$ | single | Six noise-aligned coordinate tracks |
| Absolute coordinates and masks $[N, 37 \times (3 + 1)]$ | single | Fixed atom positions |
| Internal coordinates and masks $[N, 37 \times (3 + 1)]$ | single | Motif shape with free placement |
| Coordinate-group labels $[N, 37]$ | single | Independent motif frames |
| Sequence probabilities and masks $[N, 28 \times 2]$ | single | Residue preferences and exclusions |

Continued on next page

| Feature & Shape | Type | Description |
|---|---|---|
| Hotspots<br>$[N, 37]$ | single | Target contact sites |
| Protein secondary structure<br>$[N, 16]$ | single | Local backbone structure |
| Nucleic-acid pairing state and mask<br>$[N, 2]$ | single | Paired and unpaired bases |
| Solvent accessibility<br>$[N, 37]$ | single | Burial and exposure |
| Chain centroids<br>$[N, 3]$ | single | Approximate chain placement |
| Hydrogen-bond roles<br>$[N, 74]$ | single | Donors and acceptors |
| $\pi$-stacking<br>$[N, 37]$ | single | Aromatic contacts |
| Salt bridges<br>$[N, 37]$ | single | Salt-bridge participation |
| Cation–$\pi$ roles<br>$[N, 74]$ | single | Aromatic and cationic partners |
| Metal coordination<br>$[N, 37]$ | single | Metal-contact participation |
| Hydrophobic contacts<br>$[N, 37]$ | single | Nonpolar contacts |
| Coordinate time embedding<br>[256] | adaLN | Coordinate noise level |
| Latent time embedding<br>[256] | adaLN | Latent noise level |
| Recycle indicator<br>[1] | adaLN | Mark conditioned recycling passes |
| CATH label embeddings<br>$[N, 96]$ | adaLN | Class, architecture and topology |
| Noisy token distances<br>$[N, N, 30]$ | pair | Current spatial relationships |
| Previous pair representation<br>$[N, N, 384]$ | pair | Reuse pair-level reasoning |
| Bond presence<br>$[N, N, 1]$ | pair | Non-backbone connectivity |
| Polymer bond direction<br>$[N, N, 1]$ | pair | Backbone directionality |
| Bond order<br>$[N, N, 5]$ | pair | Chemical bond type |
| Aromaticity<br>$[N, N, 1]$ | pair | Aromatic bonding |
| Bond-graph path length<br>$[N, N, 64]$ | pair | Chemical separation |
| Same index group<br>$[N, N, 1]$ | pair | Indexed versus unindexed context |
| Same molecule instance<br>$[N, N, 1]$ | pair | Molecular membership |
| Same molecular entity<br>$[N, N, 1]$ | pair | Repeated molecular identity |
| Nucleic-acid base pairs and masks<br>$[N, N, 2]$ | pair | Pairing partners |

### S3 Architecture

We started from the flow model and autoencoder architectures of Proteina-Complexa [7] and introduced the modifications described below. We focus on the core architectural changes and their ablations; the released code provides complete implementation details.

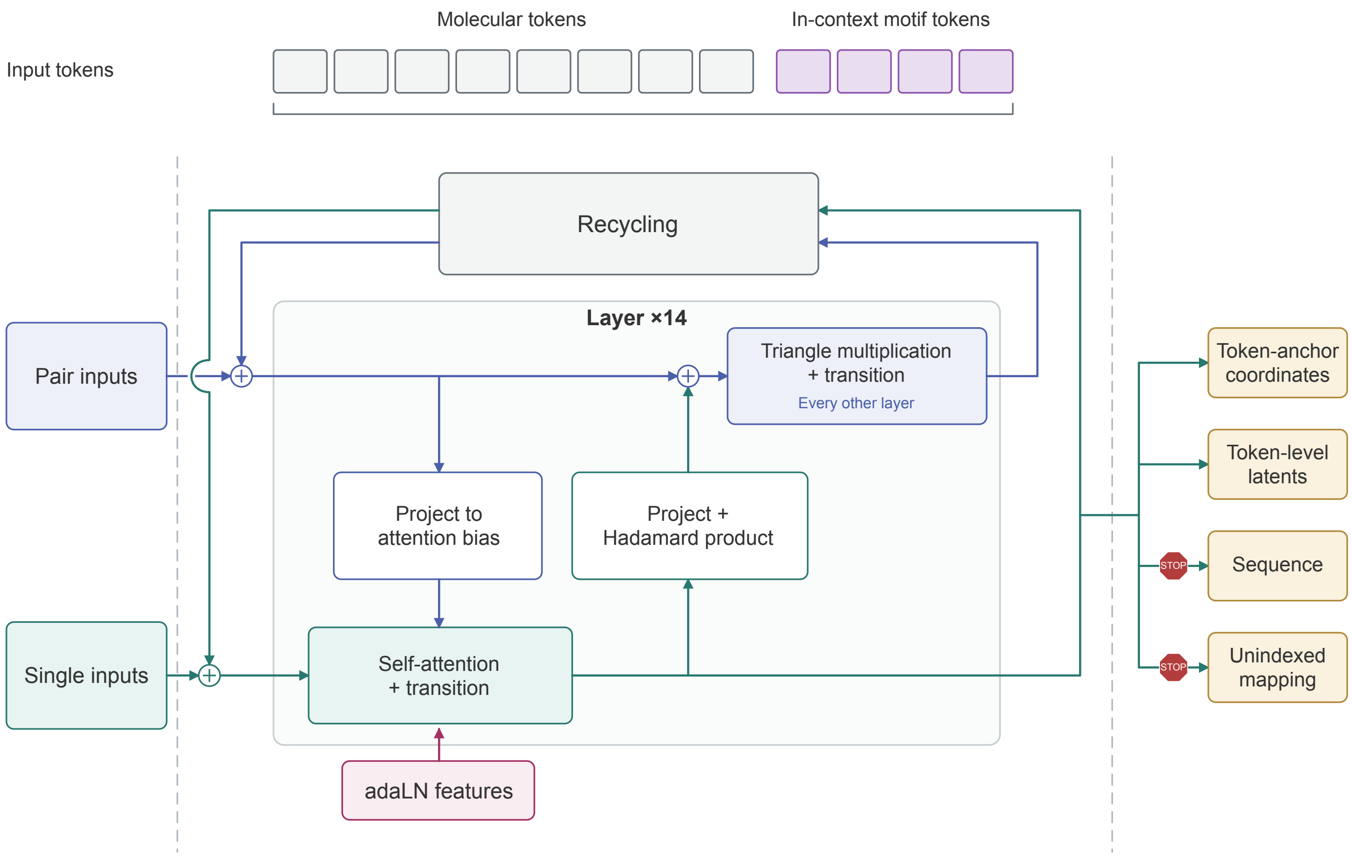

**Supplementary Fig. 5 Flow-model architecture.** The input sequence combines molecular tokens with additional in-context motif tokens for unindexed scaffolding. Single and pair input features are concatenated and projected to initialize token representations $s_i$ and token-pair representations $z_{ij}$. The single track applies pair-biased self-attention and a token-wise transition. Pair features are normalized and projected to an additive bias for each attention head. Adaptive layer normalization (adaLN) injects conditioning features into both operations. The pair track receives an additive update from pairwise Hadamard products of projected single features, followed by outgoing and incoming triangle multiplication and a pair transition. Each forward pass consists of 14 single-track layers, with pair updates every other layer. On recycling iterations after the first, the single and pair representations output by the previous pass are projected and added to the original input embeddings. Four readout heads operate on the single representation output by the final recycling iteration, predicting coordinate and latent velocities, residue identities, and motif correspondences. The two auxiliary heads receive detached representations, so their losses do not update the trunk.

#### S3.1 Flow model

The flow model processes molecular tokens alongside in-context conditioning tokens that specify motifs without predefined sequence positions (Supplementary Fig. 5; Section S2.4). We maintain a single representation $s_i$ for each token and a pair representation $z_{ij}$ for each token pair. We describe below the processing and interactions between these two tracks. Model dimensions and other salient hyperparameters are listed in Table S13.

##### S3.1.1 Single–pair communication

Single features update the pair track through an elementwise (Hadamard) product of learned projections:

$$\begin{aligned} u_i &= \mathrm{LayerNorm}(s_i), \quad a_i = W_a u_i, \quad b_i = W_b u_i, \\ z_{ij} &\leftarrow z_{ij} + a_i \odot b_j. \end{aligned} \tag{S4}$$

Here, $W_a$ and $W_b$ are bias-free projections to the pair dimension, and $\odot$ denotes elementwise multiplication. In contrast to earlier sequence–structure codesign models, which make use of additive updates [7], we use a Hadamard product to introduce bilinear interactions that allow each token to gate the other's contribution to the pair update. For channel $k$,

$$\frac{\partial(a_{ik}b_{jk})}{\partial a_{ik}} = b_{jk}, \qquad \frac{\partial(a_{ik}+b_{jk})}{\partial a_{ik}} = 1.$$

Thus, setting $b_{jk} = 0$ suppresses the multiplicative update regardless of $a_{ik}$, whereas suppressing the additive update requires cancellation, $b_{jk} = -a_{ik}$.

Direct ablation against additive updates supported the benefit of multiplicative single–pair communication for design success (Section S3.1.6).

##### S3.1.2 Pair–pair communication

After the single-to-pair update, the pair track applies outgoing and incoming triangle multiplicative updates, followed by a pair transition with residual connections [14]. Triangle updates allow each token pair to incorporate information from other pairs sharing a token. To balance performance and cost, the model updates the pair track after every other transformer block (Supplementary Fig. 5).

Ablation supported the contribution of triangle updates to backbone agreement and design success (Section S3.1.6).

##### S3.1.3 Single–single and pair–single communication

Tokens exchange information through multi-head self-attention on the single track. Normalized pair features are projected to an additive bias for each attention head, allowing the pair track to modulate which tokens attend to one another [7, 14] (Supplementary Fig. 5). For head $h$, the attention weights are

$$\alpha_{ij}^{h} = \mathrm{softmax}_j\left(\frac{(q_i^h)^\top k_j^h}{\sqrt{d_h}} + \beta_{ij}^h\right),$$

where $q_i^h$ and $k_j^h$ are normalized query and key projections from the single track, $d_h$ is the head dimension, and $\beta_{ij}^h$ is the projected pair bias. The weighted value sum is gated, projected and added to the single representation, followed by a token-wise transition. Adaptive normalization supplies conditioning information to both the attention and transition operations.

##### S3.1.4 Recycling within denoising steps

In AF3-like architectures, recycling refines single and pair representations in an upstream Pairformer trunk before diffusion [19]. We extend recycling directly into the diffusion module, applying recurrent refinement within each denoising step, analogous to concurrent developments in flow-based reasoning models [16, 17].

Earlier protein design models use self-conditioning to promote consistency across denoising steps, feeding the previous step's clean-state prediction into the next [1, 7, 12, 23, 90]. Our recycling instead refines hidden representations within each step while holding the noisy coordinates, latents and molecular conditions fixed. The same trunk is applied repeatedly, with normalized and projected single and pair states from the previous pass added to the original input embeddings. The final single representation supplies the prediction heads.

Algorithm S1 specifies one denoising step at inference. $X_t$ contains the noisy coordinates and latents, $t$ their noise times, and $C$ the molecular conditions. The scheduled depth $R_t$ counts all trunk passes, including the first. The operators $P_s$ and $P_z$ normalize and linearly project the recycled representations.

**Algorithm S1** Recycling within one denoising step

**Input:** Noisy state $X_t$, noise times $t$, conditions $C$, trunk-pass count $R_t$
**Output:** Velocities $v_x, v_\ell$, sequence logits $L_{\text{seq}}$, correspondence scores $M$

**function** DenoiseStep($X_t, t, C, R_t$):
$(s_0, z_0) \leftarrow \text{Embed}(X_t, t, C)$
$(s, z) \leftarrow (0, 0)$ *# Reset at each denoising step*
**for** $r = 1, \ldots, R_t$ **do**
  $s_{\text{in}} \leftarrow s_0 + P_s(s)$
  $z_{\text{in}} \leftarrow z_0 + P_z(z)$
  $c_r \leftarrow \text{ConditionForPass}(t, C, r, R_t)$
  $(s, z) \leftarrow \text{Trunk}(s_{\text{in}}, z_{\text{in}}; c_r)$ *# Shared weights*
**end for**
$(v_x, v_\ell) \leftarrow \text{FlowHeads}(s)$
$(L_{\text{seq}}, M) \leftarrow \text{AuxiliaryHeads}(s)$
**return** $(v_x, v_\ell, L_{\text{seq}}, M)$

During training, we sample the total number of trunk passes $R$, including the first pass, from a clamped Poisson distribution, similar to ESMFold2 [15]:

$$K \sim \text{Poisson}(3), \qquad R = \min(9, \max(1, K))\,.$$

We propagate gradients through only the final trunk pass, detaching the hidden states produced by earlier passes. Additional training settings are given in Section S4.2 and Table S13.

###### S3.1.5 Sequence and motif-mapping prediction heads

Two flow heads predict coordinate and latent velocities from the final single representation. Two auxiliary heads predict residue identities and unindexed motif correspondences. Both auxiliary heads receive detached single representations, so their losses train the heads without updating the trunk (Supplementary Fig. 5).

To preserve sequence constraints during decoding, we condition the autoencoder on user-specified residue identities where available and on sequence-head predictions elsewhere. This approach carries sequence-probability conditioning from the flow model into the decoded structure (Sections S2.1.4 and S3.2.1). We find that without such explicit conditioning, the model may alter the identity of sequence-conditioned residues upon decoding.

The motif-mapping head scores correspondences between unindexed motif tokens and generated tokens. Prior methods infer these correspondences from coordinates using greedy matching [2, 7, 12] or atom-level linear assignment [6]. However, internal-coordinate motifs lack a prescribed global placement, precluding direct matching by absolute coordinate proximity. We therefore predict correspondence scores from the final single representations:

$$m_{ij} = \frac{(W_q\,\text{sg}(s_i))^\top (W_k\,\text{sg}(s_j))}{\sqrt{d_s}}, \tag{S5}$$

where $i$ indexes motif tokens, $j$ indexes candidate generated tokens, $d_s$ is the single-track dimension and sg stops gradients. A per-chain Hungarian assignment maximizes the summed scores while assigning each motif token to a distinct generated token. The resulting mapping is returned with the design.

###### S3.1.6 Architectural ablations

We evaluated the contributions of triangle multiplication, multiplicative single–pair communication, and recycling using the variants below. Results are reported in Table S8.

**Reference model.** The reference uses Hadamard single-to-pair updates, outgoing and incoming triangle multiplication, and within-step recycling. For these experiments, the pair representation has 256 channels and is updated every third block, compared with 384 channels and updates every other block in the model.

**No triangle updates.** We remove outgoing and incoming triangle multiplication while retaining the single-to-pair update and pair transition (Section S3.1.2).

**Additive updates.** We replace $a_i \odot b_j$ in Equation S4 with $a_i + b_j$, retaining the learned projections and residual connection.

**Self-conditioning.** We replace within-step recycling with conditioning on the previous denoising step's clean-state prediction (Section S3.1.4).

**Evaluation procedure.** All variants use the same training seed, autoencoder, and diffusion batch size of 128. To reduce variance, we report the mean of two evaluations. All ablation runs—and baselines—are trained on 16 H200 NVIDIA GPUs; training end-to-end takes approximately three days. Recycling models use 60 sampling steps and 185 total trunk passes. The self-conditioning model uses 100 sampling steps with one trunk pass per step. We measure backbone RMSD, minimum interface PAE, and clustered pass rate across enzyme scaffolding, protein binding, small-molecule binding, and DNA binding tasks. The comparison also includes explorative modeling ($K = 4$). We report training losses averaged over the final epoch in Table S9.

**Results.** Replacing the Hadamard product with addition lowered clustered pass rates for enzyme scaffolding (7.48% to 5.32%), protein binding, and DNA binding (Table S8). Removing triangle multiplication lowered clustered pass rates sharply across all four tasks, including protein binding (4.32% to 0.73%) and enzyme scaffolding (7.48% to 2.49%). This decrease was accompanied by deteriorated backbone agreement between designed and predicted structures, with mean self-aligned backbone RMSD increasing from 1.13 to 1.68 Å for enzyme scaffolding and from 1.41 to 1.90 Å for small-molecule binding. Recycling, meanwhile, gave higher clustered pass rates and lower minimum interface PAE than self-conditioning across all four tasks under the evaluated sampling schedules.

We apply triangle updates after every other transformer block.

#### S3.2 Autoencoder

The autoencoder represents each supported residue by a token anchor $x_i \in \mathbb{R}^3$ and a latent $\ell_i \in \mathbb{R}^{32}$. The anchor retains the residue's global placement, while the latent encodes its identity and local structure. A shared encoder and decoder operate on proteins, nucleic acids, glycans and supported noncanonical or modified polymer residues.

##### S3.2.1 Architecture and features

We simplified the autoencoder architecture by removing the pair track [7]. Both the encoder and decoder use standard self-attention, token-wise gated transitions and residual connections. We complement this simpler architecture with additional synthetic training data, described in Section S1. Table S10 lists the input features.

Atoms occupy residue-specific slots, with masks identifying those present. The token anchor is C$\alpha$ for amino acids, C1$'$ for nucleotides and the anomeric carbon for sugars. The encoder receives atomic coordinates in the shared global frame as well as coordinates relative to the anchor, along with chemical features. These inputs are concatenated and projected into the transformer.

The decoder receives the latents, token anchors, molecular class, and any specified residue identities. Separate heads predict residue identity, atom positions relative to the anchor, and atom presence. This last head allows the decoder to represent the presence or absence of leaving atoms. Taken together, these outputs reconstruct the residue's identity and atomic structure.

| Task | Model | RMSD (Å) ↓ | Min. interface PAE (Å) ↓ | Clustered pass rate (%) ↑ |
|---|---|---|---|---|
| Enzyme scaffolding | Reference | **1.13** | **2.46** | **7.48** |
| | No triangle updates | 1.68 | 3.21 | 2.49 |
| | Self-conditioning | 1.86 | 2.89 | 4.93 |
| | Additive updates | 1.36 | 2.70 | 5.32 |
| | Explorative modeling ($K = 4$) | 1.22 | 2.50 | 6.87 |
| Protein binding | Reference | 25.37 | 13.21 | 4.32 |
| | No triangle updates | 27.96 | 15.04 | 0.73 |
| | Self-conditioning | 27.31 | 14.96 | 1.86 |
| | Additive updates | 26.53 | 13.76 | 3.85 |
| | Explorative modeling ($K = 4$) | **22.87** | **12.06** | **6.38** |
| Small-molecule binding | Reference | 1.41 | 2.67 | 7.46 |
| | No triangle updates | 1.90 | 3.47 | 2.86 |
| | Self-conditioning | 2.43 | 3.33 | 3.57 |
| | Additive updates | 1.59 | 2.76 | 7.71 |
| | Explorative modeling ($K = 4$) | **1.32** | **2.49** | **8.88** |
| DNA binding | Reference | 40.85 | 8.58 | 1.50 |
| | No triangle updates | 47.27 | 10.31 | 0.08 |
| | Self-conditioning | **40.00** | 9.47 | 0.75 |
| | Additive updates | 44.18 | 9.54 | 0.92 |
| | Explorative modeling ($K = 4$) | 41.46 | **8.47** | **1.67** |

**Table S8 Architecture and explorative-modeling ablations.** Values are means over two evaluations. For protein and DNA binding, RMSD measures binder backbone atoms after alignment on the conditioned target protein backbone or target DNA atoms, respectively. For enzyme scaffolding and small-molecule binding, RMSD measures the designed protein backbone after self-alignment.

| Model | Anchor loss ↓ | Latent loss ↓ |
|---|---|---|
| Reference | 0.62 | 48.61 |
| No triangle updates | 0.68 | 48.78 |
| Self-conditioning | 0.66 | 48.82 |
| Additive updates | 0.61 | 48.58 |
| Explorative modeling ($K = 4$) | 0.46 | 48.48 |

**Table S9 Training losses for the ablation models.** Values are means over the final training epoch.

**Table S10**: Autoencoder input features. $N$ is the padded token count. Atomic features use 37 slots per token; coordinates are expressed in nanometers.

| **Feature & Shape** | **Description** |
|---|---|
| **Encoder** | |
| Token type<br>$[N, 32]$ | Molecule type (protein, DNA, ...) |
| Atomic coordinates and masks<br>$[N, 37 \times (3 + 1)]$ | Geometry in the shared global frame |
| Anchor-relative coordinates and masks<br>$[N, 37 \times (3 + 1)]$ | Local geometry and orientation |
| Chain identifier<br>$[N, 1]$ | Chain membership |

Continued on next page

| Feature & Shape | Description |
|---|---|
| Incident bond-type counts $[N, 37 \times 10]$ | Ten bond categories per atom slot |
| Element and formal-charge embeddings $[N, 37 \times (8 + 16)]$ | Chemical identity: 8 element and 16 charge channels per slot |
| **Decoder** | |
| Token anchor coordinates $[N, 3]$ | Position of the anchor atom |
| Local latents $[N, 32]$ | Learned representation of identity and conformation |
| Token type $[N, 32]$ | Molecular class |
| Conditioned residue identity $[N, 147]$ | One-hot identity; zero if unspecified |

### S4 Training recipe

#### S4.1 Muon optimization

We use Muon for trainable matrix-valued weights, except input-feature embeddings and output heads, which use Adam. Muon approximately orthogonalizes momentum updates using Newton–Schulz iterations [91]. We clip gradients and maintain exponential moving averages of the model weights. Optimizer settings are given in Table S13.

#### S4.2 Training stages

We first trained the unified biomolecular autoencoder from scratch with 256-token crops and a global batch size of 512 (Section S4.3; dataset sampling ratios in Table S1). Complete training of the autoencoder required $\sim$ 4,000 GPU hours on NVIDIA H200s.

For the flow model, we distinguish between model development-scale runs and release-scale runs. For development, we used one noise candidate per training example ($K = 1$) for efficiency, training each run for approximately 4,000 NVIDIA H200 GPU-hours. For the release model, we scaled explorative modeling [24] with $K = 4$: we evaluated four independently noised candidates at the same sampled time and optimized the candidate with the lowest combined coordinate and latent flow-matching loss. Pretraining with 256-token crops to 320,000 optimizer steps thus required $\sim$ 16,000 GPU-hours (cost scales linearly with $K$). We then fine-tuned the model for another $\sim$ 2,000 GPU-hours with low-rank adaptation (LoRA) with rank 32, scaling factor 64, and no adapter dropout. Table S11 compares training compute and performance across the ablation reference, development, and release models. Both flow stages use the same frozen autoencoder; the released model includes this fine-tuning. Dataset mixtures are given in Table S1, and stage-specific settings are summarized in Table S12.

| Model | GPU-hours | Protein | Small molecule | Enzyme | DNA |
|---|---|---|---|---|---|
| Ablation reference | ∼1,200 | 4.3 | 7.5 | 7.5 | 1.5 |
| $K = 1$ (development-scale model) | ∼4,000 | 9.8 | 8.7 | 8.9 | 2.2 |
| $K = 4$ (release-scale model) | ∼18,000 | 10.6 | 14.3 | 12.1 | 3.5 |

**Table S11** Approximate flow-training compute and clustered pass rates (%) across model variants. Compute excludes autoencoder training and includes fine-tuning for RFD4-Proteina. The ablation reference is described in Section S3.1.6. The single-candidate ($K = 1$) and RFD4-Proteina ($K = 4$) models use high-thinking sampling, with 200 attempts per target across 24 protein-binding, 77 small-molecule-binding, 37 enzyme-scaffolding and three DNA-binding targets.

| Setting | Pretraining | Fine-tuning |
|---|---|---|
| Initialization | From scratch | Pretrained model |
| Optimizer updates | 320,000 | 60,000 additional |
| Trainable flow parameters | Full model | LoRA adapters in attention query/key/value and output projections |
| Maximum crop / padding | 256 tokens | 384 tokens |
| Global batch size | 512 examples | 256 examples |
| Learning rate | $10^{-4}$ | $5 \times 10^{-5}$ |
| Noise candidates per example | 4 | 4 |
| LoRA rank / scaling parameter | — | 32 / 64 |
| LoRA dropout | — | 0 |
| Recycle gradients | Final trunk pass only | Final trunk pass only |
| Estimated GPU-hours | 16,000 | 2,000 |

**Table S12** Flow-model training stages. Batch sizes count processed examples per optimizer update, including gradient accumulation but excluding the four noise candidates per example.

**Table S13**: Flow-model architecture and training settings. Table S12 gives stage-specific settings; Section S2.3 describes cropping.

| Setting | Value |
|---|---|
| **Architecture** | |
| Trunk depth | 14 blocks |
| Single / pair dimensions | 768 / 384 |
| Attention heads | 12, with 64 channels per head and query/key normalization |
| Token transition | SwiGLU, hidden dimension 3,072 |
| Normalization | RMSNorm, with adaptive conditioning |
| Pair updates | Every other block; Hadamard single–pair updates and triangle multiplication |
| Triangle hidden dimension | 256; bias-free projections |
| Conditioning dimension | 256 for each of the single and pair pathways |
| Latent dimension | 32 per token |
| Sequence head | 147 classes; no gradients to the trunk |
| **Flow matching and recycling** | |
| Predicted quantities | Coordinate and latent velocities |
| Coordinate time distribution | $0.98\,\mathrm{Beta}(1.9, 1) + 0.02\,\mathcal{U}(0, 1)$ |
| Latent time distribution | $0.8\,\mathrm{Beta}(1, 1.5) + 0.2\,\mathcal{U}(0, 1)$; sampled independently |
| Flow loss | Squared clean-state error, summed over dimensions and averaged over supervised tokens, with time weight $[(1 - t)^2 + 10^{-5}]^{-1}$ |
| Token weights | Base weight 1; add 2.5 for nonstandard chemical components and 2.5 for source tokens of unindexed motifs |
| Auxiliary loss weights | Sequence and motif-correspondence cross-entropy: 1 each; signed-volume chirality loss: 2.5 |
| Training trunk passes | Poisson(3), clamped to 1–9 total passes, including the first |
| Recycle gradients | Final trunk pass only |
| Recycle indicator | One channel; enabled on passes after the first with probability 0.5 per training step |
| **Optimization** | |
| Optimizers | Pretraining: Muon and Adam; LoRA fine-tuning: Muon |
| Learning-rate warmup | Pretraining Adam: 10,000 optimizer updates; Muon: none in either stage |
| Muon | Momentum 0.95; Nesterov; weight decay 0.1; 5 Newton–Schulz iterations |

Table S13 (continued)

| Setting | Value |
|---|---|
| Adam (pretraining) | $\beta_1 = 0.9$, $\beta_2 = 0.999$, $\epsilon = 10^{-8}$; no weight decay |
| Gradient clipping | Norm threshold 1.0 per optimizer |
| Precision | BF16 mixed precision |

#### S4.3 Autoencoder training

We trained the autoencoder to reconstruct atomic coordinates, residue identities, and which atoms are present while applying a KL penalty to constrain the latent distribution toward a standard normal prior. For reconstruction, we supervise with a weighted average across molecular classes (e.g., protein, DNA, RNA, glycans) to combat the intrinsic class imbalance. Tables S1 and S14 give the dataset mixture and training settings, respectively.

We also fine-tuned the autoencoder to minimize Rosetta energies. Section S4.3.1 describes this fine-tuning procedure; Section S6.1 evaluates its effect on generated structures. We provide both the energy-supervised and reconstruction-only decoders for use at inference.

**Table S14**: Autoencoder architecture and pretraining settings. Section S4.3.1 describes Rosetta energy fine-tuning.

| Setting | Value |
|---|---|
| **Architecture** | |
| Encoder / decoder depth | 12 transformer layers each |
| Hidden dimension | 768 |
| Attention heads | 12, with 64 channels per head and query/key normalization |
| Token transition | SwiGLU, hidden dimension 3,072 |
| Pair representation | None in either encoder or decoder |
| Latent dimension | 32 per token |
| **Training** | |
| Initialization | From scratch |
| Approximate GPU-hours | ~4,000 |
| Global batch size | 512 examples; no gradient accumulation |
| Crop size | 256 tokens |
| Sequence-conditioned reconstruction task | 10% of training examples |
| Coordinate loss | Squared error after matching equivalent atoms; weight 1 |
| Sequence loss | Cross-entropy; weight 1 |
| Atom-presence loss | Binary cross-entropy; weight 1 |
| KL penalty | Weight $10^{-4}$; no annealing |
| Reconstruction balancing | Residue losses: 75% token average, 25% equal-class average |
| **Optimization** | |
| Optimizers | Muon for matrices; Adam for input embeddings, output heads and non-matrix parameters |
| Learning rate | $10^{-4}$ for both optimizers; no warmup |
| Gradient clipping | Norm threshold 1.0 per optimizer |
| Precision | BF16 mixed precision |

##### S4.3.1 Energy-supervised autoencoder fine-tuning

We supplemented the autoencoder objectives described in Section S4.3 with reconstruction of constrained-minimized structures and direct Rosetta energy supervision [9, 10]. Starting from the pretrained autoencoder, we trained low-rank adapters in the decoder while keeping the encoder and original decoder weights fixed. Each adapted matrix was parameterized as $W = W_0 + (\alpha/r)BA$, with rank $r = 16$ and scaling parameter $\alpha = 32$. Training used AFDB monomers, a global batch size of 256, and a learning rate of $10^{-4}$ with 100 warm-up updates. We fine-tuned for 14,000 updates.

**Structure preparation and reconstruction targets.** We prepared structures for Rosetta energy evaluation using the beta2016 potential implemented in TMol, adding missing hydrogens and optimizing their orientations. The code provides the complete structure-preparation procedure.

Let $P(x)$ denote the prepared all-atom structure corresponding to input coordinates $x$. We generated reconstruction targets on the fly using up to 200 Cartesian minimization iterations, fixing C$\alpha$ positions and restraining other heavy atoms to their input positions:

$$y^\star \approx \underset{y:\, y_{\mathrm{C}\alpha}=P(x)_{\mathrm{C}\alpha}}{\arg\min} \left[ E_{\mathrm{Rosetta}}(y) + \frac{w}{\sigma^2} \sum_{i \in \mathcal{H} \setminus \mathrm{C}\alpha} \|y_i - P(x)_i\|_2^2 \right], \tag{S6}$$

where $\mathcal{H}$ is the set of heavy atoms, $w = 1$ and $\sigma = 0.5$ Å. Hydrogens were unrestrained. We mapped $y^\star$ back to the observed input atoms and detached it from the computation graph to obtain the reconstruction target $x^\star$. The encoder received the original coordinates $x$, whereas the symmetry-aware coordinate loss used $x^\star$.

**Energy normalization.** We retained the autoencoder losses described in Section S4.3, using the prepared reconstruction targets $x^\star$, and added a Rosetta energy term normalized by the number of scored protein residues:

$$\mathcal{L}_{\mathrm{energy}} = \lambda_E \frac{E_{\mathrm{Rosetta}}(P(\hat{x}))}{N_{\mathrm{res}}}, \tag{S7}$$

where $\hat{x}$ denotes decoded coordinates, $N_{\mathrm{res}}$ is the number of scored protein residues and $\lambda_E = 3 \times 10^{-4}$. Normalizing per residue prevents the energy term from scaling directly with protein length.

### S5 Inference

#### S5.1 Sampling

##### S5.1.1 Stochastic TSR sampling

We sample the backbone coordinates and local atomistic latents with a stochastic extension of temporal score rescaling (TSR) [92], which Proteina-Pronto adapted and validated for La-Proteina and Proteina-Complexa [93], motivating its adoption here. For a linear Gaussian flow interpolant, the intermediate score is recovered from the learned velocity $\mathbf{v}_\theta$ and current position $\mathbf{y}_t$ as [94, 95]

$$\mathbf{s}_\theta(\mathbf{y}_t, t) = \frac{t\mathbf{v}_\theta(\mathbf{y}_t, t) - \mathbf{y}_t}{1 - t}. \tag{S8}$$

TSR rescales this score by

$$r_t(k, \sigma) = \frac{\mathrm{SNR}(t)\sigma^2 + 1}{\mathrm{SNR}(t)\sigma^2/k + 1}, \qquad \mathrm{SNR}(t) = \frac{t^2}{(1-t)^2}, \tag{S9}$$

and converts it back to the tempered velocity $\widetilde{\mathbf{v}}_\theta = [\mathbf{y}_t + (1-t)r_t\mathbf{s}_\theta]/t$. Thus $k = 1$ recovers the original velocity, whereas $k > 1$ increasingly sharpens the distribution toward the end of the trajectory.

Our construction injects stochasticity into TSR sampling analogously to Karras et al. [96], in which a deterministic ODE is augmented with a "Langevin-like churn" term:

$$d\mathbf{y}_t = \underbrace{\mathbf{v}_{\mathrm{ODE}}(\mathbf{y}_t, t)\, dt}_{\text{ODE transport}} + \underbrace{\beta(t)\nabla_{\mathbf{y}} \log p_t(\mathbf{y}_t)\, dt}_{\text{score drift}} + \underbrace{\sqrt{2\beta(t)}\, d\mathbf{W}_t}_{\text{noise injection}}, \tag{S10}$$

where $p_t$ denotes the ODE marginal density, $\mathbf{W}_t$ is a standard Wiener process, and $\beta(t) \geq 0$ is a free parameter that controls the churn strength. With the exact ODE-marginal score, this construction preserves the ODE marginals while injecting stochasticity. Our practical sampler uses the TSR-rescaled model score as an approximation. In La-Proteina and Proteina-Complexa [7, 12], the base ODE is the linear Gaussian flow interpolant, while in RFD4-Proteina we use the ODE defined by the TSR-rescaled score $\widetilde{\mathbf{s}}_\theta = r_t \mathbf{s}_\theta$. Empirically, we find it necessary to adjust the per-modality noise injection schedules $\beta(t)$ that were tuned in La Proteina for a standard non-TSR SDE (Table S15).

In summary, for a step of size $\Delta t$, our Euler–Maruyama update is therefore [97]

$$\mathbf{y}_{t+\Delta t} = \mathbf{y}_t + [\widetilde{\mathbf{v}}_\theta + g(t)\widetilde{\mathbf{s}}_\theta]\, \Delta t + \sqrt{2g(t)\Delta t}\, \boldsymbol{\epsilon}, \qquad \boldsymbol{\epsilon} \sim \mathcal{N}(\mathbf{0}, \mathbf{I}), \tag{S11}$$

where $g(t) = \beta(t)/r_t^2$. We use this update for both backbone and latent sampling. With this schedule, we find that additional temperature-lowering methods are not necessary, so we omit the noise-downweighting constant used in prior work [12].

We compared five sampler configurations on an unconditional design benchmark (Table S15). We observed that reweighting $\beta(t)$ was essential for stochastic TSR, and that with this new schedule stochastic TSR performed slightly better than ODE TSR sampling. We found that 400-step SDE sampling performed slightly better than stochastic TSR sampling, but that its schedule could not be shortened to realize the efficiency gains of TSR sampling.

| Sampler | $g(t)$ | Steps | Noise downweighting | Clustered pass (%) | Time (s) |
|---|---|---|---|---|---|
| Stochastic TSR | $\beta(t)/r_t^2$ | 60 | No | 34.7 | 2.99 |
| Stochastic TSR (prior schedule) | $\beta(t)$ | 60 | Yes | 0.0 | 2.97 |
| ODE TSR | 0 | 60 | No | 33.3 | 3.04 |
| Full-length SDE | $\beta(t)$ | 400 | Yes | 37.3 | 17.68 |
| Shortened SDE | $\beta(t)$ | 60 | Yes | 0.0 | 2.95 |

**Table S15 Unconditional sampling ablation.** Each sampler generated 150 unconditional designs (50 each at 100, 150 and 250 residues). The pass criterion was AF3 scRMSD $\leq$ 1.5 Å. Noise downweighting multiplies the injected noise variance by 0.3 for backbone coordinates and 0.1 for local latents; otherwise the multiplier is one.

We next compared these sampling methods on the task of searching for hydrogen bonds to a small molecule (Table S16). We found that stochastic TSR was slightly more effective than full-length SDE sampling, while both the original $\beta(t)$ schedule and a fewer-step SDE setup were much less effective. Taken together, these results motivate the use of this stochastic TSR formulation for efficient and performant inference-time sampling.

##### S5.1.2 Effect of sampling parameters on sequence distribution

In Equation S9 there are two free parameters, $k$ and $\sigma$. While both can be thought of as controlling sampling "temperature" to some extent, $k$ in particular provides interpretable control, with $k = 1$ recovering ordinary SDE sampling. To quantify this property, we analyzed the sequence distribution generated by RFD4-Proteina in unconditional generation (Supplementary Fig. 6). When setting $k = 1$ for both the token anchors and latents, we find that the resulting sequence distribution approximately matches that found in UniProt, indicating that the model has correctly learned the underlying data distribution. When we then use the default

| Sampler | $g(t)$ | Steps | Noise downweighting | Hydrogen bonds |
|---|---|---|---|---|
| Stochastic TSR (default) | $\beta(t)/r_t^2$ | 60 | No | 14.24 |
| Stochastic TSR (prior schedule) | $\beta(t)$ | 60 | Yes | 5.18 |
| Full-length SDE | $\beta(t)$ | 400 | Yes | 12.72 |
| Shortened SDE | $\beta(t)$ | 60 | Yes | 3.31 |

**Table S16 Hydrogen-bond search ablation.** Values are mean protein–ligand hydrogen-bond atom-pair counts for a 145-residue SAM-binding protein. Each sampler uses 50 independent search runs, beam width two, branching factor two, and checkpoints at 25% and 50% of integration steps, with two final survivors per search run (100 designs).

"low-temperature" inference settings, we see a notable change in the sampling distribution, as expected. Unlike prior work [12], we do not find that our default sampling procedure reproduces the sequence distribution of ProteinMPNN on the same backbones, instead staying closer to the UniProt distribution. We find that intermediate values of $k$ yield sequence distributions between the low-temperature and full-temperature distributions, enabling user control over the tradeoff between designability and nativeness.

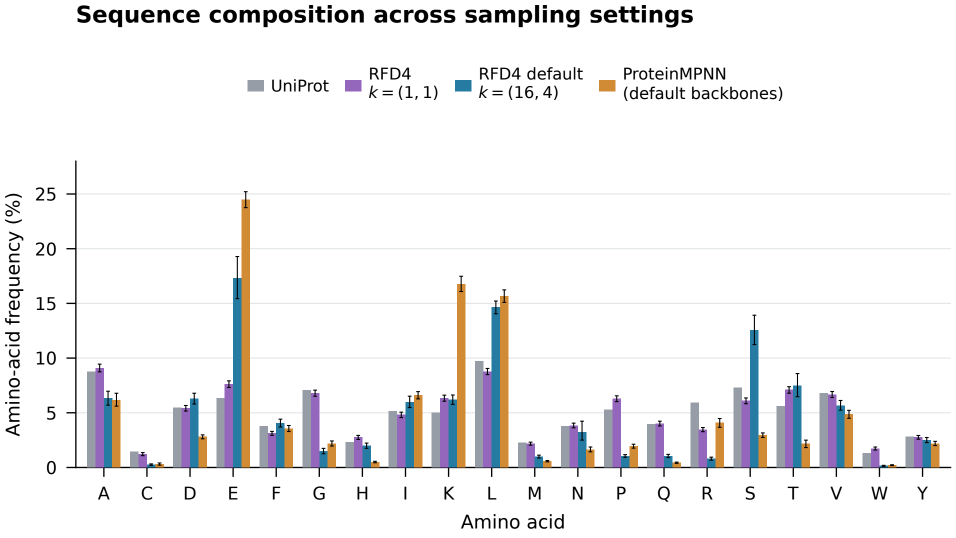

**Supplementary Fig. 6 Amino-acid composition under different RFD4-Proteina sampling settings.** Bars show, from left to right, the UniProt reference, unconditional RFD4-Proteina sequences generated with backbone and latent $k = (1, 1)$, unconditional sequences generated with the default $k = (16, 4)$, and ProteinMPNN redesigns of the RFD4-Proteina backbones generated with default sampling.

#### S5.2 Agentic design

We provide several pre-built agent skills with RFD4-Proteina, covering all aspects of a design campaign from input preparation through final production runs.

| Skill | Function |
|---|---|
| `rfd4-proteina-setup` | Installs dependencies, configures data paths, downloads checkpoints and checks the runtime environment. |
| `rfd4-proteina-inputs` | Translates design specifications into validated annotated CIF inputs. |
| `rfd4-proteina-generate` | Generates designs with specified conditioning, sampling settings and metrics. |
| `rfd4-proteina-search` | Runs reward-guided beam search with configurable branching and selection criteria. |
| `rfd4-proteina-evaluate` | Computes structural, refolding and diversity metrics and aggregates results across jobs. |
| `rfd4-proteina-visualize` | Creates molecular visualizations of final designs and generation trajectories. |
| `rfd4-proteina-autoresearch` | Runs iterative pilot experiments to compare design inputs, inference settings and search rewards. |
| `rfd4-proteina-commitment-curves` | Uses partial diffusion from a grid of restart steps to measure when design properties become determined; Figure 1e illustrates feature commitment. |
| `rfd4-proteina-campaign` | Orchestrates design campaigns and records inputs, configurations, designs, evaluations and selection decisions. |
| `rfd4-proteina-package` | Packages selected designs with metrics, selection criteria and aligned refolded structures for delivery. |

**Table S17 Core agent skills distributed with RFD4-Proteina.** The skills cover environment setup, input preparation, generation, search, evaluation, visualization, strategy exploration and campaign delivery.

### Part II: In Silico Experimentation

#### S6 Autoencoder latent-space and fine-tuning validation

We evaluated chemical relationships in the learned latent space and the effects of energy-supervised fine-tuning on generated protein structures.

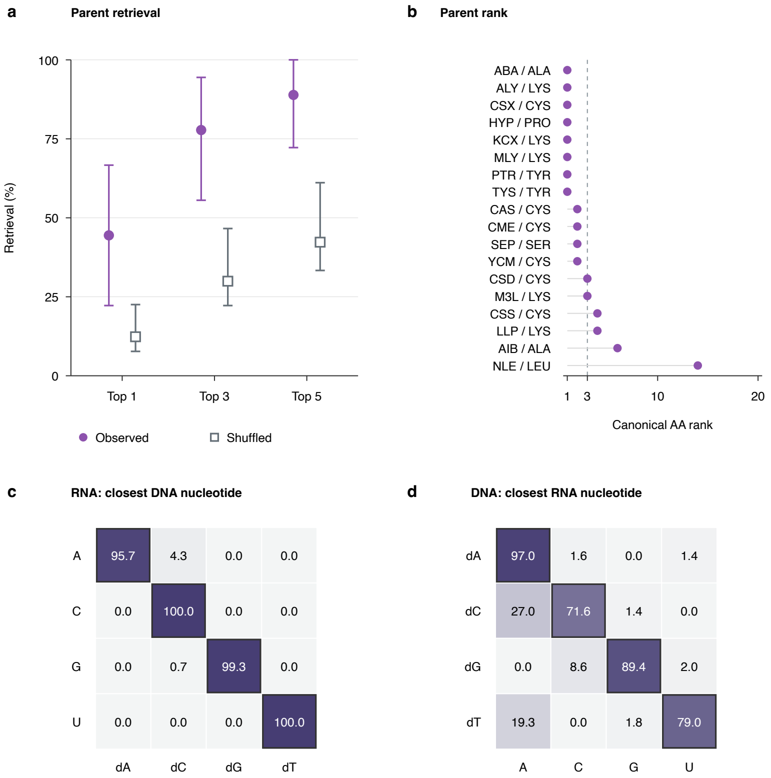

**Supplementary Fig. 7 Chemical correspondence in the latent space.** Analyses use 32-dimensional posterior-mean encodings of residues from PDB-derived structures. **a,b,** Canonical-parent retrieval for 18 modified amino acids by Wasserstein distance: top-1, -3, and -5 retrieval (a) and individual parent ranks (b). Top-1 retrieval was 44.4% versus 12.3% under frequency-preserving shuffled parent assignments. Error bars show conditional 95% bootstrap intervals. **c,d,** RNA-to-DNA (c) and DNA-to-RNA (d) nearest matches by cosine similarity. Rows show queries, columns closest matches, and cells percentages; outlines mark corresponding nucleotides. Matching rates were 98.7% and 84.3%, respectively.

##### S6.1 Evaluation in unconditional protein generation

We compared the energy-supervised autoencoder described in Section S4.3.1 with a reconstruction-only comparator using the same flow model, generation seeds and requested lengths (Supplementary Fig. 8). Each condition comprised 192 designs: 64 each at lengths of 100, 150 and 250 residues.

We scored protonated generated structures, optimized hydrogen placement with OptH in TMol, and evaluated Rosetta energies. Protein energy was divided by the number of residues and then averaged equally across designs. Energies are expressed in kcal/mol per residue. For geometry, we enumerated observed heavy-atom bonded triplets, including angles spanning peptide bonds, and compared them with the equilibrium angles in TMol's bonded-geometry parameters. For each design we computed the fraction with absolute angular deviation exceeding 5°, then averaged these fractions across designs.

Energy-supervised fine-tuning reduced mean Rosetta energy from −0.18 to −1.25 kcal/mol per residue and reduced the fraction of significant bond-angle deviations by 62%.

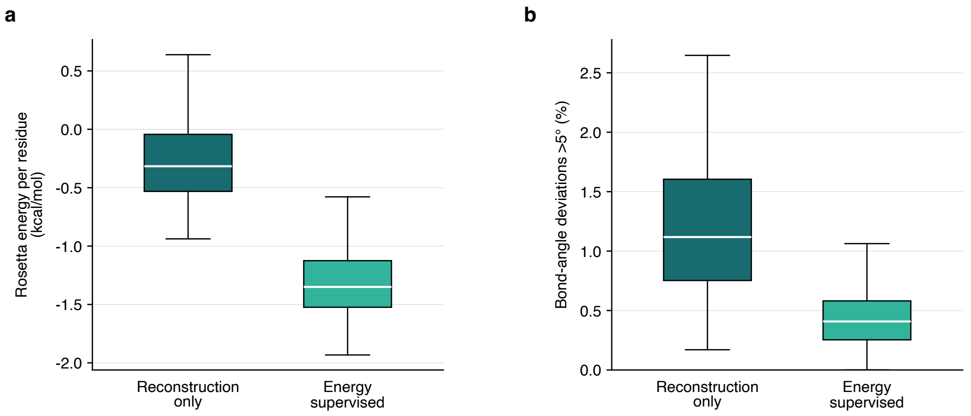

**Supplementary Fig. 8 Effect of energy-supervised fine-tuning on generated proteins. a,** Rosetta energy per residue (kcal/mol) without heavy-atom relaxation [10]. **b,** Percentage of heavy-atom bond angles deviating by $> 5°$ from TMol reference values. Each fine-tuning condition includes 192 designs (64 each at lengths of 100, 150 and 250 residues). Boxes show medians and interquartile ranges; whiskers extend to observations within 1.5 interquartile ranges. Outlier markers are omitted.

### S7 Inference-time search and recycling

#### S7.1 Search

##### S7.1.1 Time-varying search on a peptide–DNA target

We compared temporal search, constant search, and independent generation on the peptide–DNA enzyme design task described in Section 2. Final pass criteria, shared by all three strategies, are given in Table S21. Naive sampling runs every trajectory to completion and filters only on these final criteria. Temporal and constant search use the same beam-search parameters: beam width one, a branching factor of two at steps 13 and 37, and a branching factor of four at step 45 (see Tables S19 and S20 for details on rewards). Temporal search uses step-specific rewards and also allows trajectory termination if none of the branches meet the continuation criteria. Furthermore, temporal search can score either a completed rollout, or a single-step Euler prediction of the clean structure. Temporal search produced the most unique passing designs per GPU-hour (Table S18), using 24 GPU-hours per strategy. GPU cost includes generation, search and final scoring; CPU-only aggregation and structural clustering (Section S12) did not add allocated GPU-hours.

**Table S18** Search comparison on the peptide–DNA target. Passers satisfy all criteria in Table S21.

| Strategy | Completed designs evaluated | Passing designs | Unique passers | GPU hours | Unique passers per GPU-hour |
|---|---|---|---|---|---|
| Naive | **23,316** | 3 | 3 | 24.00 | 0.13 |
| Constant | 10,676 | 0 | 0 | 24.00 | 0.00 |
| Temporal | 4,057 | **35** | **9** | 24.00 | **0.38** |

**Table S19** Temporal-search rewards and continuation criteria.

| Step | Evaluation | Ranked metrics | Continuation criteria |
|---|---|---|---|
| 13 | Single-step Euler pred. | Coarse contact-pair count. | Contacts $\geq 260$ and no Foldseek match to the passer archive. |
| 37 | Single-step Euler pred. | Coarse contacts; total, DNA and peptide hydrogen-bond counts; generated motif RMSD; severe binder–target overlaps. | Contacts $\geq 260$; total H-bonds $\geq 17$; DNA H-bonds $\geq 9$; peptide H-bonds $\geq 2$; complete and correct motif mapping; motif RMSD $< 3$ Å. |
| 45 | Completed rollout and AF3 | Coarse contacts; total, DNA and peptide H-bonds; both interface min-iPAEs; AF3 motif RMSD; binder backbone RMSD; binder-aligned local DNA and peptide backbone RMSDs; binder pLDDT. | DNA min-iPAE $< 3$ and peptide min-iPAE $< 3$. |
| 60 | Completed design | Coarse contacts; total, DNA and peptide H-bonds. | None. |

Each reward sums per-metric terms, clipping every metric to a bounded range before dividing and weighting, with higher scores preferred:

$$R = \sum_i w_i \frac{\text{clip}(m_i;\, l_i, u_i)}{s_i}, \qquad \text{clip}(x; l, u) = \min(\max(x, l), u).$$

Clip bounds $[l_i, u_i]$, divisors $s_i$, and weights $w_i$ are fixed per metric; a metric absent from a given step's reward is simply omitted from the sum. The constant-search reward at every checkpoint equals the temporal step-45 reward plus two step-37-only terms (generated motif RMSD and close target-atom count), so it always scores the full metric set.

**Table S20** Constant-search rewards and continuation criteria.

| Step | Evaluation | Ranked metrics | Continuation criteria |
|---|---|---|---|
| 13, 37, 45, 60 | Completed rollout and AF3, at every checkpoint | Coarse contacts; total, DNA and peptide H-bonds; design and AF3 motif RMSD; binder-aligned local DNA and peptide backbone RMSDs; binder backbone RMSD; both interface min-iPAEs; binder pLDDT; severe binder–target overlaps. | None |

##### S7.1.2 Search under enzyme-scaffolding and ligand-burial constraints

We compared independent generation, temporal search, and constant-criterion search on six MCSA targets: M0500, M0375, M0092, M0078, M0870 and M0552. We evaluated regular enzyme-scaffolding criteria and a second definition that additionally required ligand burial (Table S25). Independent generation runs every trajectory to completion and filters only on these final criteria. At each of steps 25 and 45, search generated two branches per root and retained one candidate; surviving trajectories then completed once without further branching. See Tables S23 and S24 for details on search rewards and continuation criteria.

**Table S21** Final pass criteria shared by all three strategies.

| Structure | Criterion | Required value |
|---|---|---|
| Generated | Binder–target coarse contact pairs | $\geq 275$ |
| Generated | Total binder–target hydrogen-bond pairs | $\geq 18$ |
| Generated | Binder–DNA hydrogen-bond pairs | $\geq 9$ |
| Generated | Binder–peptide hydrogen-bond pairs | $\geq 3$ |
| Generated | Target heavy atoms within 1.5 Å of any binder heavy atom | 0 |
| Generated | Complete, correct coordinate-conditioned motif mapping | Required |
| Generated | Joint coordinate-conditioned motif RMSD | $< 3$ Å |
| AF3 | DNA-interface minimum iPAE | $< 2$ |
| AF3 | Peptide-interface minimum iPAE | $< 2$ |
| AF3 | Binder mean pLDDT | $> 80$ |
| AF3 | Binder backbone RMSD after binder-backbone alignment | $< 1.5$ Å |
| AF3 | Joint coordinate-conditioned motif RMSD | $< 3$ Å |
| AF3 | DNA residues 4–6 backbone RMSD after binder-backbone alignment | $< 3$ Å |
| AF3 | Peptide residues 3–5 backbone RMSD after binder-backbone alignment | $< 3$ Å |

A coarse contact is a binder C$\alpha$–peptide C$\alpha$ or binder C$\alpha$–DNA C4$'$ pair separated by $< 12$ Å. Each qualifying atom pair contributes one contact. Hydrogen bonds are unique interchain atom pairs identified with the AtomWorks PLIP-style interaction model. Motif mapping requires that all inserted coordinate-conditioned motifs have the expected residue/atom identities and intact reference geometry. Motif RMSD uses a single joint rigid fit of those mapped atoms to their conditioning coordinates, without atom-name symmetry permutations. Binder and local-target backbone RMSDs compare AF3 with the generated design. Local target masks retain DNA residues 4–6 and peptide residues 3–5 around the covalent linkage; distal target residues are excluded from these pass criteria.

Allocated GPU time includes calibration, generation and AF3 evaluation, but excludes CPU-only aggregation and clustering. We clustered designed protein chains separately within each target and method as described in Section S12, then counted clusters containing at least one passer. Under regular criteria, naive sampling had the highest observed yield; with burial, temporal search yielded 22.6% more passing clusters per GPU-hour than naive sampling (Table S22).

**Table S22** Passing structural clusters per GPU-hour across six MCSA targets.

| Pass definition | Method | Passing clusters | GPU hours | Clusters per GPU-hour |
|---|---|---|---|---|
| Regular | Naive best-of-$N$ | **984** | 24.00 | **41.00** |
| Regular | Temporal | 603 | 24.00 | 25.13 |
| Regular | Constant criteria | 639 | 24.00 | 26.63 |
| Burial $> 85\%$ | Naive best-of-$N$ | 177 | 24.00 | 7.38 |
| Burial $> 85\%$ | Temporal | **217** | 24.00 | **9.04** |
| Burial $> 85\%$ | Constant criteria | 165 | 24.00 | 6.88 |

***Rewards and continuation.***

Each reward clips its metrics to a bounded range before summing, so no single metric dominates once near its target ($\text{clip}(x; l, u) = \min(\max(x, l), u)$; RMSDs in Å). At step 25, search scores a clean-prediction reward favoring low designed motif RMSD, with a burial bonus when the burial criterion applies:

$$D = -\text{clip}(\text{RMSD}_{\text{motif}}, 0.5, 4)/1.5 \;+\; \underbrace{2\,\text{clip}(\text{Burial}, 0, 0.9)}_{\text{burial pass criteria only}}.$$

At step 45, and at every checkpoint for constant-criterion search, search scores a rollout-plus-AF3 reward favoring low AF3 min-iPAE, design–AF3 backbone RMSD, and AF3 motif RMSD:

$$A = -\mathrm{clip}(\mathrm{iPAE}_{\mathrm{min}}, 1.5, 8)/2 - \mathrm{clip}(\mathrm{RMSD}^{\mathrm{AF3}}_{\mathrm{bb}}, 1.5, 6)/2 - \mathrm{clip}(\mathrm{RMSD}^{\mathrm{AF3}}_{\mathrm{motif}}, 1, 4)/1.5.$$

**Table S23** Temporal-search rewards and continuation criteria for the MCSA search comparison. The final step applies Table S25 to completed outputs.

| Step | Scoring source | Reward | Continuation requirements |
|---|---|---|---|
| 25 | Clean prediction | $D$ | $\mathrm{RMSD}_{\mathrm{motif}} < 2$; also Burial $\geq 0.80$ for burial |
| 45 | Rollout + AF3 | $A$ | $\mathrm{iPAE}_{\mathrm{min}} < 3$, $\mathrm{RMSD}^{\mathrm{AF3}}_{\mathrm{motif}} < 3$, $\mathrm{RMSD}^{\mathrm{AF3}}_{\mathrm{bb}} < 3$ |

**Table S24** Constant-search rewards and continuation criteria for the MCSA search comparison. The final step applies Table S25 to completed outputs.

| Step | Scoring source | Reward | Continuation requirements |
|---|---|---|---|
| 25, 45 | Rollout + AF3 | $D + A$ | None; retain the highest reward |

**Table S25** Final pass criteria for the MCSA search comparison.

| Criterion | Regular | With burial |
|---|---|---|
| AF3 min-iPAE for the designed chain | $< 2$ | $< 2$ |
| AF3 motif all-atom RMSD | $\leq 1.5$ Å | $\leq 1.5$ Å |
| Design–AF3 backbone RMSD | $\leq 2$ Å | $\leq 2$ Å |
| Design protein–ligand clash fraction at 1.5 Å | 0 | 0 |
| AF3 burial fraction of every nonmonoatomic ligand | Not required | $> 0.85$ |

For each ligand, burial is $1 - S_{\mathrm{protein+ligand}}/S_{\mathrm{ligand}}$, where $S$ is that ligand's solvent-accessible surface area with protein present or in isolation at the same coordinates. Only protein heavy atoms contribute occlusion; other ligands are excluded. We use ProtOr radii, a 1.4 Å probe and 1,000 sphere points per atom, and take the minimum burial across ligands. Missing or invalid required metrics do not pass.

##### S7.1.3 Search under small-molecule interaction constraints

We compared independent generation with interaction-based early stopping on ten small-molecule-binding targets (Table S28). Beyond the standard small-molecule pass criteria (Section S8), we introduce a requirement that no more than one of the potential hydrogen bond and pi-stacking interactions on the small molecule can be left unfulfilled (Table S27). Independent generation completes every trajectory and applies these criteria retrospectively. Search evaluates the predicted clean structure at step 30 and stops trajectories missing more than two interactions (Table S26).

With a compute budget of 4 GPU-hours per target, search yielded 674 unique passers (16.85/GPU-h) vs 541 with naive sampling (13.53/GPU-h), a 24.6% increase (Table S28).

###### *Interaction requirements*

We assigned donor and acceptor roles using RDKit Lipinski patterns on a pH 7.4-protonated copy of each ligand and identified aromatic rings using RDKit's smallest set of smallest rings. Each assigned donor or acceptor atom received the corresponding hydrogen-bond condition; every atom of each identified aromatic ring received a $\pi$-stacking condition. Each donor role, acceptor role and aromatic ring contributes one requirement, giving 5–18 requirements per ligand.

We measured satisfaction using AtomWorks interchain interaction annotations with the PLIP-style hydrogen-bond model. Let $m$ be the number of unfulfilled requirements. The configured checkpoint reward is $-\max(m, 1)$. The final interaction threshold is applied to the completed generated structure before AF3.

**Table S26** Interaction-search scoring and continuation criteria.

| Step | Scoring source | Reward | Continuation requirements |
|---|---|---|---|
| 30 | Clean prediction | $-\max(m, 1)$ | $m \leq 2$; otherwise stop the trajectory |
| 60 | Completed design | None | $m \leq 1$ to proceed to AF3; final pass requires Table S27 |

**Table S27** Final pass criteria shared by naive sampling and interaction search.

| Criterion | Required value |
|---|---|
| Unfulfilled interactions in the generated design | $\leq 1$ |
| AF3 minimum interface PAE for the designed chain | $\leq 1.5$ Å |
| AF3 protein–ligand ipTM | $\geq 0.80$ |
| Design–AF3 protein-backbone RMSD | $\leq 1.5$ Å |
| Design–AF3 ligand RMSD after protein-backbone alignment | $\leq 5$ Å |

Stopped trajectories and skipped AF3 evaluations do not contribute passers; their allocated cost remains included in the throughput estimate.

**Table S28** Target-specific pass rates on a 4-GPU-hour budget per target and method with strict interaction criteria.

| | Naive sampling | | | Interaction search | | |
|---|---|---|---|---|---|---|
| Target | Passers | Passing clusters | Clusters/ GPU-hour | Passers | Passing clusters | Clusters/ GPU-hour |
| DFHBI | 490 | 142 | 35.50 | 861 | 193 | **48.25** |
| PLP | 111 | 59 | 14.75 | 227 | 82 | **20.50** |
| apixaban | 5 | 5 | **1.25** | 3 | 3 | 0.75 |
| bilirubin | 1 | 1 | 0.25 | 6 | 6 | **1.50** |
| chloramphenicol | 62 | 41 | 10.25 | 86 | 50 | **12.50** |
| dopamine | 352 | 134 | 33.50 | 566 | 154 | **38.50** |
| methotrexate | 1 | 1 | **0.25** | 0 | 0 | 0.00 |
| SAM | 0 | 0 | 0.00 | 0 | 0 | 0.00 |
| serotonin | 344 | 134 | 33.50 | 523 | 149 | **37.25** |
| thyroxine | 32 | 24 | 6.00 | 70 | 37 | **9.25** |
| Pooled | 1,398 | 541 | 13.53 | 2,342 | 674 | **16.85** |

##### S7.1.4 TMol-based search for point-mutant specificity

The point-mutation metric evaluates whether a designed binder preferentially interacts with a target residue over specified substitutions at that site. We use TMol for side-chain packing, local relaxation, and energy evaluation [9, 10].

For each mutation site, the metric selects a local shell including every residue with at least one atom within 8 Å of any atom at the selected site. These residues are relaxed with FastRelax using the beta2016 score function in TMol both before and after repacking of the mutation. Block-pair energies between the target and the binder are compared with and without the mutation (Equation S12). For a substitution from the wild-type residue to residue type $a$ at site $i$, the metric returns

$$\Delta E_{i,a} = E_{i,a}^{\mathrm{mut}} - E_{i}^{\mathrm{WT}}, \tag{S12}$$

together with both component energies. Positive values indicate less favorable interactions after mutation and therefore favor recognition of the wild-type residue.

#### S7.2 Time-varying recycling

At integration step $k$, we perform $R_k$ trunk evaluations at the same noisy coordinates and latent state before applying the sampler update (Section S3.1.4). For a trajectory of $K$ integration steps, the total number of trunk evaluations is $N_{\mathrm{trunk}} = \sum_{k=1}^{K} R_k$. Changing $R_k$ therefore changes the internal refinement budget without changing the integration grid. The benchmark schedules use 60 steps, with allocations that depend on the task and thinking level (Supplementary Fig. 9). Protein binding uses early-weighted schedules; small-molecule binding and enzyme scaffolding use middle-peaked schedules. DNA binding uses early-weighted schedules at low and medium thinking and uniform allocation at high thinking. Total pass counts and benchmark protocols are specified in Section S8.

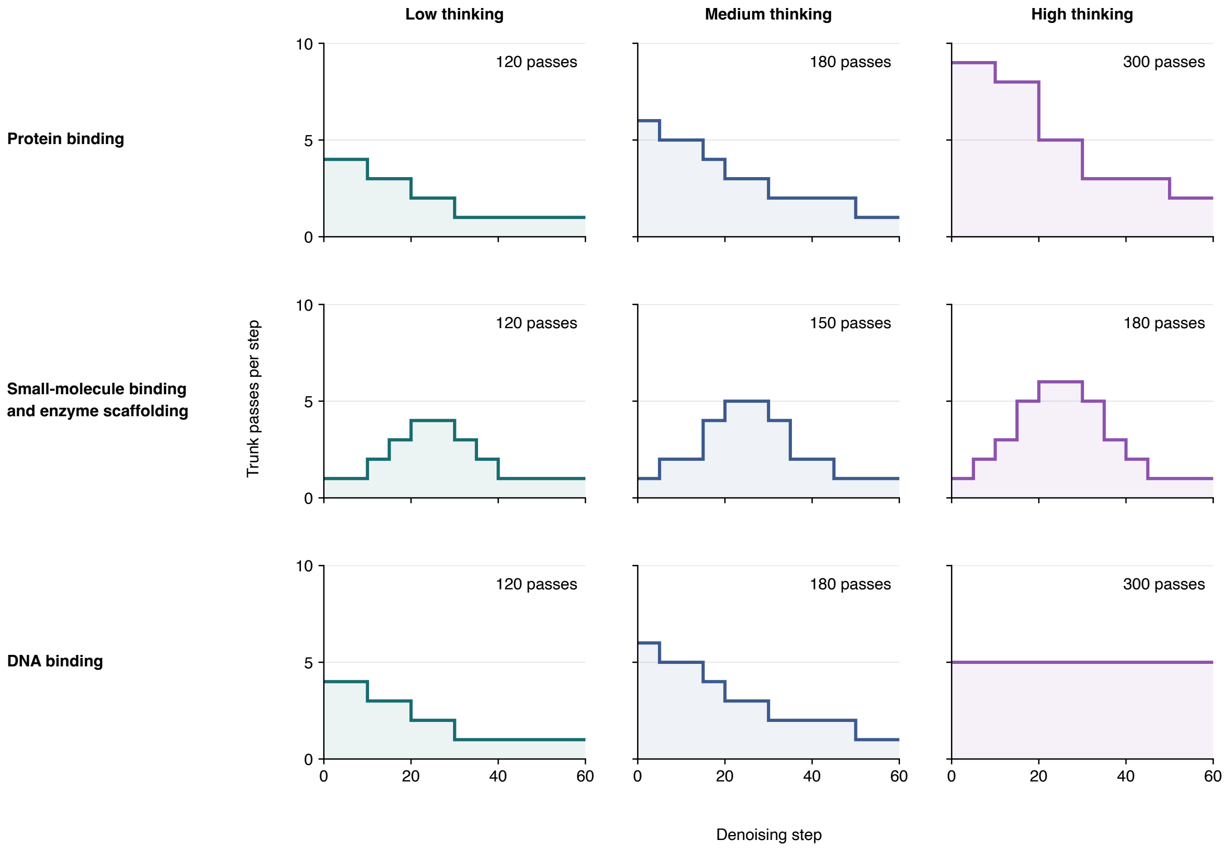

**Supplementary Fig. 9 Task-specific recycling schedules.** Columns show low, medium and high thinking; rows show the benchmark tasks. Curves give trunk passes per denoising step, including the initial pass; numbers give totals over 60 steps. The horizontal axis indexes denoising steps.

##### S7.2.1 Recycling-window ablation on the peptide–DNA task

We localized the effects of additional trunk evaluations using the same peptide–DNA enzyme design task described in Section 2. For each replicate, we compared two generations with the same input and random seed: one with a single forward pass at each integration step, and one with additional recycles in each of the six 10-step windows across the trajectory. Each of these 24 conditions and the shared one-pass baseline contained 5,000 paired replicates (Supplementary Fig. 10).

Each design was evaluated with AF3 twice with different model seeds, both with five AF3 recycles; coordinate-conditioned protein regions were supplied as templates. We averaged the two AF3 measurements within each design, and we pooled individual within-replicate changes from baseline. RMSD metrics could only be calculated on 4,435–4,590 designs per condition because of occasional invalid motif mappings. The plotted motif RMSD fits and scores eight enzyme catalytic atoms, after requiring complete, correct-identity mapping; phosphotyrosine phosphate atoms and Mn are excluded from this RMSD. Contact molecular surface (CMS) and the supplementary figure's hydrogen-bond pair counts compare the protein with the combined DNA/peptide substrate on the design and AF3 structures. We calculated CMS as $\sum_i a_i \exp(-0.5 d_i^2)$ over buried substrate molecular-surface elements, where $a_i$ is the element area in Å$^2$ and $d_i$ is its distance in Å to the nearest buried protein surface point. Larger CMS values indicate more extensive, closely packed interfaces. Binder backbone RMSD compares AF3 with the design after binder-backbone alignment. The interface hydrogen-bond quantity reported for Figure 1g is instead a TMol-calculated [10] interchain hydrogen-bond energy evaluated on the AF3 structure. Figure 1g shows absolute means for the 20–30 window, with error bars of one standard error of the mean across the population of per-design mean AF3 values.

##### S7.2.2 Recycling-schedule comparison at fixed compute

We asked whether the best allocation of recurrent refinement depends on the design task. At a fixed budget of 120 trunk passes over 60 generation steps, nonuniform allocation gave higher clustered pass rates than uniform allocation for all three tasks tested (Supplementary Fig. 11). Middle-peaked schedules performed best for enzyme scaffolding and small-molecule binding, whereas early-weighted and middle-peaked schedules tied for protein binding. The largest gain was in enzyme scaffolding, where middle-peaked allocation increased the clustered pass rate from 8.5% to 10.9%.

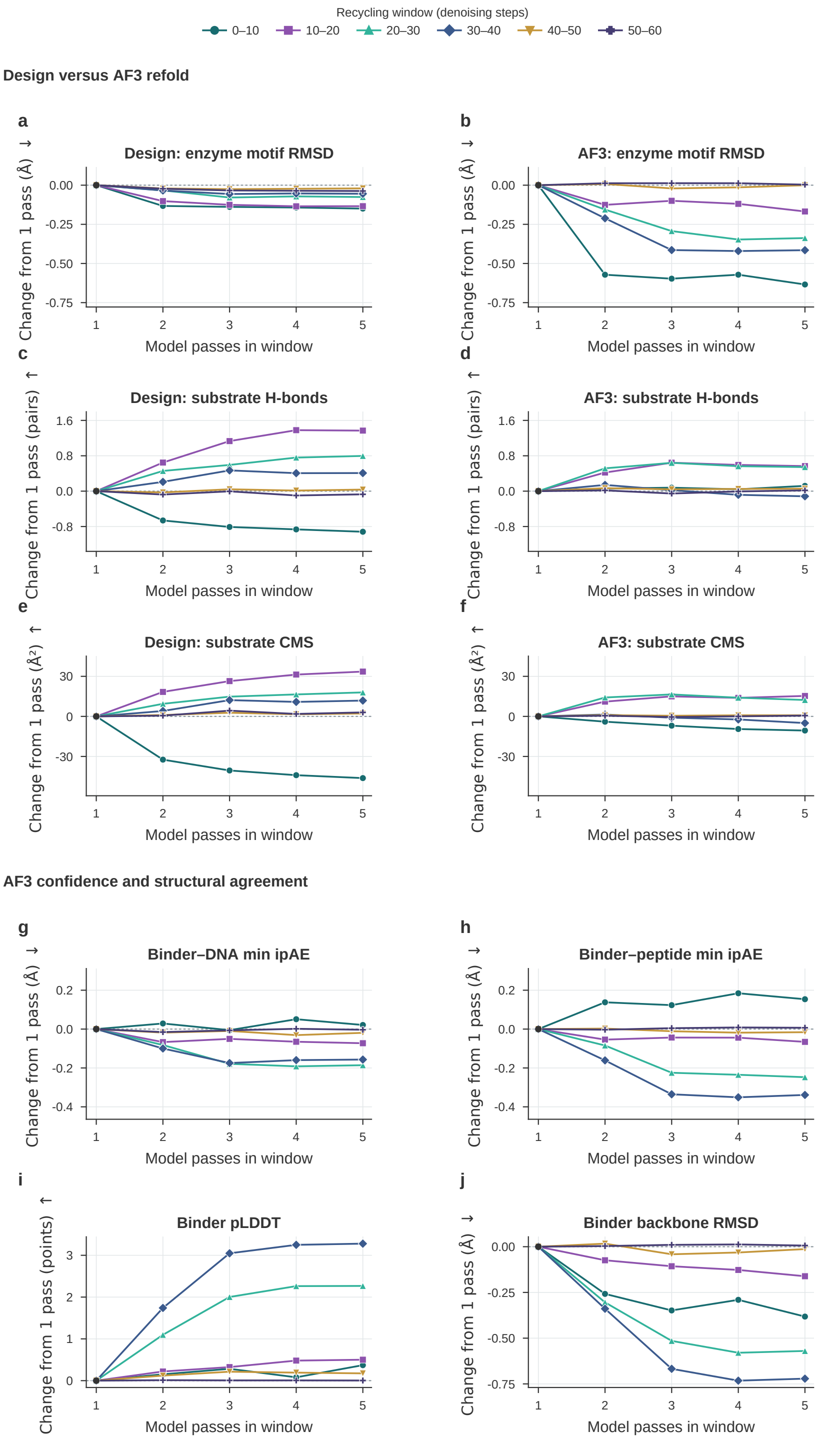

**Supplementary Fig. 10 Recycling effects depend on denoising window and metric in the peptide-DNA enzyme design task.** Mean paired changes from a shared one-pass baseline across 5,000 paired replicates per condition. **a**–**f**, Enzyme motif RMSD, substrate hydrogen bonds, and contact molecular surface (CMS) calculated across the designed structure and the AF3 refolded structure at each condition. **g**–**j**, AF3 DNA/peptide min-ipAE, binder pLDDT and binder backbone RMSD to the designed structure.

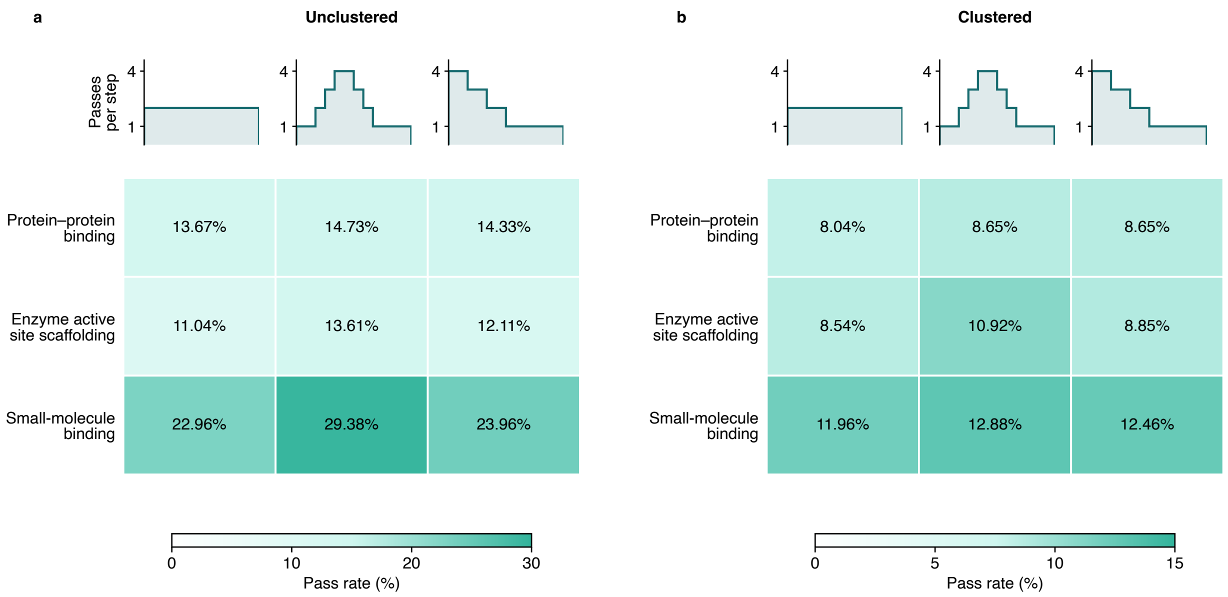

**Supplementary Fig. 11 Recurrent-pass allocation at fixed low-thinking compute. a,b,** Unclustered (a) and clustered (b) pass rates. Columns compare uniform, middle-peaked, and front-loaded schedules, each using 120 total trunk passes over 60 generation steps; diagrams show passes per step. Rows show protein–protein binding (24 targets), enzyme active-site scaffolding (37 targets), and small-molecule binding. For computational efficiency, the latter uses a smaller 12-example subset of the DISCO benchmark [22]. Each target has 200 designs per schedule. Cells show percentages, with separate color scales for the two panels. Clustered rates count distinct passing Foldseek clusters per attempted design (TM-score threshold 0.5; clustering within each target and schedule). Failed attempts remain in the denominators.

### S8 Benchmarks

#### S8.1 Evaluation cohorts and metrics

We evaluated protein binding on 24 targets collated from the Proteina-Complexa and RFdiffusion3 benchmarks, small-molecule binding on a 77-target subset of the DISCO benchmark, enzyme scaffolding on 37 Atomic Motif Enzyme (AME) tasks, and DNA binding on the three targets from the RFdiffusion3 preprint [21] (Fig. 2). Target-level results for protein binding, small-molecule binding, DNA binding, and AME are shown in Supplementary Figs. 18, 21, 19, and 20, respectively. Detailed, interactive breakdowns alongside complete protocols for each model are available on the RFD4-Proteina website and the accompanying archive.

Each model and protocol was evaluated with 200 generation attempts per target. Failed attempts and outputs with unmatched required motifs remain in the denominator. Designs were assessed using a common AlphaFold 3 (AF3) refolding procedure and the structural and confidence criteria below. Ordinary pass rate is the number of passing designs divided by 200; clustered pass rate is the number of distinct structural clusters among passing designs divided by 200, clustered as described in Section S12. Mean pass rates include all targets, including those with no passing designs.

#### S8.2 Success criteria and benchmark adaptations

**Protein binding.** Following previous methods [1, 6], passing designs require target-aligned binder backbone RMSD $\leq 2.5$ Å, binder pTM $\geq 0.80$, and minimum interface PAE $\leq 1.5$ Å. The target may not change sequence during the course of generation; any deviation from input conditioning is considered a failure (as can happen with codesign models that diffuse within a latent space).

**Small-molecule binding.** We used the 77 small-molecule tasks with published comparative results in DISCO Fig. 2c [22], excluding DNA and RNA entries because nucleic-acid binding is evaluated separately. Following previous methods [6, 69], passing designs require protein-backbone RMSD $\leq 1.5$ Å, backbone-aligned ligand RMSD $\leq 5$ Å, protein-small molecule ipTM $\geq 0.80$, and protein-small molecule minimum interface PAE $\leq 1.5$ Å. We note that these criteria are stricter than the original DISCO criteria, which use protein-backbone RMSD $\leq 2$ Å and maximum ligand-centroid displacement $\leq 2$ Å after backbone alignment, without consideration of any confidence metrics. We also show complete benchmark results for all models under the original DISCO criteria on the RFD4-Proteina website and the accompanying archive.

**Enzyme scaffolding.** Of the 41 AME tasks, we excluded three covalent protein–ligand adducts (M0054_1qfe, M0711_2esd and M0731_1mt5) requiring unindexed modified-residue support and preservation of residue–ligand bonds, and M0024_1nzy because four ligand phosphate atoms lacked experimental coordinates. Passing designs require backbone RMSD $\leq 2$ Å, motif all-atom RMSD $\leq 1.5$ Å, and minimum interface PAE $< 1.5$ Å. We modified the AME success criteria [2] by adding a minimum interface PAE threshold of $< 1.5$ Å and removing the clash filter to improve consistency across benchmarks and incorporate structure-prediction confidence at the interface, which are common practice for enzyme design campaigns. Motif RMSD is evaluated over the mapped, coordinate-conditioned protein atoms; unmatched required motifs fail. For models that do not predict the optimal insertion, a greedy matching algorithm was used to align the predicted motif with the required motif.

**DNA binding.** We used the three DNA targets (7M5W, 7N5U and 7RTE) from the RFdiffusion3 preprint [21]. Related methods for generative design of sequence-specific DNA-binding proteins are described by Sehgal et al. [98]. Passing designs require DNA-aligned binder backbone RMSD $\leq 8$ Å and minimum interface PAE $\leq 1.5$ Å. Alignment uses all target DNA atoms for both fixed- and flexible-DNA designs. We increased the designed binder length to 120–150 residues relative to the original benchmark to match the target length range of ongoing DNA-binding protein design campaigns.

#### S8.3 Baseline protocols

The main-figure BindCraft and BindCraft2 comparisons use a single-backbone protocol: one backbone-optimization trajectory followed by one ProteinMPNN sequence, with continuation gates disabled and no subsequent folding, relaxation, filtering, or ranking. These protocols differ from the full design pipelines, which include additional internal evaluation and selection. In the accompanying website and archive, full results for all pipelines—including internal evaluation and selection—are provided; the single-backbone protocols achieved higher pass rates under common AF3 evaluation than the full pipelines. We do not show BindCraft and BindCraft2 full-pipeline results in the main figure to avoid clutter and maintain focus on the single-backbone comparison.

Supplementary Fig. 16 evaluates the full BindCraft and BindCraft2 pipelines. Each full-pipeline attempt contributes its designated terminal output to common AF3 evaluation, including available outputs rejected by internal filters; internal folding, sequence design, selection, and rejected proposals contribute to generation cost.

The main-figure Genie3 comparison uses one generative draw followed by one CA-MPNN sequence, without search or ColabFold evaluation. Genie3 search is evaluated separately in Supplementary Fig. 16. In the enzyme benchmark, ODesign receives additional observed motif coordinates. AlphaProtein-novo codesign and its LigandMPNN-redesigned variant share the same generated scaffolds.

For fixed-DNA RFD3, we followed the procedure described by Sehgal et al. [98], using AF3-predicted target duplexes, a six-base-pair ORI window, donor/acceptor conditioning, and 2.5 Å ORI jitter. Each target has 200 attempts at protein lengths of 120–150 residues.

For RFD4-Proteina, all three thinking levels use 60 denoising steps, with task-specific schedules for recurrent refinement. Protein binding uses 120, 180 and 300 total trunk passes for low, medium and high thinking, respectively, concentrated toward the beginning of generation. Small-molecule binding and enzyme scaffolding use 120, 150 and 180 passes, concentrated at intermediate steps. DNA binding uses the same low and medium schedules as protein binding, but five passes at every step for high thinking (300 total). These counts include the first pass at each step. The schedules below give passes per step in successive blocks of five denoising steps.

| Schedule | Thinking | Passes per step (12 successive five-step blocks) |
|---|---|---|
| Early | Low | 4, 4, 3, 3, 2, 2, 1, 1, 1, 1, 1, 1 |
| Early | Medium | 6, 5, 5, 4, 3, 3, 2, 2, 2, 2, 1, 1 |
| Early | High | 9, 9, 8, 8, 5, 5, 3, 3, 3, 3, 2, 2 |
| Intermediate | Low | 1, 1, 2, 3, 4, 4, 3, 2, 1, 1, 1, 1 |
| Intermediate | Medium | 1, 2, 2, 4, 5, 5, 4, 2, 2, 1, 1, 1 |
| Intermediate | High | 1, 2, 3, 5, 6, 6, 5, 3, 2, 1, 1, 1 |
| Uniform (DNA) | High | 5, 5, 5, 5, 5, 5, 5, 5, 5, 5, 5, 5 |

For each thinking level, Figure 2 reports one conditioning protocol across the full target set: backbone-only target conditioning for protein binding, generated ligand conformations for small-molecule binding, fixed ligand coordinates for enzyme scaffolding, and generated DNA conformations for DNA binding. For protein binding, the conditioning mode is selected by its mean clustered pass rate across all targets, rather than separately for each target.

#### S8.4 Coordinate conditioning

Supplementary Fig. 12 compares more and less coordinate conditioning for high-thinking RFD4-Proteina and RFD3 on the same target sets. Protein binding compares all-atom target conditioning with backbone-only conditioning; small-molecule binding compares fixed with generated ligand conformations; AME compares fully with partially fixed ligands while retaining catalytic coordinate constraints; and DNA binding compares fixed with generated DNA conformations. Points are arithmetic means over all targets, including those with no passing designs. Connecting lines join the two condition means.

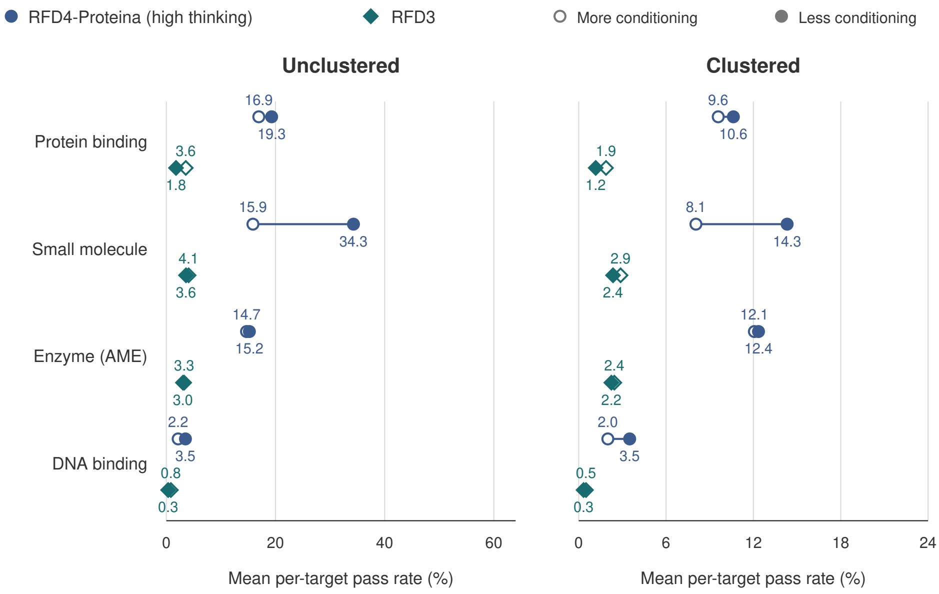

**Supplementary Fig. 12 Coordinate conditioning and design success.** Ordinary (left) and clustered (right) pass rates for high-thinking RFD4-Proteina (blue circles) and RFD3 (teal diamonds), averaged over the same benchmark targets as Fig. 2, including zero-success targets. Open symbols denote more coordinate conditioning and filled symbols less coordinate conditioning. Lines connect the two condition means. Each target has 200 original attempts. Ordinary pass rate is passing attempts divided by 200; clustered pass rate is passing backbone clusters divided by 200. PPI compares all-atom with backbone-only target coordinates ($n = 24$); AME fully with partially fixed ligand coordinates ($n = 37$); small molecules fixed with unfixed ligands ($n = 77$); DNA fixed with unfixed DNA coordinates ($n = 3$). Small-molecule results use the binder success criteria from the main benchmark. AME retains catalytic coordinate constraints in both conditions. See Supplementary Section S8 for protocols and evaluation criteria.

##### S8.4.1 Reducing binder–target structural similarity

We observed that some generated binders closely resembled their targets. For PD-L1, the frequency of target-like designs increased with binder length, reaching 80% in the longest length bin (Supplementary Fig. 13a,b). This behavior was not specific to RFD4-Proteina: across the PPI benchmark, target-like structures occurred in 8.8% of BoltzGen designs, 12.2% of Proteina-Complexa designs, and 11.1% of high-thinking RFD4-Proteina designs under all-atom target conditioning (Supplementary Fig. 13c). The predominance of homomeric complexes in structural databases provides a plausible source of this shared bias [99].

We next tested internal-coordinate conditioning, which the model encountered during training but not specifically for protein-binder design. Compared with absolute-coordinate conditioning, this approach reduced the target-like fraction from 9.4% to 2.3% with backbone-only conditioning and from 11.1% to 1.6% with all-atom conditioning (Supplementary Fig. 13d). Across the full PPI benchmark, the corresponding clustered pass rates reached 11.77% and 12.15%, respectively, compared with 10.63% and 9.58% under absolute-coordinate conditioning. These results suggest that changing how target geometry is specified can reduce target resemblance while improving design success. We therefore recommend backbone-only internal-coordinate conditioning as the default for protein-binder design because it reduces target-like outputs while leaving target side-chain conformations free to change.

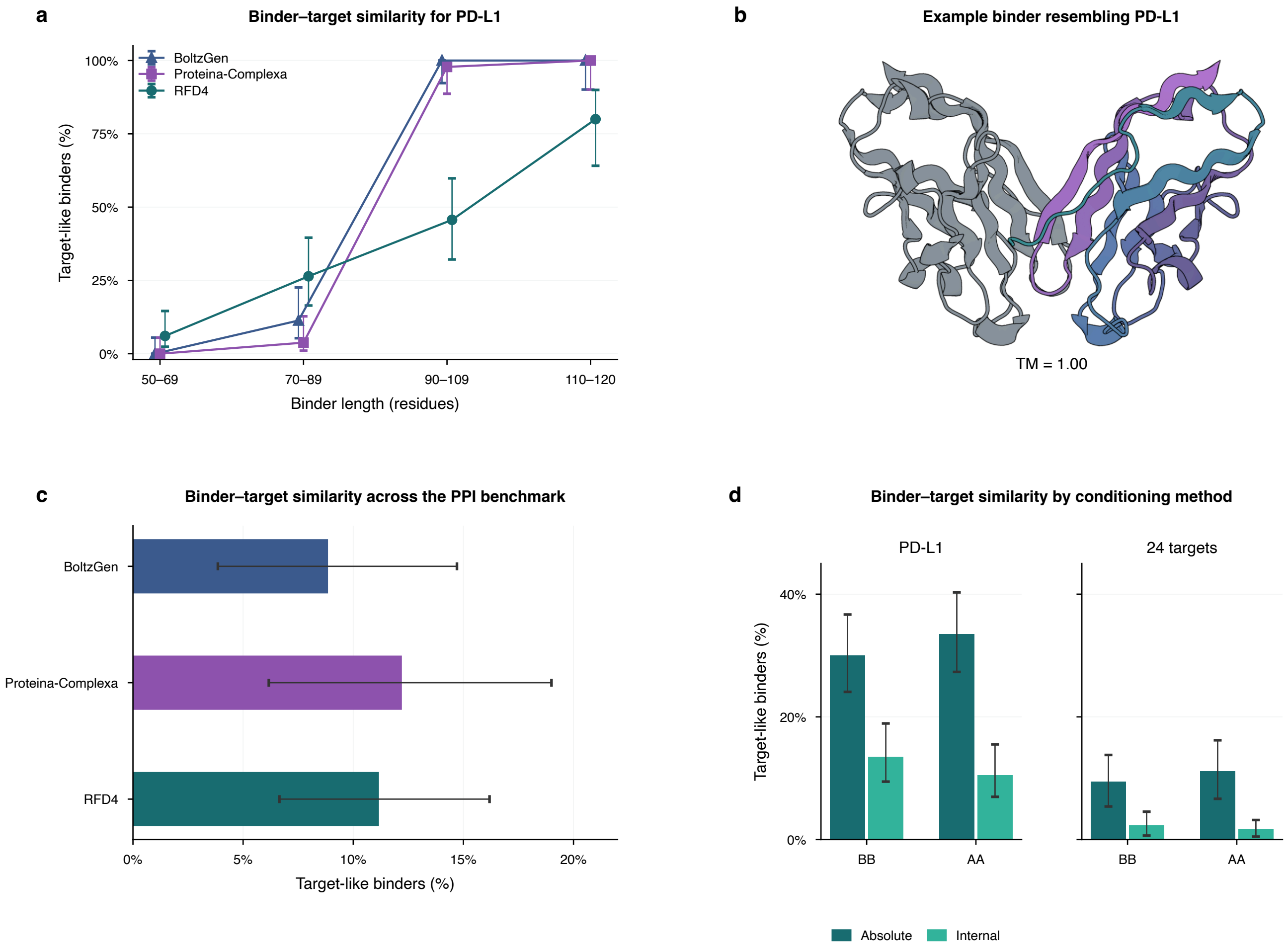

**Supplementary Fig. 13 Structural similarity between designed binders and their targets.** Target-like binders have a binder-normalized C$\alpha$ TM-score $\geq$ 0.8 with $\geq$ 90% coverage. **a,** Target-like binder frequency by length for PD-L1 (200 designs per model). **b,** Example binder (colored) resembling PD-L1 (gray). **c,** Mean frequencies across 24 protein-binding targets (200 designs per target and model). **d,** High-thinking RFD4-Proteina with absolute- or internal-coordinate conditioning of target backbone (BB) or all atoms (AA). Error bars show 95% Wilson intervals for PD-L1 and target-bootstrap intervals for benchmark means.

#### S8.5 Extrinsic search evaluation

RFD4-Proteina search uses AF2-Multimer for protein binding and OpenFold3 [100] for small-molecule binding, AME, and DNA binding. Each target has 200 initial attempts, including attempts that stop early or fail. Search and its structure-predictor calls contribute to generation cost. Common AF3 refolding is evaluated separately (Table S29).

We find that search increased clustered pass rates across all four tasks. However, due to the efficiency of RFD4-Proteina standard generation, search with the basic benchmark criteria reduced passing-cluster yield per GPU-hour.

#### S8.6 GPU-normalized yield and budgeted discovery

Generation cost includes sequence design, internal structure prediction, search, and selection. Batched GPU costs are divided among all attempts in the batch. We measure generation throughput in attempts per GPU-hour, excluding common AF3 evaluation.

| Task | Inference | Clustered pass rate (%) | Clusters/GPU-h generation only | Clusters/GPU-h including AF3 |
|---|---|---|---|---|
| Protein binding | High thinking | 10.6 | 120.7 | 66.5 |
| | Search | 14.9 | 8.8 | 8.4 |
| Small-molecule binding | High thinking | 14.3 | 787.4 | 237.1 |
| | Search | 20.9 | 23.0 | 22.0 |
| AME scaffolding | High thinking | 12.1 | 453.4 | 155.2 |
| | Search | 16.6 | 14.0 | 13.5 |
| DNA binding | High thinking | 3.5 | 140.6 | 52.3 |
| | Search | 4.8 | 5.4 | 5.1 |

**Table S29 Structure-predictor-guided search on the standard benchmarks.** Each setting uses 200 attempts per target across 24 protein-binding, 77 small-molecule-binding, 37 AME, and three DNA-binding targets.

GPU-normalized yield is computed separately for each target as

$$Y_i = \frac{3600\,C_i}{G_i + F_i},$$

where $C_i$ is the number of passing clusters and $G_i$ and $F_i$ are total generation and common AF3 evaluation costs in GPU-seconds. Main-figure efficiency bars show the arithmetic mean of $Y_i$ across targets. Supplementary efficiency panels use the same per-GPU-hour normalization for passing designs or clusters, with the corresponding success count in the numerator. Failed attempts and their consumed GPU time are included. GPU costs exclude setup, loading, compilation, CPU scoring, and idle allocations.

For the compute-budget analysis (Supplementary Fig. 16), we divided the 24 protein-binding targets into 12 easy and 12 hard targets based on the seven evaluated methods. For each of 100 paired random orderings of the 200 attempts per target, sampled without replacement, we retained the longest sequence of completed attempts within each GPU budget and reclustered passing designs within that subset.

#### S8.7 Accelerated oracle evaluation

Repeated structure prediction can dominate the cost of oracle-guided search. Our AlphaFold 2, AlphaFold 3, and OpenFold3 interfaces [14, 19, 100] use persistent workers that retain loaded models between requests. Binaries compiled at an appropriate padding length are loaded automatically and persist across runs. Featurized model inputs are prefetched from the set of folding tasks, ensuring that the GPU remains fully utilized. The runtimes support optimized attention kernels, including Pallas attention for AF2.

### S9 Benchmark result figures

Protocols, success criteria and cost accounting for the figures below are described in Section S8.

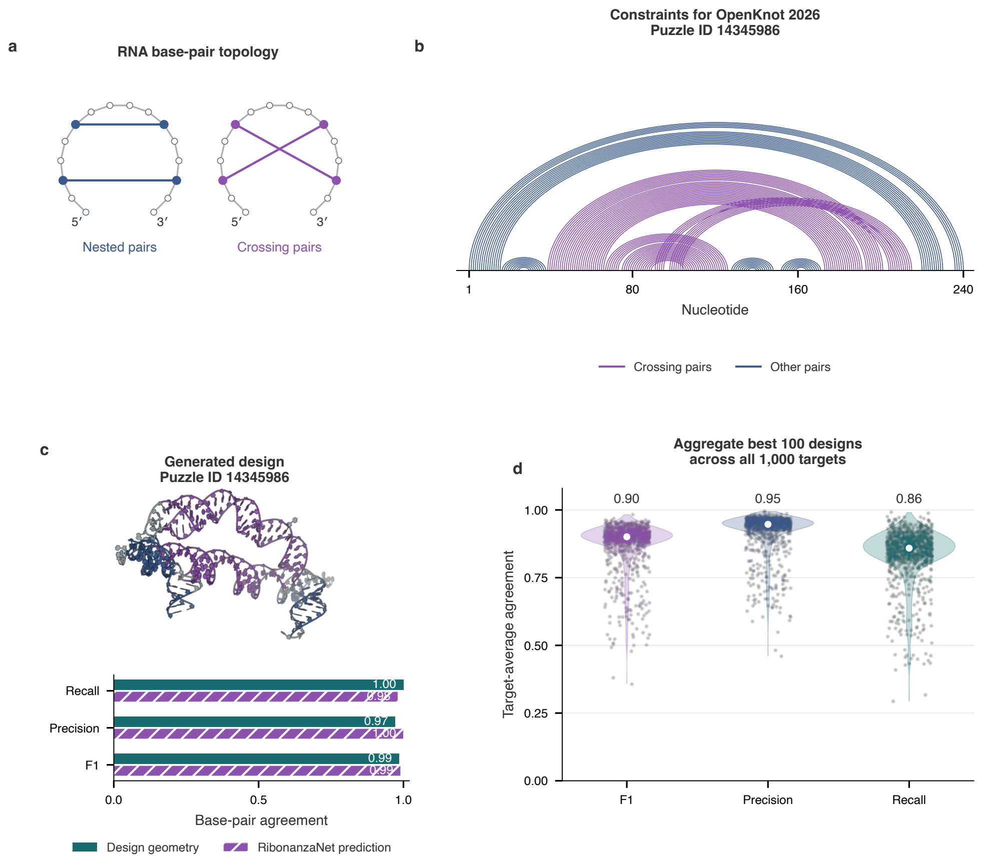

**Supplementary Fig. 14  RNA pseudoknot design. a,** Nested and crossing base-pair topologies. **b,** Pairing constraints for a 240-nucleotide OpenKnot target; purple marks crossing pairs and blue other pairs. **c,** Design for the target in panel **b** and agreement with its requested pairs, assessed from design geometry and RibonanzaNet sequence-based predictions [101]. **d,** Target-mean prediction agreement for the top designs (up to 100 per target) across 1,000 targets. Points represent targets; white markers and lines show medians and interquartile ranges.

**Task coverage**

| Model / checkpoint | Protein design: Protein binding | Protein design: Cyclic peptide | Protein design: Antibody design | Small-molecule binding: Fixed | Small-molecule binding: Unfixed | Enzyme scaffolding: Fully fixed | Enzyme scaffolding: Partially fixed | DNA binding: Fixed | DNA binding: Unfixed | RNA: Design |
|---|---|---|---|---|---|---|---|---|---|---|
| **RFD4-Proteina** | ✓ | ✓ | ✓ | ✓ | ✓ | ✓ | ✓ | ✓ | ✓ | ✓ |
| RFdiffusion3<br>Protein-design checkpoint | ✓ | – | – | ✓ | ✓ | ✓ | ✓ | – | – | – |
| RFdiffusion3<br>DNA-binder checkpoint | – | – | – | – | – | – | – | ✓ | ✓ | – |
| RFdiffusion3NA<br>Nucleic-acid generation | – | – | – | – | – | – | – | – | – | ✓ |
| RFDpoly | – | – | – | – | – | – | – | – | – | ✓ |
| BoltzGen | ✓ | ✓ | ✓ | ✓ | ✓ | – | – | ✓ | ✓ | – |
| Proteina-Complexa | ✓ | – | – | ✓ | – | ✓ | – | – | – | – |
| DISCO | – | – | – | – | ✓ | – | – | – | ✓ | – |
| ODesign<br>Protein-generation variants | ✓ | ✓ | – | ✓ | ✓ | ✓ | – | ✓ | – | – |
| ODesign<br>Nucleic-acid checkpoint | – | – | – | – | – | – | – | – | – | ✓ |
| AlphaProtein-novo | – | – | – | – | – | ✓ | – | – | – | – |
| RFdiffusion2 | – | – | – | – | – | ✓ | – | – | – | – |
| Genie3 | ✓ | – | – | – | – | – | – | – | – | – |
| RFdiffusion | ✓ | – | – | – | – | – | – | – | – | – |
| Protein-Hunter | ✓ | ✓ | – | – | ✓ | – | – | – | ✓ | – |
| BindCraft | ✓ | – | – | – | – | – | – | – | – | – |
| BindCraft2 | ✓ | ✓ | ✓ | – | – | – | – | – | – | – |
| ESMFold2 inversion | ✓ | – | ✓ | – | – | – | – | – | – | – |

✓ Supported task    – Not documented

**Supplementary Fig. 15 Task coverage across biomolecular design methods.** Checkmarks indicate documented or demonstrated task support, independently of inclusion in our benchmarks; dashes indicate that support remains unestablished in the reviewed sources. RFdiffusion3 protein-design and DNA-binder checkpoints and the separate RFdiffusion3NA model are shown separately, as are ODesign protein and nucleic-acid variants [6, 98, 102, 103]. Other rows may combine task-specific settings or adapters. Fixed and unfixed refer to ligand or DNA coordinate conditioning under each method's native semantics; enzyme columns distinguish fully and partially fixed ligand coordinates while retaining catalytic constraints. RNA design denotes generation of RNA, not protein design against a supplied RNA sequence [104]. Antibody design includes scaffolded CDR and single-domain workflows. Cyclic-peptide workflows need not support identical closure chemistries [13, 15, 105].

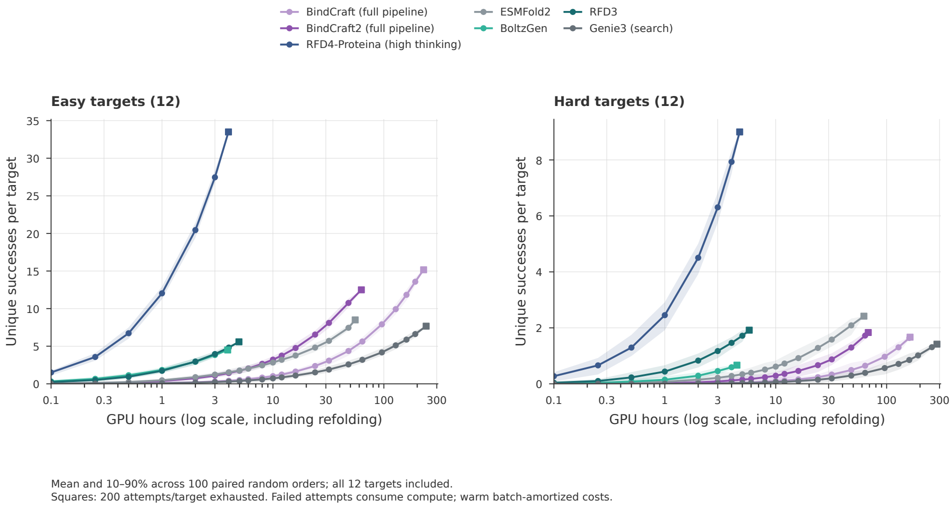

**Supplementary Fig. 16 Protein-binder discovery across compute budgets.** Mean passing clusters per target versus total GPU time across each easy or hard 12-target group, including generation and AF3 evaluation. Shading indicates the 10th–90th percentiles across attempt orderings; squares mark completion of all 200 attempts per target. Sampling and reclustering procedures are described in Supplementary Section S8.

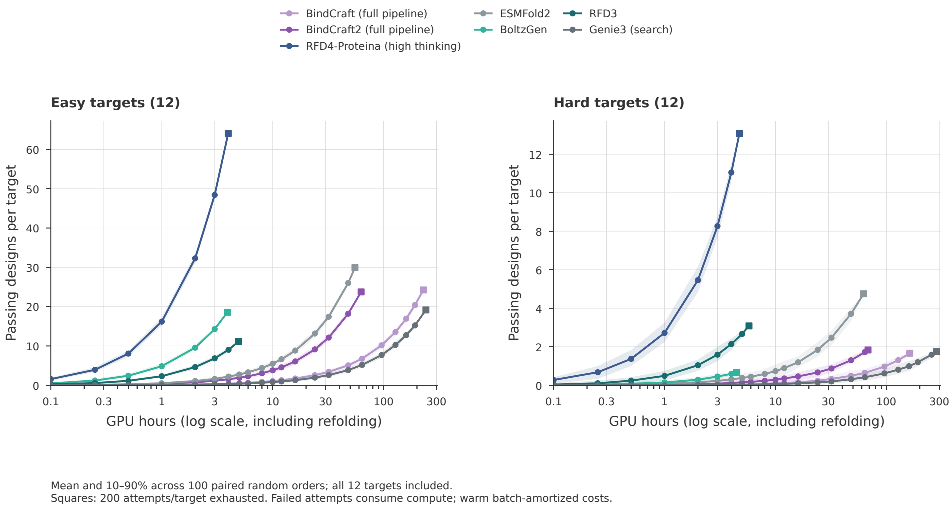

**Supplementary Fig. 17 Passing protein-binder designs across compute budgets.** Mean passing designs per target, before clustering, using the same attempts, easy/hard target split, generation-plus-AF3 costs and 100 random orderings as Supplementary Fig. 16. Shading denotes the 10th–90th percentiles across orderings; squares mark exhaustion of the recorded attempts.

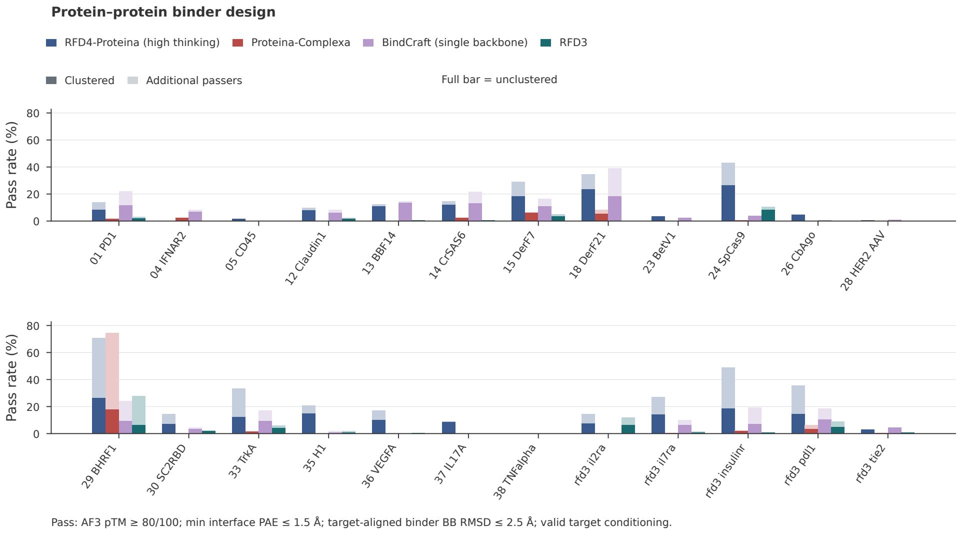

**Supplementary Fig. 18 Protein-binder design success by target.** All 24 targets are shown for high-thinking RFD4-Proteina, Proteina-Complexa, single-backbone BindCraft, and RFD3. Dark bars show distinct passing clusters divided by 200 original attempts; full bar heights show all passing designs divided by 200. Light extensions show additional passing designs within those structural clusters. Failures remain in the denominator. Success criteria and clustering follow Supplementary Section S8.

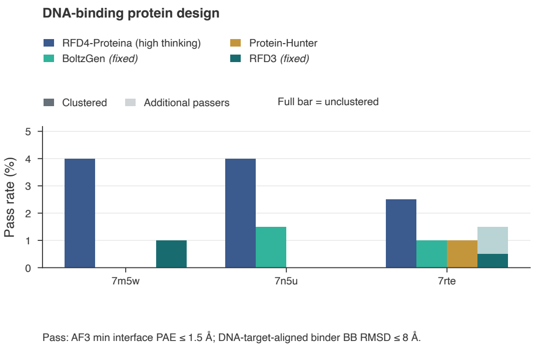

**Supplementary Fig. 19 DNA-binder design success by target.** All three targets from the RFdiffusion3 preprint [21] are shown for high-thinking RFD4-Proteina, fixed-DNA BoltzGen, Protein-Hunter, and fixed-DNA RFD3. Dark bars show clustered pass rates; full bars show unclustered pass rates, each out of 200 original attempts per target. DNA conformations are generated unless labeled fixed. Success criteria and clustering follow Supplementary Section S8.

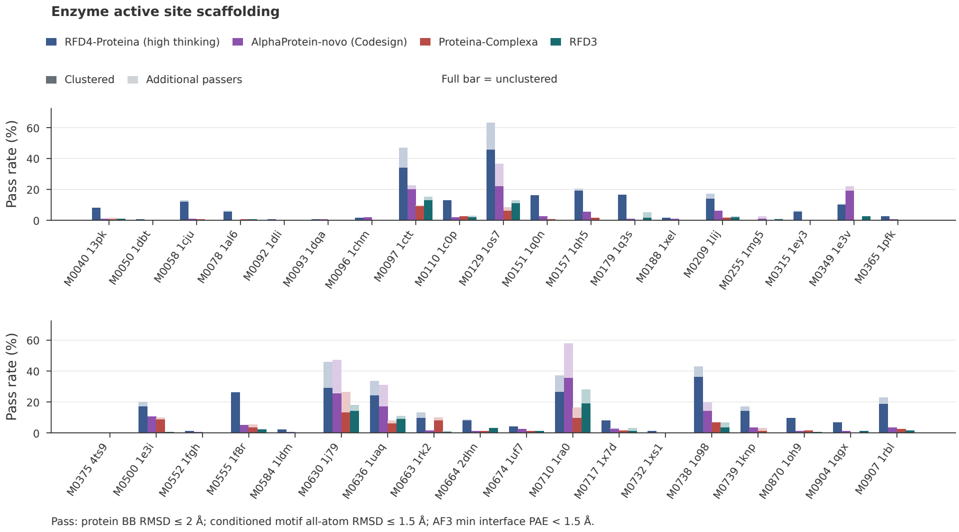

**Supplementary Fig. 20 Enzyme-scaffolding success by target.** All 37 AME tasks (MCSA target identifiers) are shown for high-thinking RFD4-Proteina, AlphaProtein-novo codesign, Proteina-Complexa, and RFdiffusion3. Ligand coordinates are fully fixed. Dark bars show clustered pass rates; full bars show unclustered pass rates, each out of 200 original attempts per task. Success criteria and clustering follow Supplementary Section S8.

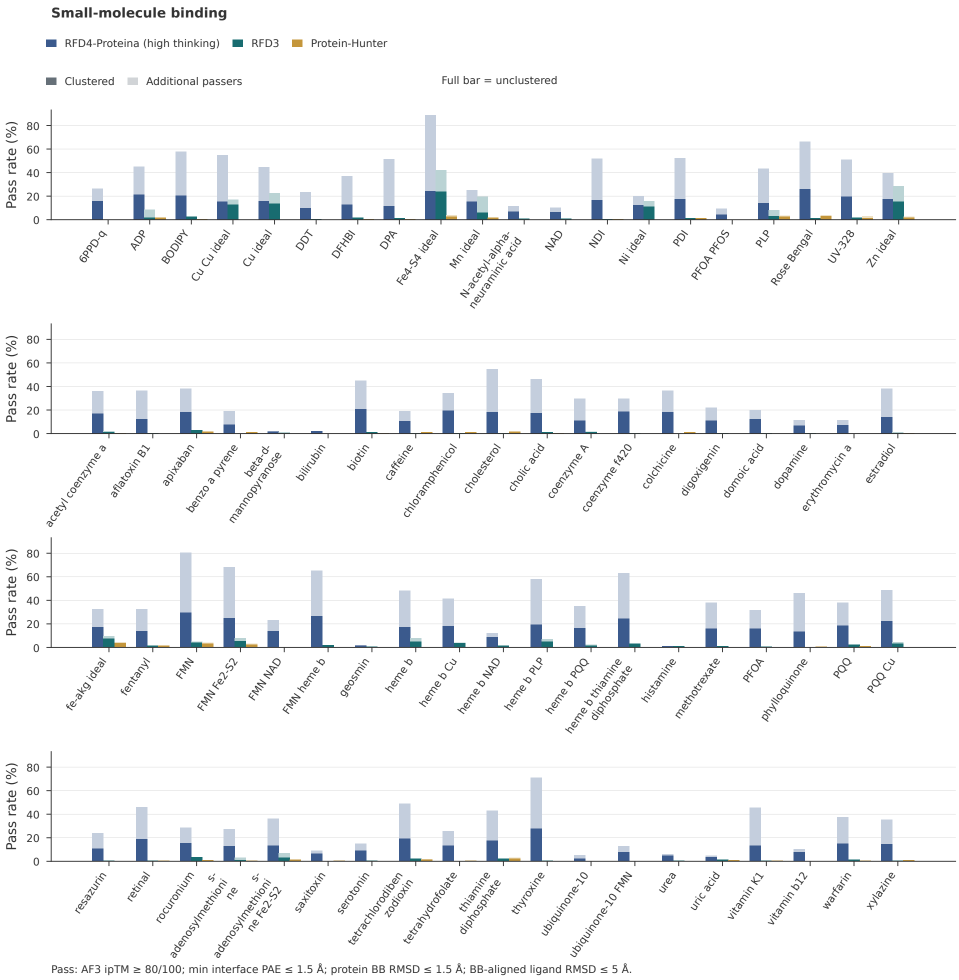

**Supplementary Fig. 21 Small-molecule-binder design success by target.** All 77 targets are shown using the small-molecule binder success criteria in Section S8. Dark bars show clustered pass rates; full bars show unclustered pass rates, each out of 200 original attempts. Complete benchmark results, including the original DISCO criterion, are available on the RFD4-Proteina website.

### S10 Molecular-workflow evaluations

We evaluated six molecular workflows using task-specific sampling settings (Fig. 3).

#### S10.1 Sidechain packing

We used PackBench [25]: 137 single-chain CASP14 (66) and CASP15 (71) targets of fewer than 2,000 residues, on their native backbones. We kept the 114 targets for which every method returned valid structures with the input sequence.

All methods sampled eight structures per target and setting, given the protein sequence and backbone coordinates. RFD4-Proteina conditioned on backbone coordinates (and non-standard residues) and on the sequence of every protein residue, with sidechain noise scales of 0.1, 1, 2, 3, 5, 7, 10 and 15. Comparators were LigandMPNN (v1.1.1, `ligand_mpnn`, all residues fixed and repacked without native-sidechain context, temperature 0.1), FlowPacker, FAMPNN and PIPPack.

Quality is the all-atom MolProbity score (cctbx; lower is better). Diversity is the mean of the 28 pairwise sidechain RMSDs between the eight samples of a target (heavy atoms present in both, after superposition on the input backbone N, C$\alpha$, C and O). Values are averaged within targets, then across targets, with target-level standard errors. All methods were scored by the same pipeline, and each setting was sampled once.

The protein-only illustration shows one of eight samples generated for 1ENH with sequence and backbone conditioning and noise scale 0.9. For the 1FMN RNA–FMN complex, we conditioned molecular identities and the RNA sugar–phosphate backbone while sampling RNA bases and the unfixed ligand. We generated six samples at each noise scale (0.3 and 0.7), restored the conditioned terminal OP3 atom, and refined geometry with the backbone constrained and bases and ligand unrestrained. A geometry-checked sample is shown. These selected overlays illustrate conformational changes; ensemble variance and retained binding affinity were not evaluated.

#### S10.2 Inverse folding

We sampled eight sequences per backbone and setting on two backbone sets and evaluated them by AF3 refolding. The native set comprises 156 single-chain PDB backbones shared by RFD4-Proteina and LigandMPNN, taken from the PDB validation split (deleaked against training): chains of 50–500 residues with standard amino acids and no unresolved gaps, one chain per 30% sequence-identity cluster (1,248 sequences per setting). The enzyme set comprises ten backbones sampled by RFdiffusion3 (RFD3) [6] for each of 34 M-CSA catalytic motifs of the AME benchmark (340 backbones, 2,720 sequences per setting). For each motif, we took the first ten of 100 sampled backbones that met the following criteria: motif residues and ligands matching the benchmark, motif tip and ligand atoms within 1.5 Å RMSD of it, and a continuous chain (consecutive C$\alpha$–C$\alpha$ distances of 3.4 to 4.2 Å). Three of the 37 motifs were excluded because LigandMPNN designs for them could not be refolded.

RFD4-Proteina was run by partial diffusion from the input backbone ($t = 0.99$) with the sequence latents at $t = 0$ and 60 steps of the temporal-score-rescaled sampler, where the backbone is effectively fixed (C$\alpha$ RMSD to the input below 0.4 Å). On the enzyme set, all atoms of the motif residues and ligands were conditioned and the remaining residues were designed. Diversity was tuned by the scale of the sampling noise: 12 values from 0 to 5.0 on the native set, without additional recycling, and nine values from 0 to 15 on the enzyme set, with the default recycling schedule. LigandMPNN (v1.1.1, `ligand_mpnn` model with side-chain context) designed eight sequences per backbone at 12 temperatures from 0.01 to 5.0 (native set) and seven from 0.01 to 1.0 (enzyme set), with X and C excluded from the alphabet, ligand atoms taken from the input and, on the enzyme set, the catalytic residues fixed.

Each sequence was refolded together with its ligands by single-sequence AF3 with five recycles and one diffusion sample, without templates. Backbone RMSD is taken over N, C$\alpha$, C and O atoms between the refold and the designed structure; motif RMSD is the all-atom RMSD of the conditioned motif residues against the input coordinates, with symmetric side-chain atoms permuted to their best assignment. A native design passed at backbone RMSD $\leq$ 1.5 Å, and an enzyme design at backbone RMSD $\leq$ 2.0 Å and motif RMSD $\leq$ 1.5 Å.

The pass rate of a setting is the fraction of its sequences that passed. Diversity of a backbone is one minus the mean pairwise sequence identity of its eight sequences (28 pairs), and that of a setting the mean over backbones. Paired differences were formed per backbone, pairing each RFD4-Proteina setting with the LigandMPNN temperature of the closest mean diversity, with 95% percentile intervals from 10,000 bootstrap resamples of backbones. RFD4-Proteina noise scales 10 and 15 reach diversities (0.71 and 0.84) beyond those of LigandMPNN (up to 0.62) and are not compared. Each setting was sampled and refolded once.

#### S10.3 Flexible-backbone interface design

We evaluated 127 intended pairs in 13 HBDesigner families [28]. Fixed and flexible RFD4-Proteina each generated 32 proposals per pair from LigandMPNN complexes using partial diffusion (fraction 0.5) and TMol-guided beam search. Sequence changes were allowed within 8 Å of the interface for fixed RFD4-Proteina and 12 Å for flexible RFD4-Proteina; backbone flexibility was restricted to 8 Å. Selection used intended-complex physical and polar-network scores, without AF3 or off-target scores; recipes were selected on six development families. All 1,249 A/B pairings per method were evaluated with three single-sequence AF3 seeds, five recycles and no templates. We averaged seeds, then calculated intended-minus-off-target ipTM within each family and averaged families equally; intervals use 100,000 family-bootstrap resamples. Fixed and flexible recipes also differ in recycling and scoring, so their comparison does not isolate backbone flexibility. This measures predicted partner separation; the flexible-RFD4-Proteina versus HBDesigner interval includes zero.

#### S10.4 Ligand stereochemistry

Methods received annotated ligands and a 100-residue expandable protein segment. We analyzed 43 ligands with outputs from every method: ten outputs per ligand, except Protein Hunter (139 total). RFD4-Proteina used medium thinking. Stereochemical adherence is the fraction of correct eligible chiral centers or E/Z bonds, averaged over outputs and then equally over ligands. PoseBusters 0.6.5 assessed isolated ligand geometry, excluding connectivity, InChI-conversion and energy tests. Failures remain in the denominator. AF3 references refold all 430 RFD4-Proteina outputs with five recycles, one sample, single sequences and no templates; no outputs were filtered for stereochemical correctness. Intervals use 100,000 paired ligand-bootstrap resamples. Generation budgets differ between pipelines.

#### S10.5 Ensemble-based activity ranking

We compared seven highly active retroaldolases ($k_{\mathrm{cat}}/K_M > 10^3$) with 399 inactive designs using RFD4-Proteina and PLACER [26]. RFD4-Proteina conditioned sequence and backbone and sampled the covalent intermediate, with sidechain noise 1.0. Each intermediate was scored using 16 conformers per design; PLACER's 16 were selected from 50 generated conformers ranked by predicted RMSD. Negative intermediate sidechain-atom spread after backbone alignment supplies the activity-ranking score. AUROC intervals use 2,000 bootstrap resamples; paired differences use 10,000 stratified design resamples. The main-panel intermediate was selected retrospectively. Both intermediates and the broader 144-active cohort remain in Supplementary Fig. 22; paired intervals do not establish a method advantage.

#### S10.6 Binder rescue

Of 146 published non-binding RFdiffusion designs, 102 failed the common AF3 filter: chain-of-interest pTM $\geq$ 0.80, minimum interface PAE $\leq$ 1.5 Å and target-aligned binder backbone RMSD $\leq$ 2.5 Å. RFD4-Proteina used six partial-diffusion attempts (fraction 0.9) per parent, compared with the first six HalluDesign outputs and six RFOptimization cycles [30, 31]. RFD4-Proteina and HalluDesign started from RFdiffusion backbones; RFOptimization used sequence-bearing AF2 complexes. A parent is rescued if any attempt passes all filters; all 612 attempts per method contribute to cost. Efficiency is rescued parents per recorded design plus AF3 GPU-hour; timing overhead is not allocated identically across methods. Rescue fractions use Wilson 95% intervals. These outcomes are computational, not experimental binding measurements.

### S11 Retroaldolase ensemble-based activity ranking

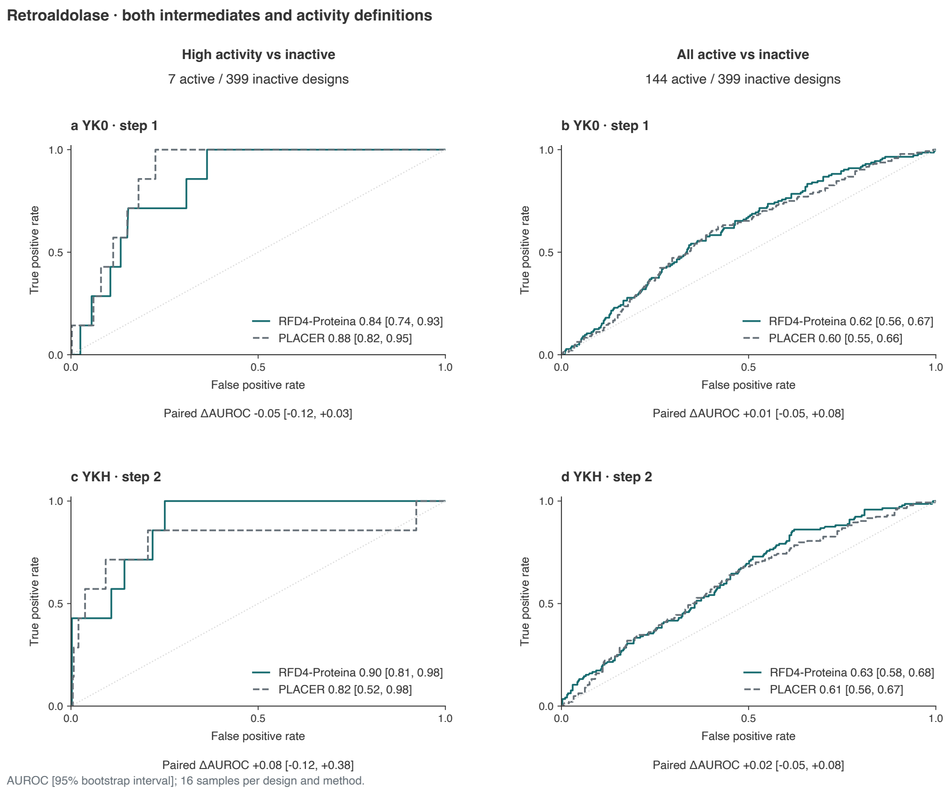

**Supplementary Fig. 22 Retroaldolase activity ranking from active-site ensemble spread.** Receiver-operating-characteristic curves for both reaction intermediates compare seven designs with $k_{\mathrm{cat}}/K_M > 10^3$ against 399 inactive designs (left) and all 144 active designs against the same 399 inactive designs (right). RFD4-Proteina and PLACER use identical cohorts and 16-member ensembles in each comparison. Smaller ensemble spread is scored as more active. Intervals are percentile bootstrap intervals. Figure 3e shows YKH, selected after examining both intermediates. The paired RFD4-Proteina-minus-PLACER AUROC differences are −0.05 (95% interval, −0.12 to 0.03) for YK0 and 0.08 (−0.12 to 0.38) for YKH, from 10,000 stratified paired-design bootstrap resamples. With the broader active/inactive split, the corresponding differences are 0.01 (−0.05 to 0.08) for YK0 and 0.02 (−0.05 to 0.08) for YKH. None of the paired intervals establishes a method advantage.

### S12 Structural clustering of designs

We use structural clustering to distinguish distinct passing designs from redundant restatements of the same solution. We report the number of passing clusters as "unique passers" and divide it by the number of original attempts to obtain the "clustered pass rate". Unless a section states otherwise, clustering is run separately within each target/task and model/protocol/strategy, on designed-chain structures only, using Foldseek `easy-cluster` [82] (version `10-941cd33`) with `alignment-type=1` (TM-align), `cov-mode=0` (bidirectional coverage), `min-seq-id=0` (no sequence-identity prefilter) and a TM-score threshold of 0.5, without an additional explicit coverage cut-off. Each distinct cluster containing at least one passing design contributes one unique passer, regardless of how many passing designs it contains.

### Part III: Wet-Lab Experimentation

### S13 Experimental methods

#### S13.1 Calcium-dependent calmodulin binders

Cell-free expression and BLI characterization were performed by Adaptyv.

##### S13.1.1 Cell-free expression of protein ligands

DNA constructs encoding the ligands, each fused to a C-terminal assay tag, were designed by reverse-translating the target protein sequences. The sequences were optimized for manufacturability and yield using codon optimization algorithms to maximize expression efficiency. The optimized DNA constructs of the variants and the 3′ fragment containing the linker and affinity tags were ordered as Gene Fragments from Twist Bioscience.

Constructs were assembled using the NEBuilder HiFi DNA Assembly Kit (NEB) in 2 µL reactions. The assembled products were characterized by capillary electrophoresis (Agilent ZAG DNA Analyzer, ZAG-135-5000 Kit, FA/ZAG 96-Capillary Array Short, 33 cm), and their concentrations were measured using the Qubit DNA Quantification Kit (Invitrogen). Ligand expression was carried out in 8 µL reactions using an optimized prokaryotic in vitro translation system and 4 nM of assembled gene fragment. Reactions were incubated at 37 °C for 8 hours. Soluble yields were measured by GFP11/GFP1–10 complementation against a standard curve, using the molecular weight of the tagged construct. The quantitative range was 2.5–25 µM; values outside this range were reported as below or above the quantification limits.

##### S13.1.2 Affinity characterization by biolayer interferometry (BLI)

Kinetic binding measurements were performed on a BLI instrument (Gator Bio) using Strep-Tactin XT probes to capture Twin-Strep-tagged ligands.

**Sensor preparation and capture.**

Probes were equilibrated in running buffer before use. Ligand capture consisted of a first baseline of 60 s, ligand loading for 120 s, and a second baseline of 180 s (300 s in later holo runs). The median loading shift was 3.99 nm across both experiments, including subsequently excluded replicates.

The apo running buffer was 10 mM HEPES, 150 mM NaCl, 3 mM EDTA and 0.2% Tween-20, pH 7.4. Holo buffer omitted EDTA and contained 2 mM $CaCl_2$. The appropriate buffer was used throughout capture, baseline, association and dissociation. Because EDTA also chelates magnesium, the comparison distinguishes calcium-loaded from divalent-depleted conditions. Temperature was held at 25 °C and data were sampled at 5 Hz.

**Association and dissociation (multi-cycle kinetics).**

Human calmodulin (CALM1) was the soluble analyte, and the Twin-Strep-tagged designed binder was immobilized. The primary analyte series was 31.6, 100, 316.2 and 1,000 nM, plus a 0 nM blank. Each concentration was measured in a separate cycle on the same sensor, with ligand reloaded before each cycle. Each cycle consisted of association for 300 s followed by dissociation for 300 s. Cycles were quality-controlled independently; rejected cycles were omitted without imputing missing concentrations. Selected holo designs were remeasured. Figure 4a shows the replicate-1 response curves for the illustrated design.

Between cycles, probes were regenerated in 10 mM glycine-HCl, pH 1.5, applied 5 times for 10 s each, followed by neutralization in running buffer. Negative controls (buffer-only and a non-binding ligand) were included for reference subtraction and drift correction.

**Data processing and model fitting.**

Data were processed with Adaptyv fitting software. All sensorgrams underwent standardized preprocessing: trimming to the relevant phases (baseline, association, dissociation), correcting signal jumps at phase transitions, aligning the association and dissociation phases, and subtracting baseline and reference signals.

Each replicate was fitted across the concentration series: its association and dissociation curves were fitted simultaneously, with $k_{on}$ and $k_{off}$ shared across concentrations and $R_{max}$ fitted per cycle to account for differences in ligand loading. The default model was a 1:1 Langmuir interaction, and $K_D$ was calculated as $k_{off}$ / $k_{on}$, with 95% confidence intervals for each parameter. Alternative binding models (bivalent analyte and conformational change) were also fitted. Where one described the curves better than 1:1, it was selected and its apparent $K_D$ reported.

Candidate fits were filtered on quality before selection. A fit was kept only if its residual error in both the association and dissociation phases, normalized to signal size, stayed below a fixed threshold and bulk refractive-index jumps were small relative to the binding signal. The remaining fits were ranked by residual error, then penalized when a rate constant sat at the edge of what the assay can resolve, then ranked by confidence-interval width.

Ligands were classified as binders or non-binders based on the presence of quantifiable binding curves and calculated $K_D$ values. Where a ligand produced a significant signal shift during the association phase ($\geq$300% over the negative control) but the signal could not be reliably fit, the binding label was assigned based on the magnitude of the observed shift. We report overall holo affinity estimates from 1:1 fits, combining independent replicates. For the two lowest apparent $K_D$ estimates, $k_{off}$ reached the assay's lower resolvable limit of $10^{-5}$ $s^{-1}$ under the 300-s dissociation conditions. The affinity estimates and confidence intervals are conditional on this limit and do not capture the unresolved uncertainty in $k_{off}$.

#### S13.2 Peptide–MHC binders

##### S13.2.1 DNA library preparation

For yeast surface display, all designed protein sequences were first padded at both termini using GS to a uniform length of 102 amino acids for a 350 bp oligonucleotide library. Then the sequences were reverse translated using dnachisel while codon optimizing for S. cerevisiae. DNA sequences were ordered as oligonucleotide libraries via Twist Biosciences 350 bp sizes.

##### S13.2.2 Yeast surface display screening with FACS

Transformed S. cerevisiae EBY100 strain library cultures were grown in C-Trp-Ura (2% glucose w/v) medium and induced for expression in SGCAA (0.2% glucose w/v) medium. Cells were washed with PBSF (PBS with 1% BSA w/v) and incubated with FITC-conjugated anti-C-Myc chicken antibody (ICL, CMYC-45F) for expression sorting. For binding sorts, cells were additionally incubated with peptide-MHC biotinylated tetramer (Fred Hutchinson Cancer Center Immune Monitoring Services) conjugated with phycoerythrin (PE) or allophycocyanin (APC). Cells were incubated for 30 minutes, then washed and resuspended with PBSF. All sorts were performed on the Sony SH800 FACS instrument. FACS data was analyzed using the FlowJo software.

##### S13.2.3 Yeast surface display clonal binding assay

Clonal yeast surface display samples were obtained by plating sorted pools from FACS on glucoseTrp-Ura agar plates prepared in-house, incubating at 30 °C for 48 hours and picking single colonies. Cultures were grown in C-Trp-Ura (2% glucose w/v) medium and induced for expression in SGCAA (0.2% glucose w/v) medium for 12–18 hours. 50 µL of culture was transferred to a 96-well plate and washed with PBSF. Cells were stained at 1:100 v/v with FITC-conjugated anti-C-Myc chicken antibody (ICL, CMYC-45F) for expression and with 1:50 to 1:102400 v/v dilutions of peptide-MHC biotinylated tetramer (Fred Hutchinson Cancer Center Immune Monitoring Services, PE- or APC-conjugated) for on-target titration curves, or at a single concentration for specificity assessment. Cells were incubated for 30 minutes and then washed and resuspended in PBSF. All flow cytometry experiments were performed on the Attune NxT instrument and data was analyzed using the FlowJo software.

#### S13.3 Intrinsically disordered region binders

##### S13.3.1 Gene synthesis and cloning

Genes encoding the designed binders and target proteins were optimized for bacterial expression, ordered as synthetic double-stranded DNA fragments and assembled by Golden Gate cloning into the LM0627 expression vector, which confers kanamycin resistance and encodes a C-terminal $His_6$ tag.

##### S13.3.2 Target constructs

Each IDR epitope was expressed as a fusion protein with an N-terminal AviTag (GLNDIFEAQKIEWHE), a solubilizing GB1 or SUMO domain, and a C-terminal $His_6$ tag: M–SGGG–AviTag–linker–(GB1 or SUMO)–linker–IDR–GSGSHHWGSTHHHHHH.

##### S13.3.3 Protein expression and purification

Constructs were transformed into E. coli BL21(DE3), grown overnight in LB containing kanamycin, and used to inoculate Terrific Broth autoinduction medium supplemented with kanamycin, 2 mM $MgSO_4$ and 1× 5052 autoinduction mixture. Cultures were grown for approximately 21 h at 37 °C. Cells were harvested by centrifugation and lysed with BugBuster reagent supplemented with DNase I (0.01 mg $ml^{-1}$) and protease inhibitors. Clarified lysates were incubated with nickel-charged resin, which was washed with 25 mM Tris-HCl (pH 8.0), 300 mM NaCl and 30 mM imidazole, and bound protein was eluted in the same buffer containing 400 mM imidazole. Eluates were then purified by size-exclusion chromatography (SEC) on an HPLC system with a Superdex 75 Increase 5/150 GL column (Cytiva) equilibrated in TBS (25 mM Tris-HCl, pH 8.0, 150 mM NaCl). Fractions were collected into 384-well plates, and for each sample the fraction at the peak nearest the expected elution volume of the intended construct was selected. Concentrations were calculated from the $A_{280}$ signal using extinction coefficients derived from each construct's sequence. Selected fractions were diluted in TBS to 10 $\mu$M with an Opentrons OT-2 liquid handler.

##### S13.3.4 Target biotinylation

Targets were biotinylated in vitro at the AviTag lysine using the BirA500 biotin ligase kit (Avidity) according to the manufacturer's instructions and repurified by SEC as above to remove free biotin and enzyme.

##### S13.3.5 Surface plasmon resonance

Experiments were performed at 25 °C on a Biacore 8K (Cytiva) in HBS-EP+ running buffer. Biotinylated targets were captured on Series S Sensor Chip CAP chips with the Biotin CAPture Kit (Cytiva).

**Individual binder screening.** Binders were measured in serial dilution series, with concentrations shown in Supplementary Fig. 25. Analytes were injected in single-cycle runs without regeneration between injections, and the surface was regenerated between cycles. Data were fitted globally to a 1:1 Langmuir interaction model using Biacore Insight Evaluation Software to determine $K_D$ from the fitted association and dissociation rate constants.

**All-by-all specificity screen.** To assess specificity, binder hits were tested against all their target peptides as single-cycle series at 8, 40, 200 and 1,000 nM, with each pair measured once. Responses were normalized to target capture level (response ÷ captured RU × 100). The summary figure shows, for each target, the binder with the highest capture-normalized on-target response at 1,000 nM as well as its normalized off-target responses.

#### S13.4 Trk receptor binders

##### S13.4.1 Bio-layer Interferometry (BLI)

Commercially available tagged proteins for hTrkA (Recombinant Human TrkA Protein (His & hFc Tag), HPLC-verified, Sino Biological, 11073-H03H), mTrkA (Recombinant Mouse TrkA Fc Chimera Protein, CF, R&D Systems, 10235-TK), hTrkB (Human TrkB / NTRK2 Protein, Fc Tag, Acro Biosystems, NT2-H5254), hTrkC (Human TrkC / NTRK3 Protein, Fc Tag (MALS verified), Acro Biosystems, TRC-H5256) were used for binding assays. The BLI experiments were performed on an OctetRED96 BLI system (ForteBio) at room temperature in HBS-EP buffer (Cytiva Life Sciences). Prior to measurements, ProA-coated biosensors were first equilibrated for at least 10 min in the assay buffer. Fc-tagged target ligands were immobilized onto the biosensors by dipping them into a solution with 10 µg/mL protein until the loading signal reached 0.5–1 nm. Association and dissociation of the analytes were monitored by dipping biosensors in the solutions containing analytes at various concentrations for 60 s when screening and 120–180 s when measuring affinity followed by dipping the biosensors in fresh buffer for the same duration. Experiments were performed at 22 °C while shaking. Global kinetic or steady-state fits were performed on bare-tip-subtracted data assuming a 1:1 binding model.

##### S13.4.2 Surface Plasmon Resonance (SPR)

Commercially available Fc-tagged hTrkA and mTrkA proteins described above were used for binding assays. SPR experiments were performed on a Biacore 8K instrument (Cytiva) in 1× HBS-EP+ buffer. Fc-tagged receptors were captured onto a Protein A sensor chip by injecting receptor solutions at 0.1 mg/mL. Designed binders were diluted in the running buffer. For hTrkA-specific designs, four concentrations were tested in a threefold dilution series with a highest concentration of 2 µM. For cross-reactive designs, two concentrations, 0.2 and 2 µM, were tested against both targets. Analytes were injected sequentially at increasing concentrations using a single-cycle kinetics setup, without regeneration between injections. Dissociation was monitored by flowing running buffer over the chip surface following the final injection. Binding affinities were determined by global kinetic fitting of the sensorgrams to a 1:1 binding model using Cytiva Biacore evaluation software.

#### S13.5 Glycosylation

Proteins were produced in Expi293F (ThermoFisher A14527) cells at 1 mL scale in four repeats for 4 mL total volume, purified, and characterized by LC-MS/MS as previously described [48].

#### S13.6 Phosphopeptide binders

**Methods**

##### S13.6.1 DNA library preparation and transformation of the naïve library

Selected binder sequences were reverse translated and codon optimized for expression in *Saccharomyces cerevisiae*. Binder-encoding DNA was obtained as full-length synthetic oligonucleotide pools from Twist Bioscience. Pools were amplified with KAPA HiFi polymerase (Kapa Biosystems) in 25 µL reactions monitored on a Bio-Rad CFX96 real-time PCR instrument. Amplification was terminated at approximately half the maximum yield to limit overamplification. Products of the expected size were resolved by agarose gel electrophoresis, excised, and purified using a QIAquick cleanup kit (Qiagen). Approximately 6 µg of amplified insert DNA and 2–3 µg of pETCON vector were cotransformed into *S. cerevisiae* strain EBY100 to generate the naïve yeast display library. Full-length inserts were used directly, without an in vitro oligonucleotide-fragment assembly step.

##### S13.6.2 Library expansion, surface expression, and expression sorting

Transformed yeast were expanded overnight at 30 °C in selective synthetic complete medium lacking tryptophan and uracil (C−Trp−Ura) and supplemented with 2% (w/v) glucose. To induce surface display, cultures were diluted 1:10 into SGCAA autoinduction medium containing 0.2% (w/v) glucose and incubated overnight at 30 °C. Induced cells were washed with chilled phosphate-buffered saline containing 1% (w/v) bovine serum albumin (PBSF).

For selections that included an expression-enrichment step, cells were stained with FITC-conjugated anti-c-Myc antibody (Miltenyi Biotec) to detect the c-Myc epitope tag on the displayed binder. FITC-positive cells were collected by fluorescence-activated cell sorting (FACS), regrown in selective medium, and reinduced before target-binding selection.

##### S13.6.3 First target-binding selection (sort 1)

Induced yeast were washed in PBSF and stained with biotinylated target peptide at a final concentration of 500 nM for sort 1. For labeling with avidity, cells were incubated with biotinylated peptide, streptavidin–phycoerythrin (SAPE; Invitrogen, Thermo Fisher Scientific, cat. no. S866), and FITC-conjugated anti-c-Myc antibody. The SAPE molar concentration was one-quarter of the peptide concentration. For labeling without avidity, cells were first incubated with biotinylated peptide, washed, and then stained with SAPE and FITC-conjugated anti-c-Myc antibody.

Sort 1 was performed on a Sony SH800 cell sorter to collect cells displaying the binder and showing target-binding signal in the PE channel. Collected populations were regrown in selective medium and reinduced for subsequent selections.

##### S13.6.4 Competition sorting for phosphopeptide specificity

Competition sorting was performed with avidity labeling to enrich binders that preferentially recognized the phosphorylated target over its unphosphorylated counterpart. The two biotinylated peptides were labeled separately before incubation with yeast: the unphosphorylated decoy peptide was labeled with APC-conjugated anti-biotin antibody (BioLegend, cat. no. 405207), and the phosphorylated target peptide was labeled with SAPE. Induced, washed yeast were challenged with the labeled decoy and target at final peptide concentrations of 200 nM and 100 nM, respectively. Cells with high PE and low or undetectable APC signal were collected on a Sony SH800 cell sorter to enrich for phosphopeptide-preferring binders. APC-positive populations indicated decoy recognition, while populations positive in both channels indicated recognition of both peptides. Selected populations were regrown and reinduced between rounds.

##### S13.6.5 Single-colony isolation, colony picking, and clonal expression

Following the final selection, enriched yeast were plated on selective agar and incubated at 30 °C until isolated colonies were available. Individual colonies were transferred with a QPix XE colony picker into separate wells of culture plates containing C−Trp−Ura medium supplemented with 2% (w/v) glucose. Each colony was maintained as a separate clonal culture, with its source population and well position recorded. Cultures were grown overnight at 30 °C, diluted 1:10 into SGCAA autoinduction medium containing 0.2% (w/v) glucose, and incubated overnight at 30 °C to induce binder display.

##### S13.6.6 Clonal staining and Attune flow cytometry

Induced clonal cultures were washed in chilled PBSF and aliquoted for peptide-binding measurements. Cells were stained with biotinylated peptide, SAPE, and FITC-conjugated anti-c-Myc antibody using the avidity or sequential-labeling format described above. Where paired specificity measurements were performed, separate aliquots of each clone were stained with the phosphorylated and corresponding unphosphorylated peptides at matched concentrations. Selected clones were also evaluated across peptide dilution series.

Stained cells were analyzed on an Attune flow cytometer (Thermo Fisher Scientific), with FITC reporting surface expression and PE reporting peptide binding. The titrations in Fig. 5d,e report PE-positive events as a percentage of the full expression-positive, singlet-enriched parent population. For paired screens, phosphorylated- and unphosphorylated-peptide responses were compared for the same clone. Selected clones were advanced to sequence identification and further characterization.

Each peptide state was measured at 14 concentrations, including zero, with one acquisition per concentration. Matched zero-peptide controls defined sloped FITC–PE gates at the 99th percentile, held fixed across peptide states and concentrations. Four-parameter Hill curves were fitted to the phosphorylated-peptide responses and apparent response midpoints were reported, not equilibrium dissociation constants. Unmodified-peptide measurements were connected without fitting.

##### S13.6.7 Sequence identification of individual clones

Following colony picking, the binder-encoding insert from each picked colony was amplified by colony PCR using Phire Plant Direct PCR Master Mix and pETCON vector-specific primers. PCR products of the expected size were purified and submitted for Sanger sequencing. Sequences were compared with the ordered library to identify the recovered binder designs.

##### S13.6.8 NGS sequence counting and histone phosphosite selectivity

NGS was used to quantify recovery of H3pS10- and H3pT11-directed binders from a shared expression-selected library and from parallel first-round selections against each phosphopeptide. Paired-end reads were merged, quality filtered and translated. Sequences with ambiguous coding bases or stop codons were excluded. Binders were counted by exact amino-acid sequence, and their frequencies were normalized to the total accepted read pairs in each sample.

Enrichment was calculated as the sequence frequency after selection divided by its frequency in the shared input. Phosphosite selectivity was calculated as the ratio of sequence frequencies recovered from intended-target and off-target selections. For heatmaps, recovery for each binder was normalized to its intended target, set to 100%. Observations below 20 reads were retained. One population was sequenced per condition; these measurements describe relative recovery through sorting and regrowth.

#### S13.7 Phosphotriesterases

##### S13.7.1 Small-scale protein expression and purification for screening

Cloning, expression, and purification largely followed our previously described procedures for designing and characterizing *de novo* metallohydrolases [6, 55], with modifications as described below. Genes encoding 288 designs were codon-optimized for *E. coli* expression and ordered from IDT as three 96-well eBlock plates with Golden Gate overhangs. DNA fragments were resuspended in nuclease-free water (Invitrogen) to 10 ng/$\mu$L and cloned into pGG-T7-cStrepII (Addgene plasmid #262810), yielding proteins in the format MSG-design-GSA-WSHPQFEK with a C-terminal Strep-tag II. Golden Gate assembly was performed in 5 $\mu$L reactions using the NEBridge Golden Gate Assembly Kit (BsaI-HF v2; New England Biolabs), with insert DNA at $\geq$ 8.8 nM and vector at 4.4 nM. Reactions were thermocycled for 30 cycles of 37 °C for 2 min and 16 °C for 2 min, followed by 60 °C for 10 min and a hold at 12 °C, with the heated lid set to 105 °C. Assembly products (1.5 $\mu$L) were added to 6 $\mu$L chemically competent BL21(DE3) *E. coli* (Intact Genomics), mixed by pipetting, and transformed by heat shock at 42 °C for 45 s in a 96-well format. Following recovery in SOC medium for 2.5 h at 37 °C, cells were grown overnight in LB containing 50 $\mu$g/mL kanamycin at 37 °C with shaking. Glycerol stocks were prepared by mixing 100 $\mu$L overnight culture with 100 $\mu$L 50% glycerol and stored at −80 °C. These initial cultures were not sequence-verified and could be polyclonal.

Proteins were expressed in two 1.1 mL cultures per design using TB-II medium (MP Biomedicals; 50 g/L), supplemented with 50 $\mu$g/mL kanamycin, in 2 mL deep-well plates (Corning). Cultures inoculated from overnight LB cultures were incubated at 37 °C and 1200 rpm for 1.5 h, induced with 1 mM IPTG (Fisher Scientific) and 1 mM $ZnSO_4$ (MilliporeSigma), and immediately transferred to room temperature for 21 h at 1200 rpm. Cells were harvested by centrifugation (4,198 × $g$, 8 min, 25 °C). For batches purified immediately after harvest, pellets were processed directly for lysis without freezing.

The base purification buffer contained 40 mM HEPES, 50 mM NaCl, and 200 $\mu$M $ZnSO_4$, pH 8.0 (buffer components from MilliporeSigma). Buffers were prepared in Milli-Q water and filtered through 0.2 $\mu$m PES filters. Each pellet from a 1.1 mL culture was resuspended in 300 $\mu$L base buffer supplemented with 1 mg/mL lysozyme, 10 $\mu$g/mL DNase I, 1 mM $MgCl_2$, 0.5 mg/mL polymyxin B sulfate (lysis reagents from Sigma-Aldrich), and one Pierce protease inhibitor tablet (Thermo Fisher Scientific) per 50 mL, giving 600 $\mu$L lysis buffer per design. Plates were incubated at 37 °C and 1200 rpm for 1 h, and lysates were clarified separately by centrifugation (4,200 × $g$, 40 min, 16 °C).

Purification was performed in 96-well fritted plates (Agilent) containing a 100 $\mu$L bed of Strep-Tactin XT 4Flow resin (IBA Lifesciences) per well, equilibrated with 3 × 400 $\mu$L base buffer. Corresponding clarified lysates from the two culture wells for each design were combined only upon loading onto the same resin well. The fritted plates were sealed at the top and bottom, gently inverted, and incubated at 4 °C with orbital agitation at 20 rpm for 20–40 min to allow binding. Lysates were then drawn through the resin using a vacuum manifold, and the resin was washed with 3 × 400 $\mu$L base buffer. The underside of each fritted plate was rinsed with water and thoroughly dried before elution. Protein was eluted into deep-well plates with 300 $\mu$L base buffer containing 50 mM biotin (MilliporeSigma), readjusted to pH 8.0. Elution proceeded by gravity for > 10 min, followed by centrifugation at 10 × $g$ for 90 s and 250 × $g$ for 1 min. Eluates were prepared for screening as described below.

##### S13.7.2 Large-scale protein expression and purification for characterization

Single colonies corresponding to designs selected following screening were isolated by dilution and plating of their glycerol stocks on kanamycin-containing LB agar. For each design, 1–3 colonies were cultured individually and screened by colony PCR using T7 and T7 terminator primers followed by Sanger sequencing (Genewiz/Azenta). Glycerol stocks were prepared from selected single colonies, and one clone per design was used for subsequent experiments.

Stocks from the selected colonies were used to inoculate 5 mL LB cultures containing 50 $\mu$g/mL kanamycin, which were grown at 37 °C and 225 rpm for 16–20 h. Starter cultures were transferred into 50 mL TB-II containing 50 $\mu$g/mL kanamycin in 250 mL baffled flasks and grown at 37 °C and 225 rpm to $OD_{600} \sim 0.6$. Expression was induced with 1 mM IPTG and 1 mM $ZnSO_4$, followed immediately by incubation at 18 °C and 225 rpm for 16–20 h. Cells were harvested by centrifugation (4,198 × $g$, 8 min, 25 °C), and pellets were stored at −80 °C.

Pellets were resuspended in 20 mL lysis buffer with the same composition as above, but without polymyxin B, and sonicated on ice using a Qsonica Q500 with a four-tip horn (60% amplitude; 10 s on/20 s off; 180 s total on-time). Lysates were clarified by centrifugation (14,000 × $g$, > 45 min, 4 °C). Supernatants were loaded onto gravity-flow columns (Bio-Rad) containing 1 mL Strep-Tactin XT resin equilibrated with 3 × 3 column volumes (CV) of base buffer. Affinity purification was performed at 25 °C. Columns were washed with 3 × 5 CV base buffer; an initial 0.6 CV biotin elution fraction was discarded, followed by collection of two 1.1 CV fractions. Eluates were purified by size-exclusion chromatography on a Superdex 75 Increase 10/300 GL column using an ÄKTA pure system (Cytiva) in base purification buffer. Samples were loaded in approximately 1.0–1.1 mL volumes, with concentration using 10 kDa (or 3 kDa) MWCO Amicon centrifugal filters (MilliporeSigma) when necessary. Selected SEC fractions were collected and used immediately for characterization.

##### S13.7.3 Colorimetric activity screening of designed phosphotriesterases

Purified proteins were preincubated with bicarbonate to promote lysine carbamate formation. A 550 mM $NaHCO_3$ stock (MilliporeSigma) prepared in reaction buffer (40 mM HEPES, 50 mM NaCl, 200 $\mu$M $ZnSO_4$, pH 8.0) was added at 30 $\mu$L per 300 $\mu$L eluate, giving 50 mM $NaHCO_3$. The three elution plates were tightly sealed with aluminum film and incubated for $> 1.5$ h at room temperature with shaking at 600 rpm.

Activity was measured in Corning 384-well black/clear flat-bottom nonbinding assay plates. Aliquots of bicarbonate-treated enzyme (30 $\mu$L) were transferred to assay wells and centrifuged at 1,750 $\times\ g$ for 2 min before adding 30 $\mu$L substrate solution containing 1.2 mM paraoxon-ethyl (PESTANAL analytical standard; Sigma-Aldrich, 36186) and 2% (v/v) methanol (MeOH) in reaction buffer. The substrate solution was prepared by adding methanol, paraoxon, and buffer in that order and mixing thoroughly but gently by pipetting. Final 60 $\mu$L reactions contained 40 mM HEPES, 50 mM NaCl, 200 $\mu$M $ZnSO_4$, 25 mM $NaHCO_3$, approximately 23 mM biotin, 600 $\mu$M paraoxon, and 1% (v/v) MeOH, pH 8.0. Enzyme concentrations were not normalized across designs and were diluted twofold upon substrate addition. Plates were covered with clear plastic plate lids, shaken with double-orbital motion at 282 cpm for 15 s, and monitored for nitrophenol formation at 405 nm for 1 h at 25 °C using a BioTek Synergy Neo2 plate reader.

Screening activities were subsequently normalized to protein concentrations estimated from $A_{280}$ measurements and sequence-derived extinction coefficients for the expressed, tagged constructs. Concentration-normalized activities, together with scaffold assignments, were used to prioritize hits and scaffold families for large-scale expression and kinetic characterization.

##### S13.7.4 Kinetic analysis of designed phosphotriesterases

Steady-state kinetics were measured for eight affinity- and SEC-purified *de novo* phosphotriesterase designs generated with RFD4-Proteina. Enzyme solutions in SEC buffer were supplemented with 50 mM $NaHCO_3$ using a 550 mM stock in the same buffer and incubated for at least 2 h at room temperature in sealed tubes shaking at 600 rpm.

The selected assays used 500 nM enzyme for BLITZ-5 (p1D10), p1F2, BLITZ-2 (p2G5), BLITZ-3 (p2G6), p2H7, and BLITZ-1 (p2H11), and 1 $\mu$M for p2H10 and BLITZ-4 (p3D1). Measurements were performed in technical triplicate at each substrate concentration. Protein concentrations were determined by absorbance at 280 nm using a NanoDrop spectrophotometer and sequence-derived extinction coefficients for the expressed, tagged constructs [106]. Enzyme-concentration uncertainty was propagated as described below.

Reactions were performed in Corning 96-well half-area black/clear flat-bottom nonbinding plates at 25 °C. Each 80 $\mu$L reaction contained 40 $\mu$L enzyme and 40 $\mu$L substrate solution, giving final conditions of 40 mM HEPES, 50 mM NaCl, 200 $\mu$M $ZnSO_4$, 25 mM $NaHCO_3$, pH 8.0, and 1% (v/v) MeOH. A 4.8 mM paraoxon stock was prepared in reaction buffer containing 2% (v/v) MeOH by adding methanol, paraoxon, and buffer in that order and mixing thoroughly but gently by pipetting. The stock was used to prepare six successive twofold dilutions down to 75 $\mu$M paraoxon in the same buffer containing 2% MeOH. Upon 1:1 mixing with enzyme solution, the seven final paraoxon concentrations ranged from 37.5 $\mu$M to 2.4 mM. Plates were covered with clear plastic plate lids, shaken with double-orbital motion at 282 cpm for 5 s, and monitored at 405 nm for 15–25 min using the BioTek Synergy Neo2.

Absorbance was calibrated using an eight-point 4-nitrophenol standard curve (spectrophotometric grade; Sigma-Aldrich, 1048), prepared by serial dilution from 200 to approximately 1.6 $\mu$M in the final assay buffer and measured in technical triplicate at 80 $\mu$L volume for 10 min at 25 °C. For plotting, no-enzyme background traces were subtracted at each substrate concentration, and progress curves were shifted to zero at the first recorded measurement. Initial velocities were determined by fitting the raw, unsmoothed enzyme and substrate-matched blank traces separately by linear regression with a free intercept, subtracting the blank slope from the enzyme slope, converting to product formation rates using the standard curve, and normalizing by the final enzyme concentration.

Initial-rate fitting windows were selected automatically at each substrate concentration using fixed analysis criteria, with a common window for the three enzyme wells and the matched blank. Each window began at the first recorded measurement ($t = 0$) and was extended one reading at a time from a minimum of five time points (four only for the 37.5 $\mu$M condition of the selected BLITZ-1 (p2H11) assay), up to 180 s. Extension stopped before the first candidate window in which product accumulation in any enzyme well exceeded 10% of the initial substrate concentration, raw absorbance in any included well exceeded 2.5 AU, or curvature was detected. Product accumulation was calculated from the blank-corrected change since the first read. The last acceptable window was retained, ending at the last available reading at or before the time limit if no criterion failed.

Curvature was assessed once a candidate window contained at least eight readings by comparing linear slopes fitted to the first and second halves of the mean blank-corrected trace. Curvature was flagged only when the absolute slope difference exceeded both 20% of the larger absolute slope and three times the root-sum-of-squares of the two slope uncertainties. Substrate conditions were excluded if the minimum window already exceeded the conversion or absorbance cutoff. The selected BLITZ-1 (p2H11) assay retained all seven substrate concentrations; its 37.5 $\mu$M condition used four readings at 0, 16, 32, and 48 s, with estimated conversion of 7.6%.

The remaining mean velocities, each calculated from three technical replicates, were fitted by inverse-variance-weighted nonlinear regression, using rate and window-sensitivity uncertainties as weights, to the Michaelis–Menten equation:

$$\frac{v_0}{[E]_0} = \frac{k_{\text{cat}}[S]}{K_{\text{M}} + [S]}, \tag{S13}$$

where $v_0$ denotes the initial velocity, $[E]_0$ the total expressed protein-chain concentration, and $[S]$ the substrate concentration. The reported $k_{\text{cat}}$ values are therefore apparent turnover numbers; active enzyme concentrations were not independently determined. One selected assay per design is reported. Rate enhancement is reported as $k_{\text{cat}}/k_{\text{uncat}}$, where $k_{\text{uncat}}$ is the first-order rate constant for uncatalyzed paraoxon hydrolysis.

Uncertainties in $k_{\text{cat}}$, $K_{\text{M}}$, and $k_{\text{cat}}/K_{\text{M}}$ were propagated through 2,000 seeded Monte Carlo realizations, with a joint Michaelis–Menten refit for each realization. Enzyme-concentration uncertainty used the measured sample SD/mean from triplicate $A_{280}$ readings for the corresponding stock preparation, without division by $\sqrt{3}$, together with assumed relative SDs of 10% for the predicted extinction coefficient and 2% for working-enzyme preparation. The 10% extinction-coefficient term was an explicit analysis assumption. Substrate preparation included assumed relative SDs of 2% for the substrate stock and 2% for the top dilution, a 2% volume CV for each serial transfer and diluent dispense, and uniform pipette biases bounded by ±1%. The serial-dilution model assumed equal 100 $\mu$L transfer and diluent volumes; upstream dilution errors were inherited by subsequent dilutions. Assay-volume biases jointly affected enzyme and substrate concentrations and the assumed volume-dependent absorbance path length.

Rate uncertainty used the larger of technical-replicate SD and slope-regression uncertainty, including an autocorrelation adjustment and a read-quantization floor of

$1 \times 10^{-3}/\sqrt{12}$ AU. Blank-slope uncertainty, with an assumed additional floor of 10% of the blank rate, was shared among the enzyme wells at each substrate concentration. Window sensitivity was estimated from the SD of slopes obtained using 70%, 85%, and 100% of the selected initial window and propagated as a shared component across substrate concentrations. Random assay dispensing was not added again because it was already represented in observed replicate dispersion. Calibration-slope uncertainty incorporated standard-well dispersion and uncertain standard concentrations. Shared enzyme-stock, extinction-coefficient, calibration, and pipette-bias terms were retained across substrate concentrations, and $k_{cat}/K_M$ was calculated within each refit to preserve $k_{cat}$–$K_M$ covariance.

Parameter values were reported as Monte Carlo mean ± propagated SD, with medians and 95% propagation intervals retained in the analysis. Rate-plot error bars show propagated standard uncertainties in $v_0/[E]_0$ and $[S]$, rather than SEM; progress-curve shading shows technical-replicate SD. These propagated uncertainties do not quantify biological variability or unmeasured between-day or between-reader calibration shifts. Uncertainties were rounded upward to one significant figure, and central values were rounded to the nearest value at the corresponding decimal place, directly from full-precision results.

###### S13.7.5 PTE active-site alanine variants

We compared BLITZ-1 (p2H11) with alanine variants of seven first-shell catalytic residues targeting the six designed metal-coordinating residues (including the general base D17) and the tryptophan transition-state stabilizer W76. Each construct was screened in one eluate well with 600 $\mu$M paraoxon in 80 $\mu$L reactions containing 1% methanol and legacy buffer at approximately 25 °C. Expression cultures were normalized to 2 mL. The mean trace from 12 plate-reference wells was subtracted at each time point. Observed rates were calculated by ordinary least-squares fits with a free intercept over the same first 60 s after the first recorded read (six readings at 12-s intervals). Figure 6g shows $100\, s_{variant}/s_{parent}$, where $s$ is the fitted absorbance slope.

For Supplementary Fig. 32, each reference-corrected trace was additionally shifted by its first value to give $\Delta A_{405}$; this constant shift leaves the fitted slopes unchanged. Traces stop at their original overflow boundaries. Triangles mark the first overflow using the raw instrument limit of 4.0 absorbance units transformed by the same reference and first-reading subtractions.

##### S13.8 Lenacapavir-precursor ligases

The lenacapavir-precursor campaign is also described in the companion amide-ligase study [62]; D52 and D58 here correspond to LL52 and LL58 there.

###### S13.8.1 Construction of ligase variants

Genes encoding D52_C177A and D58_C144A were constructed by overlap-extension PCR with end primers designed for Golden Gate assembly (Table S30) and cloned into a pET26-ccdB vector using BsaI-HFv2, fusing each gene to a C-terminal hexahistidine tag to give pET26_D52_C177A and pET26_D58_C144A.

###### S13.8.2 Protein production and purification

Designs were expressed in chemically competent *E. coli* BL21(DE3). 5 mL LB medium containing kanamycin (50 $\mu$g/mL) was inoculated and cultured for 18 h; 500 $\mu$L of this starter culture was used to inoculate baffled flasks containing auto-inducible TBII medium (50 mL) and kanamycin (50 $\mu$g/mL). Cultures were grown at 37 °C, 180 r.p.m. for 6 h, then shifted to 25 °C for a further 16 h. Cells were collected by centrifugation (3,220 $\times g$, 10 min), resuspended in lysis buffer (50 mM HEPES, 300 mM NaCl, pH 7.4, 10 mM imidazole, 10 $\mu$g/mL DNase I, 1 mM phenylmethylsulfonyl fluoride), and lysed by sonication. Lysates were clarified by centrifugation (27,216 $\times g$, 15 min), and the soluble fraction was purified by Ni-NTA affinity chromatography (Qiagen), washing

**Table S30** Primers used to construct D52_C177A and D58_C144A by overlap-extension PCR and Golden Gate assembly.

| Primer | Sequence (5′–3′) |
|---|---|
| D52_C177A_F | GCATTAAAGGCATTGGCAACGCGATTGTGACCATT |
| D52_C177A_R | GTTGCCAATGCCTTTAATGC |
| D52_GGFlank_F | atactacggtctcaaggaGAAGAAGAAATCGAGAAAGCG |
| D52_GGFlank_R | gcattacggtctcgggaaccCAGTTCTTCCAGAATTTTCAGC |
| D58_C144A_F | GTCATCCGGCGAACGCGGAAGTGACTACC |
| D58_C144A_R | GTTCGCCGGATGAC |
| D58_GGFlank_F | atactacggtctcaaggaCCGCCGGCTGAA |
| D58_GGFlank_R | gcattacggtctcgggaaccCGCCGGTTCAAACAC |

with lysis buffer supplemented with an additional 10 mM imidazole before eluting with 50 mM HEPES, 300 mM NaCl, pH 7.4, 300 mM imidazole. Purified protein was desalted into 100 mM potassium phosphate buffer, pH 6, using 10DG columns (Bio-Rad).

##### S13.8.3 Analytical-scale biotransformations

Analytical-scale biotransformations were performed with activated ester **1** (1 mM), enantiopure amine **2** (0.5 mM), and the relevant purified enzyme (50 $\mu$M) in 100 mM potassium phosphate buffer, pH 6, with 20% DMSO, incubated at 30 °C, 850 r.p.m. for 16 h. No-enzyme controls were run separately for each enantiomer under the same conditions. Reactions were quenched with 1 volume of acetonitrile containing 0.1% HCl and shaken at 30 °C, 850 r.p.m. for 20 min; precipitated protein was removed by centrifugation (3,220 $\times g$, 15 min), and the supernatant was used for UPLC analysis. Conversion was calculated as $100(A_{\text{amide}}/315.89)/(A_{\text{amide}}/315.89 + A_{\text{amine}}/188.76)$, where $A$ denotes integrated chromatographic peak area. We calculated means and population standard deviations (denominator $n$) from three measurements per condition, without background subtraction.

##### S13.8.4 Campaign comparison

For the campaign comparison in Figure 6m, RFD3 designs were initially screened in lysate, whereas RFD4-Proteina designs were screened after purification. Each design's mean conversion was centered on its campaign's background mean and divided by the background sample SD. The available comparison contains 71 RFD3 designs and the 55 RFD4-Proteina designs that expressed well among 67 total [62]. Figure 6n shows D52 and no-enzyme controls from the enantiopure-amine assay described above, without background subtraction.

##### S13.8.5 Chromatographic analysis

Reaction supernatants were analyzed on a 1290 Infinity II UPLC system (Agilent) with the diode-array detector set to 254 nm, using a Kinetex XB-C18 column (100 Å, 5 $\mu$m, 50 × 2.1 mm; Phenomenex). Mobile phase A was water with 0.1% trifluoroacetic acid and mobile phase B was acetonitrile; the flow rate was 1 mL min$^{-1}$ and the injection volume was 2 $\mu$L. After injection the mobile phase was held at 20% B, increased to 95% B over 60 s, held at 95% B for 30 s, and returned to 20% B over 6 s. Peaks were assigned by comparison with chemical standards. Representative traces for a no-enzyme control and for D52 with the ($S$)-amine are shown in Supplementary Fig. 23.

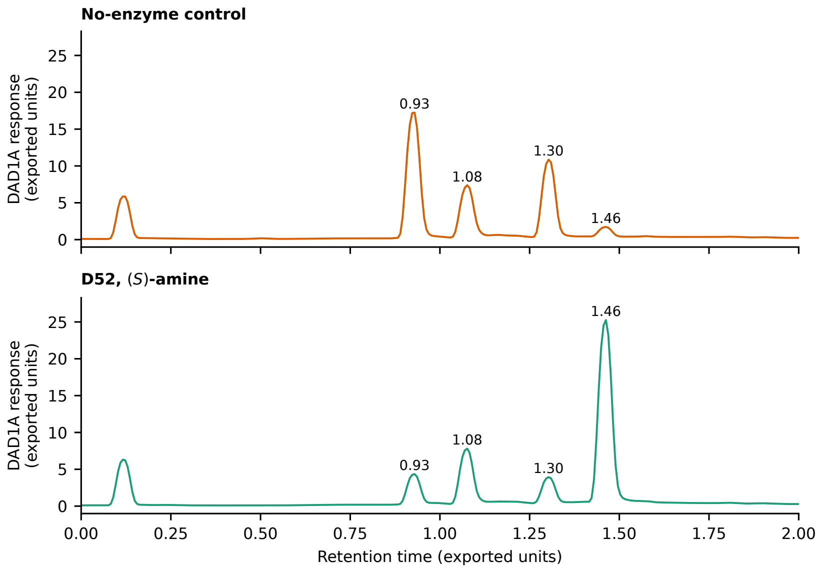

**Supplementary Fig. 23 Representative UPLC traces for the lenacapavir-precursor amide ligation.** DAD1A detector responses at 254 nm for a no-enzyme control (top) and D52 with the ($S$)-amine (bottom) after 16 h, run on the column and gradient described above. Labels mark peak retention times.

### S14 Supplementary data

#### S14.1 Calcium-dependent calmodulin binders

**Table S31 Calmodulin binding.** Reported overall apparent sub-micromolar $K_D$ estimates (95% confidence intervals) measured by BLI with 2 mM calcium or 3 mM EDTA. Estimates combine independent replicates. Intervals for off-rate-limited estimates are conditional on the assay resolution limit (Supplementary Section S13.1). A dash indicates that a sub-micromolar affinity was not established. Calcium-dependent hits: 12/18 (66.7%).

| Design | Calcium | EDTA |
|---|---|---|
| `c13_31860910_4_bsfinal_2` | 1.52 (1–2.2) nM | 55.8 (48–64) nM |
| `c09_31717194_6_bsfinal_3` | — | — |
| `c09_31822041_1_bsfinal_3` | — | — |
| `c09_31822041_1_bsfinal_4` | — | — |
| `c06_31713662_9_bsfinal_5` | — | — |
| `c09_31868515_1_bsfinal_6` | 3.31 (3.1–3.6) nM | — |
| `c06_31713662_5_bsfinal_6` | 10.2 (9–11) nM | — |
| `c09_31717194_6_bsfinal_7` | 798 (270–2400) pM | — |
| `c09_31822041_1_bsfinal_7` | 37 (35–39) nM | — |
| `c13_31868515_4_bsfinal_10` | 69 (3.4–92) pM | — |
| `c09_31822041_1_bsfinal_11` | 12.3 (11–13) nM | — |
| `c09_31860910_1_bsfinal_11` | 2.48 (2.4–2.6) nM | — |
| `c06_31713662_10_bsfinal_13` | 3.31 (2.5–4.4) nM | — |
| `c13_31860910_4_bsfinal_15` | 12.6 (12–13) nM | — |
| `c09_31868515_1_bsfinal_15` | 21.6 (16–29) nM | — |
| `t31682405_1_bsfinal_16` | 90.8 (3.7–94) pM | — |
| `c06_31713662_12_bsfinal_17` | 748 (440–1300) pM | — |
| `c09_31822041_1_bsfinal_18` | 6.51 (6.3–6.7) nM | 149 (130–170) nM |

#### S14.2 Peptide–MHC binders

Clonal measurements are listed in Supplementary Table S32. Library screening is shown in Supplementary Fig. 24; individual-binder titrations and specificity measurements are shown in Fig. 4b.

**Table S32 Clonal characterization of pMHC binders.** Data for pMHC binders tested clonally on yeast display. Binders were identified by sequencing single colonies after library screens. Normalized enrichment is the percentage of sequenced colonies for each target containing that sequence. Specificity was determined by staining with related and unrelated off-targets. Estimated $K_D$ values represent empirical half-maximal concentrations with avidity. A dash indicates an unmeasured affinity.

| Binder | Colony count | Normalized enrichment (%) | Specific | Estimated $K_D$ with avidity |
|---|---|---|---|---|
| **SARS-CoV-1 (30 colonies sequenced)** | | | | |
| SARS_c7 | 1 | 3.3 | Yes | 281.3 pM |
| SARS_c9 | 2 | 6.7 | No | — |
| SARS_c10 | 2 | 6.7 | Yes | 241.4 pM |
| SARS_c11 | 1 | 3.3 | Yes | 226.6 pM |
| SARS_c12 | 3 | 10.0 | Yes | 700 pM |
| SARS_c1 | 1 | 3.3 | No | — |
| SARS_c13 | 1 | 3.3 | Yes | 248.2 pM |
| SARS_c15 | 2 | 6.7 | Yes | 1.2 nM |
| SARS_c20 | 1 | 3.3 | No | — |
| SARS_c21 | 1 | 3.3 | Yes | 411.6 pM |
| SARS_c3 | 1 | 3.3 | Yes | 259 pM |
| SARS_c27 | 1 | 3.3 | No | — |
| SARS_c30 | 2 | 6.7 | Yes | 680 pM |
| SARS_c5 | 3 | 10.0 | No | — |
| SARS_c32 | 2 | 6.7 | Yes | 1.2 nM |
| SARS_c33 | 1 | 3.3 | No | — |
| SARS_c34 | 1 | 3.3 | Yes | 250.2 pM |
| SARS_c37 | 3 | 10.0 | Yes | 147.3 pM |
| SARS_c41 | 1 | 3.3 | Yes | 245.4 pM |
| **Phox2B (41 colonies sequenced)** | | | | |
| PHOX_c4 | 3 | 7.3 | No | — |
| PHOX_c5 | 1 | 2.4 | No | 890 pM |
| PHOX_c9 | 10 | 24.4 | No | 1.25 nM |
| PHOX_c10 | 10 | 24.4 | No | 4.99 nM |
| PHOX_c12 | 13 | 31.7 | Yes | 4.19 nM |
| PHOX_c13 | 2 | 4.9 | Yes | 1.6 nM |
| PHOX_c29 | 2 | 4.9 | No | 4.26 nM |
| **MART-1 (20 colonies sequenced)** | | | | |
| MART_c11 | 15 | 75.0 | Yes | 400 pM |
| MART_c23 | 3 | 15.0 | Yes | 320 pM |
| MART_c27 | 2 | 10.0 | Yes | 400 pM |

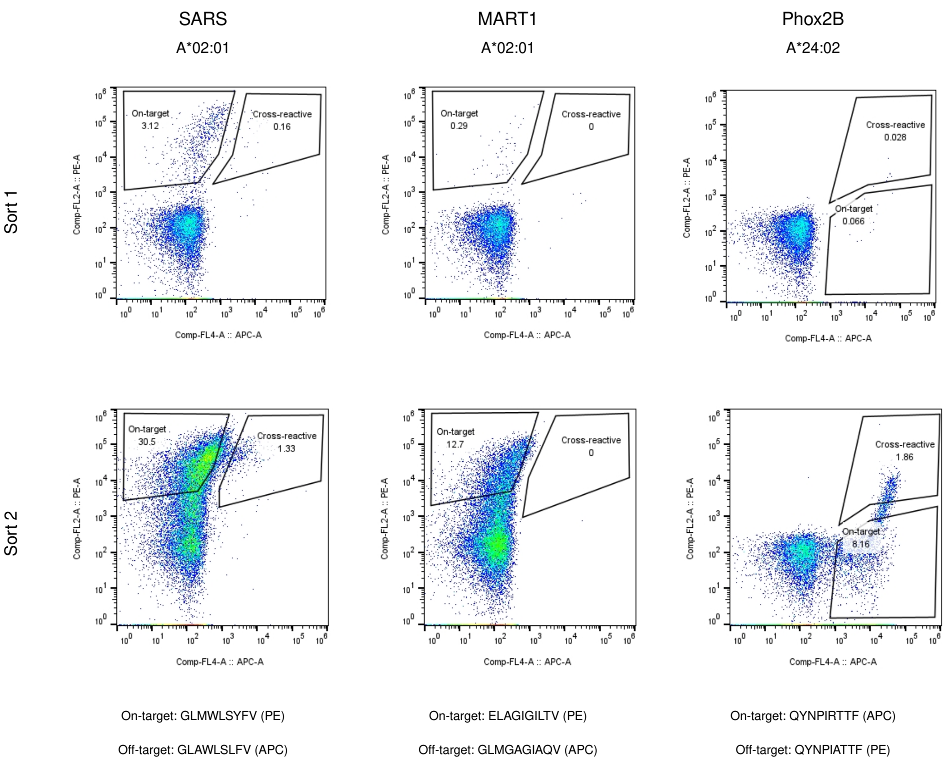

**Supplementary Fig. 24 Yeast surface display library screening for pMHC binders.** Libraries were first sorted for surface expression using an anti-C-Myc FITC antibody (not shown), followed by two binding sorts with target and related off-target pMHC tetramers. Columns show SARS, MART1 and Phox2B; rows show the first and second binding sorts. The vertical axis shows PE fluorescence and the horizontal axis shows APC fluorescence. The on-target signal is PE for SARS and MART1 and APC for Phox2B. Gates distinguish on-target and cross-reactive populations. Peptide sequences and fluorophores are indicated below each column.

#### S14.3 Intrinsically disordered region binders

**Table S33 IDR screening hit rates.** Screening hits among 94 designs tested per target.

| Target | Screening hits | Hit rate |
|---|---|---|
| G3BP1 | 11/94 | 11.7% |
| PAR2 | 16/94 | 17.0% |
| YAP1 | 18/94 | 19.1% |
| EA1 | 0/94 | 0.0% |
| TDP-A1 | 0/94 | 0.0% |
| N17 | 6/94 | 6.4% |
| Tau-153 | 3/94 | 3.2% |
| PER2 | 5/94 | 5.3% |

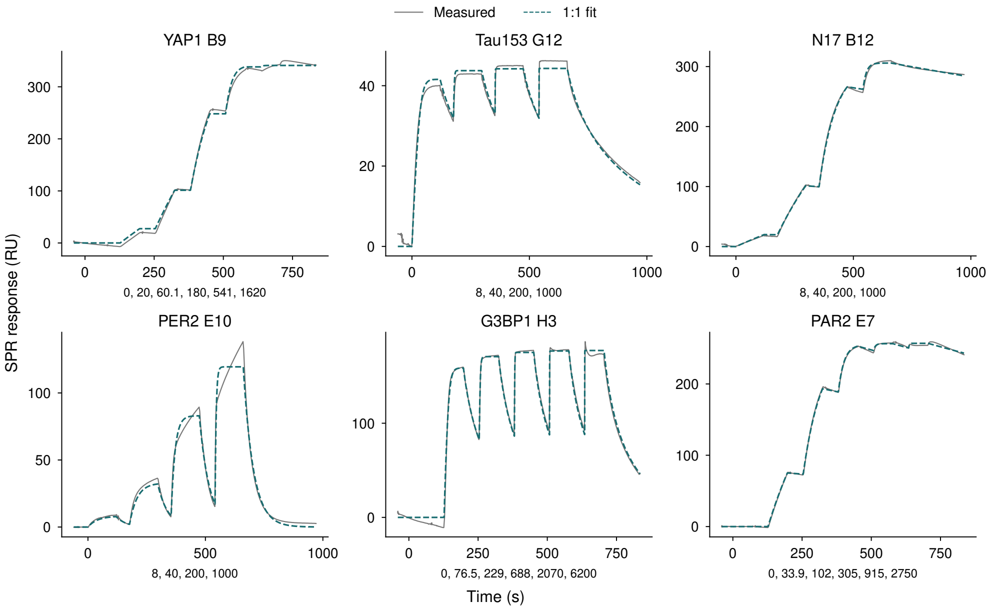

**Supplementary Fig. 25 IDR binder titrations.** Single-cycle SPR titrations for one selected binder per target. Gray solid lines show measured responses; teal dashed lines show 1:1 kinetic fits. Analyte concentrations (nM) are listed beneath each panel. Axes vary between panels. The G3BP1 trace contains an injection artifact.

#### S14.4 Trk receptor binders

**Table S34 Trk binding hit rates.** TrkA and TrkB hits require an estimated $K_D < 1$ $\mu$M; cross-reactive hits must meet this cutoff for both targets. TrkA hits also require an SPR response $\geq 20$ RU after reference and buffer subtraction. TrkC hits are based on screening; precise affinities have not yet been determined. Denominators include transformation failures and exclude positive and buffer controls.

| Design task | Hits/designs | Rate |
|---|---|---|
| Human-specific TrkA | 68/94 | 72.3% |
| Human/mouse cross-reactive TrkA | 18/94 | 19.1% |
| TrkB | 1/94 | 1.1% |
| TrkC | 20/94 | 21.3% |

#### S14.5 Glycosylation

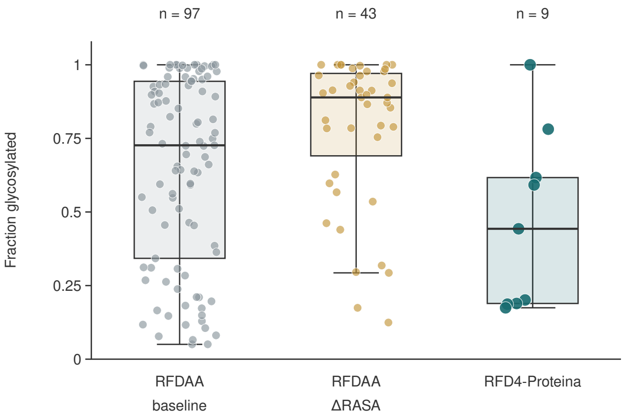

**Supplementary Fig. 26 Fraction glycosylated in the Figure 5 cohorts.** Boxplots and individual points show the fraction of sites glycosylated, measured by LC–MS, for 97 baseline RFDAA, 43 ΔRASA-filtered RFDAA and nine RFD4-Proteina profiles, including C12. Fraction glycosylated is one minus the unoccupied fraction. The same profiles are used for the underprocessing-score boxplots in Figure 5c: profiles below 5% occupancy and the F8 trimer are excluded. Boxes span the 25th–75th percentiles, with the median marked by a horizontal line. Whiskers extend to the most extreme observed values within 1.5 interquartile ranges of the box. All profiles, including outliers, are shown as points; horizontal jitter is used only for visibility. Counts describe the characterized samples; campaign hit rates were not assessed.

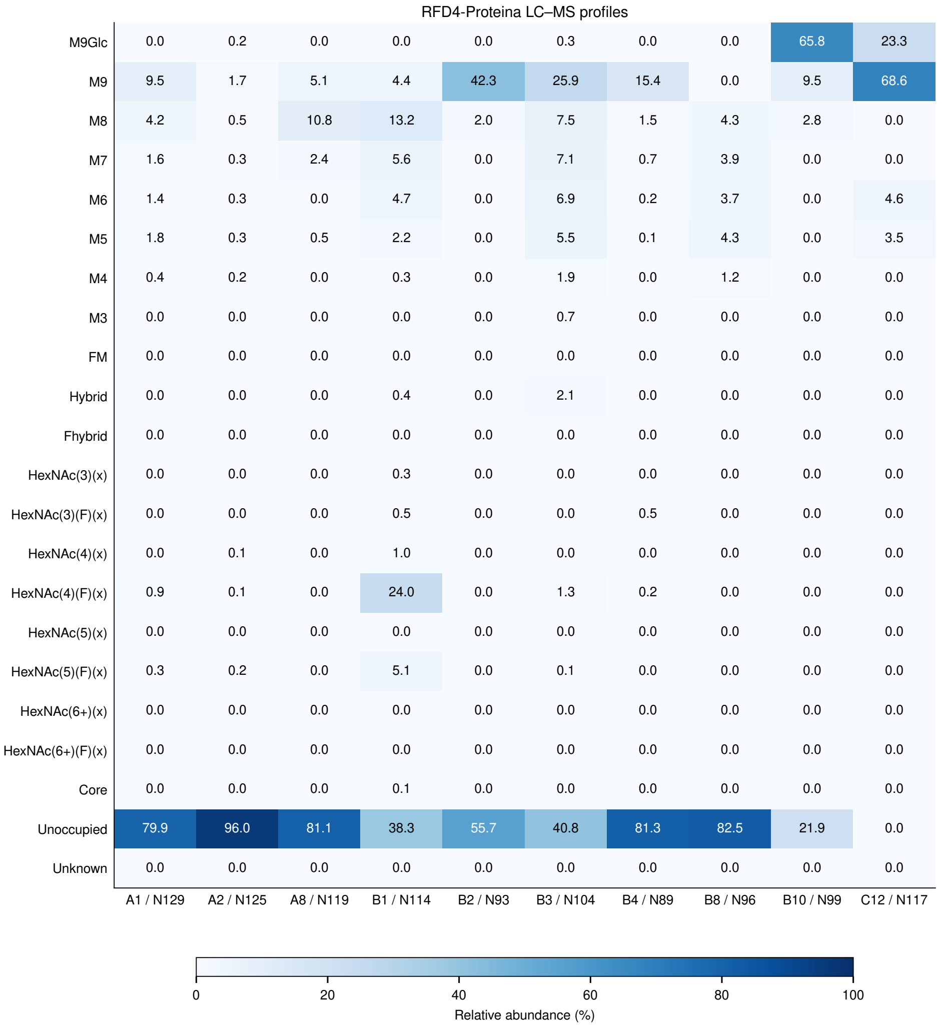

**Supplementary Fig. 27 RFD4-Proteina glycosylation profiles.** LC–MS glycoform abundances for ten designs, including the low-occupancy A2 sample excluded from the main-text cohort comparison. Columns identify samples; rows show glycoforms and unoccupied sites. Values are percentages of the total profile, including unoccupied sites.

#### S14.6 Phosphopeptide binders

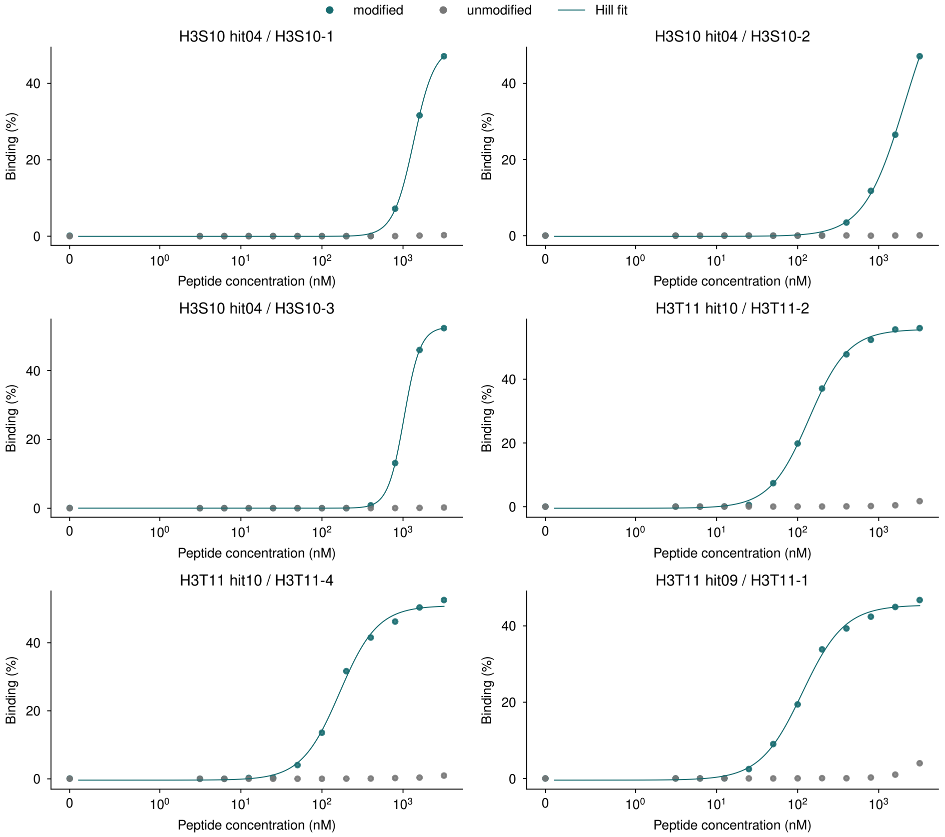

**Supplementary Fig. 28 Phosphopeptide-binder dilution series.** Yeast-display binding to phosphorylated and unmodified peptides for H3pS10 hit04, H3pT11 hit10 and H3pT11 hit09. Points show measurements and lines show Hill fits describing apparent binding in yeast display, rather than monovalent solution affinity. Concentration axes are linear near zero and logarithmic above 1 nM.

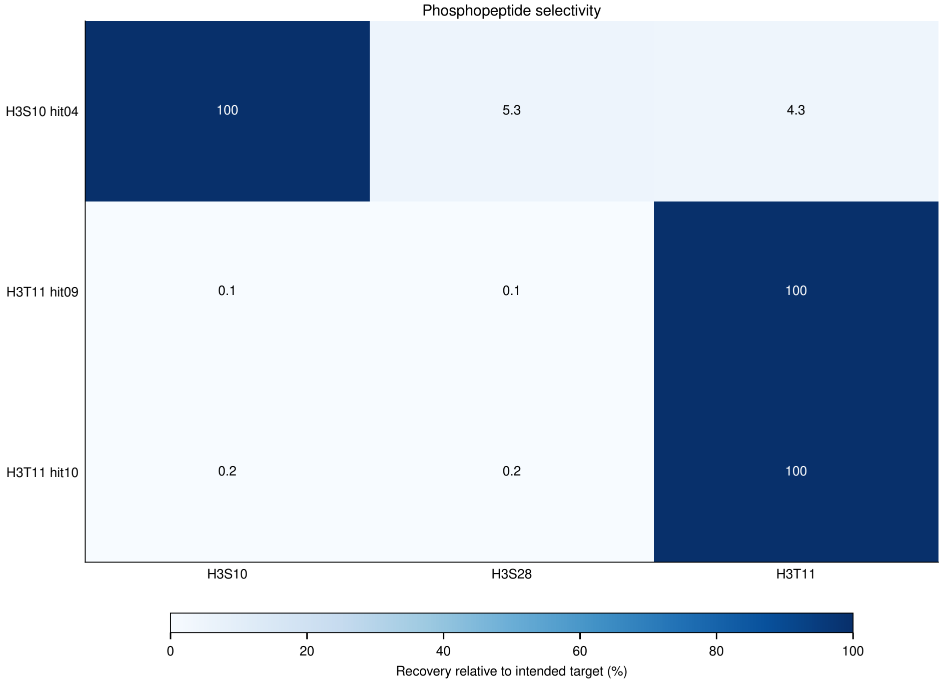

**Supplementary Fig. 29 Phosphopeptide selectivity.** Sequencing frequencies after sorting against each phosphopeptide are normalized to the corresponding binder's frequency after sorting against its intended target. Values show percent of intended-target recovery; cell labels give values above the color-scale maximum. This enrichment-based readout is not a binding affinity.

#### S14.7 Phosphotriesterases

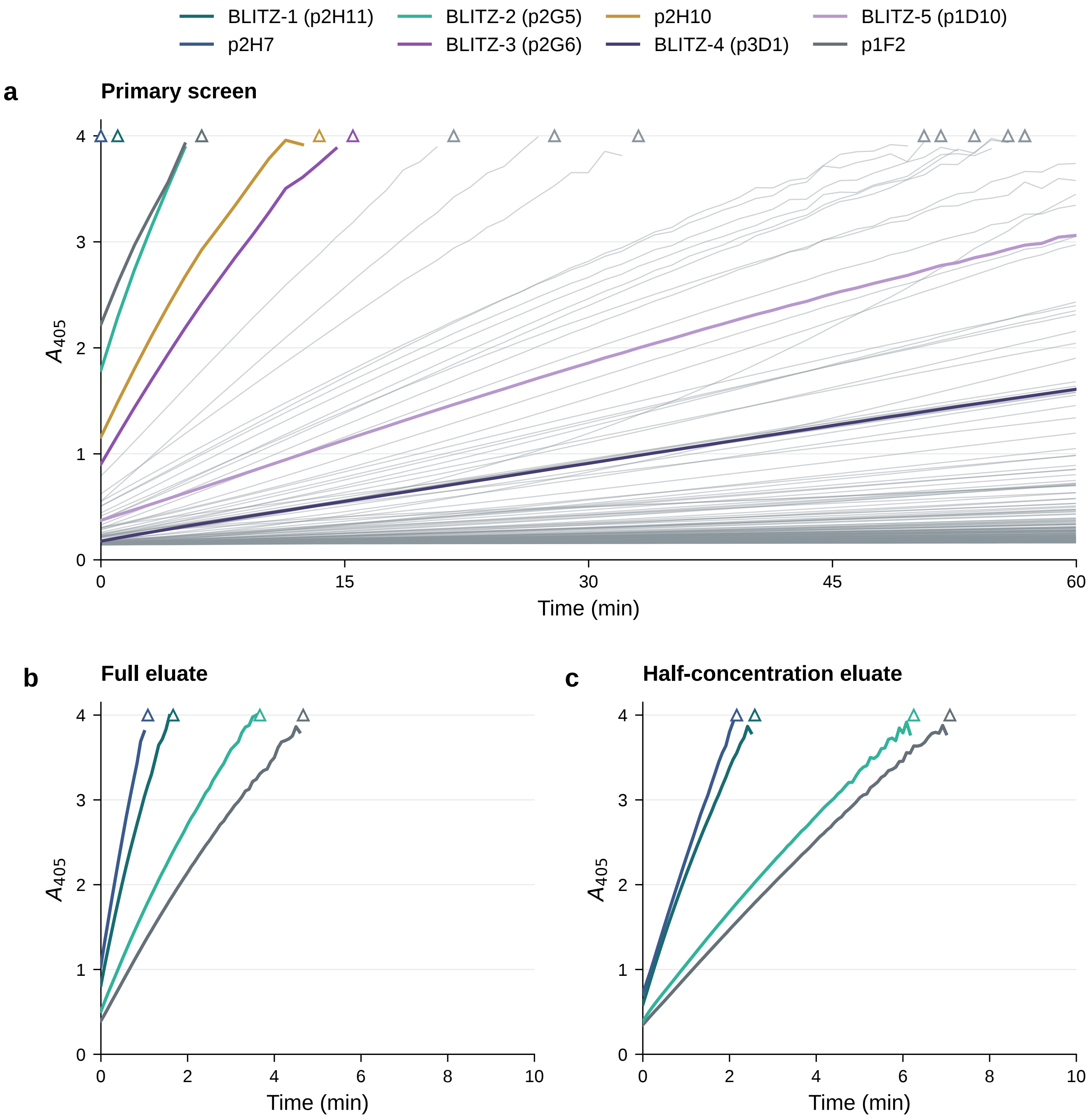

**Supplementary Fig. 30 PTE screening and rapid rescreening. a,** Eluate screen of 288 designs with 600 $\mu$M paraoxon and 1% methanol at 25 °C. Colored traces identify the eight designs taken forward for saturation kinetics; other designs are gray. **b,c,** Rapid rescreening of four designs with full or half-concentration eluate and 5-s reads. All panels show raw absorbance without smoothing or baseline subtraction. Time is measured from the first read. Traces stop at the first detector overflow, marked by an open triangle at 4 absorbance units.

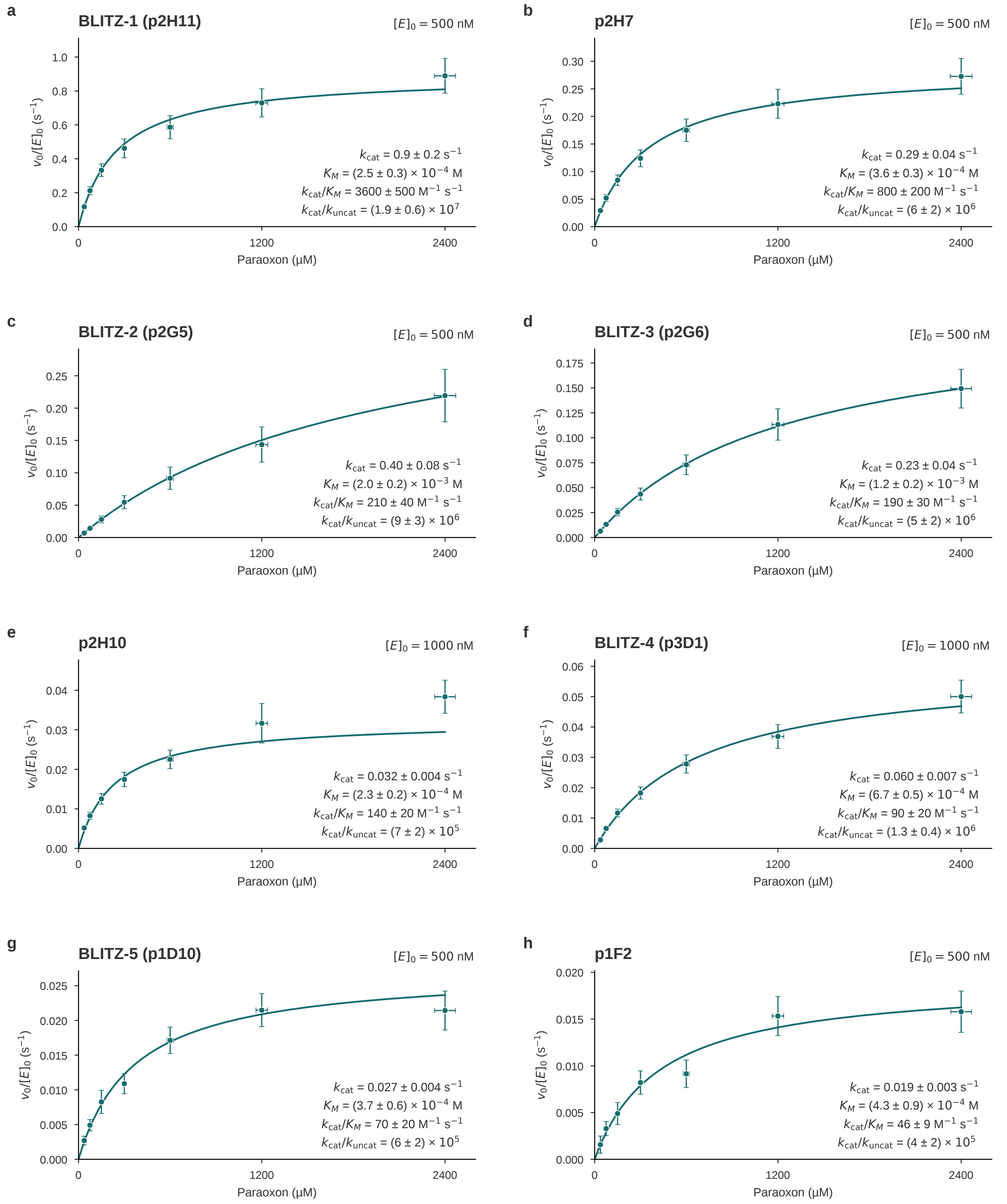

**Supplementary Fig. 31 Purified PTE saturation kinetics.** Selected assays for eight designs: BLITZ-1 (p2H11), p2H7, BLITZ-2 (p2G5), BLITZ-3 (p2G6), p2H10, BLITZ-4 (p3D1), BLITZ-5 (p1D10), and p1F2. Names rank the highest-efficiency characterized design from each scaffold cluster by mean $k_{cat}/K_M$. Enzyme concentrations are 500 nM except for p2H10 and BLITZ-4 (1 $\mu$M). Points show mean initial turnover rates from three technical replicates at each substrate concentration, with horizontal and vertical bars indicating propagated substrate and rate SD. Curves show nominal Michaelis–Menten fits, and parameter annotations give Monte Carlo means and propagated SD.

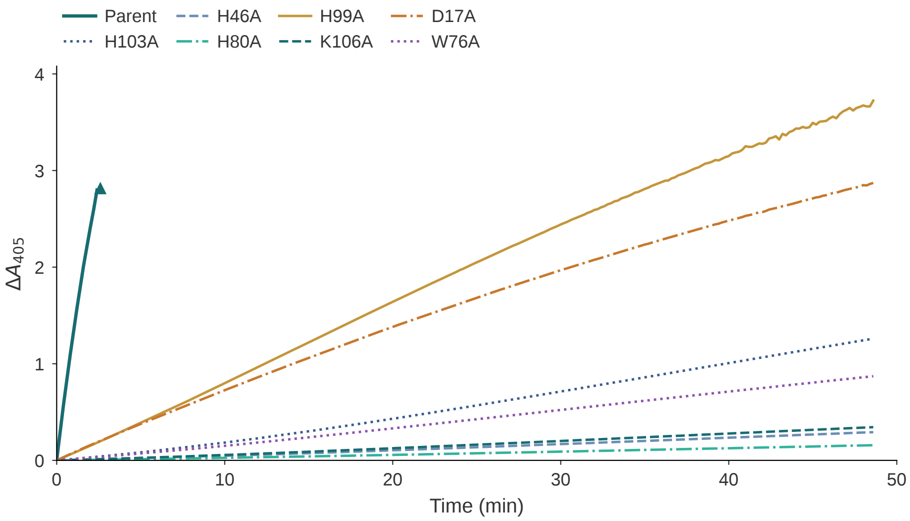

**Supplementary Fig. 32 Paraoxon-hydrolysis time courses for BLITZ-1 active-site variants.** Background-corrected $\Delta A_{405}$ traces for parent BLITZ-1 (p2H11) and alanine variants of seven first-shell catalytic residues. Triangles indicate detector overflow, at which point traces terminate. See Supplementary Section S13.7 for more details.

#### S14.8 Lenacapavir-precursor ligases

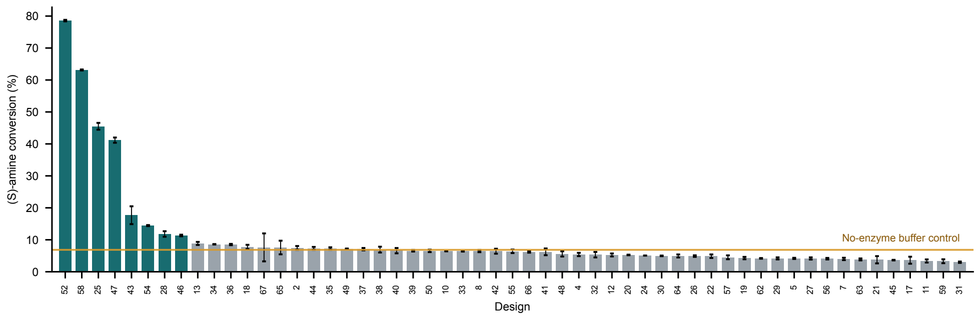

**Supplementary Fig. 33 Initial lenacapavir-precursor ligase screen.** ($S$)-amine conversion, amide/(amide + amine) from calibrated HPLC peak areas, for the designs with data in the initial screen (mean ± SD of three wells), ranked by conversion. The orange line and band mark the no-enzyme buffer control (6.8 ± 0.4%). Teal bars mark the eight designs above 10% conversion (52, 58, 25, 47, 43, 54, 28 and 46); the next design reaches 8.9%.

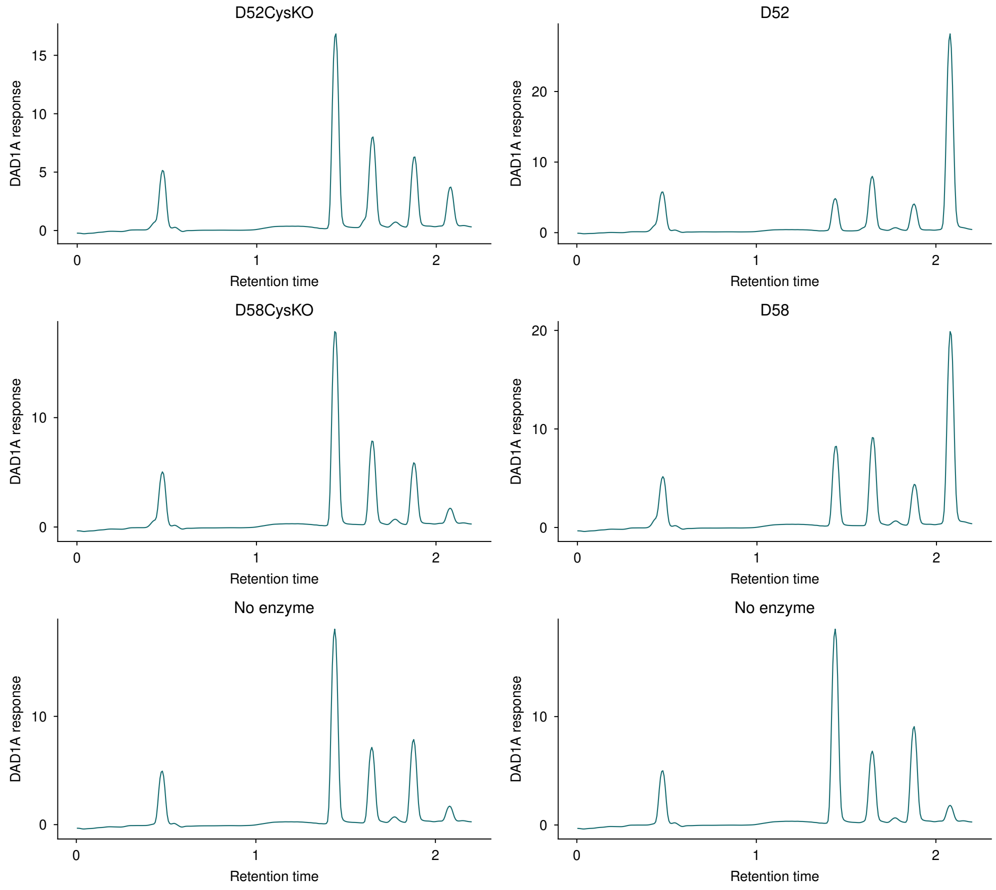

**Supplementary Fig. 34 Ligase parent, cysteine-knockout and no-enzyme chromatograms.** Six DAD1A chromatograms are shown without baseline correction, rescaling or peak assignment. The two no-enzyme controls are plotted separately.

### S15 Statistical analysis

#### S15.1 Autoencoder latent-space analysis

Canonical-parent retrieval in Supplementary Fig. 7 weights the 18 modified-amino-acid identities equally. Rankings use empirical Wasserstein distances averaged over 50 reference-sampling repetitions, with 100 residues per identity. Shuffled expectations preserve parent frequencies while permuting parent assignments across identities. Conditional 95% bootstrap intervals use 10,000 resamples of identities, with parent frequencies recomputed and the observed distance matrices held fixed.

#### S15.2 Adaptive reasoning and search for illustrative design case

For the feature-commitment analysis in Figure 1e, Spearman correlations are calculated across 500 trajectories. In Figure 1g, points show means ± one standard error of the mean (SEM) across design-level averages of two AF3 seeds. Each condition contained 5,000 planned paired replicates. Valid sample counts are 4,597–4,773 for motif RMSD and 4,992–4,998 for interface hydrogen-bond energy and min-iPAE. Missing measurements are not imputed. Supplementary Fig. 10 shows mean within-replicate changes from the shared one-pass baseline. Pairing and metric-specific inclusion criteria are described in Section S7.2.

##### S15.3 Cross-task benchmarks and binder–target similarity

For Figure 2, clustered pass rates divide the number of distinct passing clusters by 200 original attempts per target, including failures, then average targets equally. Target coverage is the fraction of targets with at least one passing design. GPU-normalized yields are calculated per target before averaging. For the compute-budget curves in Supplementary Fig. 16, shading shows the 10th–90th percentiles across 100 paired random orderings of attempts, sampled without replacement. Aggregation and reclustering are described in Section S8.

In Supplementary Fig. 13, PD-L1 fractions use Wilson 95% intervals within each length bin or conditioning group. Benchmark-wide frequencies average target-level fractions equally across 24 targets and use target-bootstrap intervals.

##### S15.4 Molecular-workflow evaluations

Figure 3a uses 114 PackBench targets and eight samples per method and setting. Quality and diversity are averaged within targets, then across targets; error bars show target-level standard errors.

Figure 3b uses 156 native backbones shared by both methods and 340 enzyme backbones (34 motifs × 10), with eight sequences each. Intervals for paired differences at matched diversity are 95% percentile intervals from 10,000 bootstrap resamples of backbones.

For Figure 3c, three AF3 seeds are averaged for each pairing, and intended-minus-off-target ipTM is calculated within each of 13 HBDesigner families. Families receive equal weight; intervals use 100,000 family-bootstrap resamples.

For Figure 3d, rates weight ligands equally after averaging within each ligand. Chiral-center adherence includes 33 eligible ligands, and E/Z-bond adherence includes nine. Error bars show 95% intervals from 100,000 paired ligand-bootstrap resamples. The AF3 reference includes all 430 RFD4-Proteina outputs.

For Figure 3e and Supplementary Fig. 22, AUROC intervals use 2,000 design-bootstrap resamples; paired method differences use 10,000 stratified design resamples, preserving active and inactive groups. Rescue fractions in Figure 3f use Wilson 95% intervals across 102 parent designs, with rescue defined by at least one passing result among six attempts. Sampling and scoring procedures for all six workflows are described in Section S10.

##### S15.5 Binder specificity

The calmodulin BLI curves in Figure 4a show the replicate-1 measurements.

For the pMHC titrations in Figure 4b, points and error bars show the mean and sample standard deviation (SD) of two plate-row observations per concentration. Zero-baseline Hill curves were fitted to all individual observations by unweighted least squares, with upper asymptotes constrained to 0–100%. The reported apparent $K_D$ values are the fitted half-maximal concentrations with avidity. IDR specificity responses in Figure 4d are normalized to peptide capture, with one measurement per binder–peptide pair. Kinetic fitting procedures are described in Sections S13.1, S13.3 and S13.4.

##### S15.6 Glycosylation and phosphopeptide recognition

The glycosylation boxplots in Figure 5c and Supplementary Fig. 26 summarize 97 RFDAA, 43 ΔRASA-filtered RFDAA and nine RFD4-Proteina profiles. Profiles below 5% occupancy and the F8 trimer are excluded. Boxes span the interquartile range, center lines show medians, and whiskers extend to observations within 1.5 interquartile ranges.

Phosphopeptide titrations in Figure 5d,e use one acquisition at each of 14 concentrations per peptide state. Four-parameter Hill curves were fitted to phosphorylated-peptide responses. For Figure 5f, NGS counts from one selected population per condition were normalized to sequencing depth, then to intended-target recovery (100%). Gating, fitting and sequence-count processing are described in Section S13.6.

#### S15.7 Enzyme kinetics and ligase conversion

PTE kinetic fits use three technical replicates per substrate concentration from one selected assay per design. Mean initial velocities were fitted by inverse-variance-weighted nonlinear regression to the Michaelis–Menten equation. Parameter uncertainties were propagated through 2,000 seeded Monte Carlo realizations with a joint refit in each realization, retaining shared concentration and calibration errors and parameter covariance. Parameter annotations report Monte Carlo means ± propagated SD; rate-plot bars show propagated standard uncertainties. The error model and fitting exclusions are described in Section S13.7.

PTE knockout rates for alanine variants of seven first-shell catalytic residues in Figure 6g are ordinary least-squares slopes of reference-corrected $A_{405}$ over the common 0–60 s interval after the first recorded read (six observations), expressed as percentages of the parent slope. The corresponding time courses in Supplementary Fig. 32 are additionally shifted by each trace's first value and stop at their original overflow boundaries.

Ligase scores in Figure 6m are the difference between each design's mean conversion and its campaign's mean background, divided by the campaign background sample SD. The RFD3 lysate screen includes 71 designs with three measurements each; the RFD4-Proteina purified-protein screen includes 55 designs, 49 with three measurements and six with six measurements. The respective background means and sample SDs are 34.21 ± 30.95% and 6.30 ± 1.02%.
